# Base-resolution models of transcription factor binding reveal soft motif syntax

**DOI:** 10.1101/737981

**Authors:** Žiga Avsec, Melanie Weilert, Avanti Shrikumar, Sabrina Krueger, Amr Alexandari, Khyati Dalal, Robin Fropf, Charles McAnany, Julien Gagneur, Anshul Kundaje, Julia Zeitlinger

## Abstract

The arrangement of transcription factor (TF) binding motifs (syntax) is an important part of the cis-regulatory code, yet remains elusive. We introduce a deep learning model, BPNet, that uses DNA sequence to predict base-resolution ChIP-nexus binding profiles of pluripotency TFs. We develop interpretation tools to learn predictive motif representations and identify soft syntax rules for cooperative TF binding interactions. Strikingly, Nanog preferentially binds with helical periodicity, and TFs often cooperate in a directional manner, which we validate using CRISPR-induced point mutations. Our model represents a powerful general approach to uncover the motifs and syntax of cis-regulatory sequences in genomics data.

**Highlights:**

- The neural network BPNet accurately predicts TF binding data at base-resolution.
- Model interpretation discovers TF motifs and TF interactions dependent on soft syntax.
- Motifs for Nanog and partners are preferentially spaced at ∼10.5 bp periodicity.
- Directional cooperativity is validated: Sox2 enhances Nanog binding, but not vice versa.

## Introduction

Understanding the cis-regulatory code of the genome is vital for understanding when and where genes are expressed and how genetic variation and somatic mutations affect disease. Despite extensive molecular efforts to map millions of putative enhancers in a wide variety of cell types and tissues^1–3^, identifying the critical bases that alter their regulatory information remains a major challenge. It is known that short sequence motifs are critical for the binding of sequence-specific transcription factors (TFs), but whether motifs are bound *in vivo* may depend on the combination and syntactic arrangements of other motifs nearby. For example, two or more strictly spaced motifs may form composite motifs that provide a platform for DNA-mediated cooperativity between the corresponding TFs^4^. However, whether less strict (“soft”) motif spacing preferences exist and influence TF cooperativity is not clear. The precise rules by which combinations of motifs and their syntax form the cis-regulatory code remain to be elucidated.

Experimental manipulations of enhancer sequences, such as mutations or synthetic designs, strongly support the existence of subtle motif syntax^e.g. 5–12^. However, genome-wide analyses have rarely identified statistically over-represented motif syntax rules, questioning whether they exist and impose evolutionarily constraints on enhancer function^13–17^. When patterns are discovered computationally^18–24^, they are difficult to validate experimentally and the mechanism by which they might affect TF cooperativity is not clear. For example, over-represented instances of strict motif spacings are sometimes associated with retrotransposons that contain multiple TF binding motifs^19,20^. Thus, the appearance of syntax may be the result of biases inherent to genome composition, rather than functional constraints on enhancer function.

There is hence a critical need for a general method that can identify cis-regulatory motif syntax based on genome-wide experimental data. Recently, convolutional neural networks (CNNs) have been applied towards accurately predicting diverse molecular phenotypes including TF binding from DNA sequence^25–28^. The advantage of CNNs is that they can learn arbitrarily complex *de novo* patterns from raw sequences in the form of flexible predictive models composed of hierarchical layers of non-linear transformations, allowing them to capture sequence motifs and their higher-order organizational context without making strong prior biological assumptions. However, the complexity of these models makes them particularly challenging to interpret. While several *ad hoc* methods have been developed to visualize TF binding motifs from trained CNNs^25,26,28–32^, methods for extracting the rules by which motif syntax informs TF binding are lacking^33^.

Another critical limitation of current methods is resolution. The standard approach for mapping motif instances bound by TFs *in vivo* is to extract bound regions from chromatin immunoprecipitation experiments coupled to sequencing (ChIP-seq) using peak-callers^34–39^ and identify over-represented motifs in these sequences as position weight matrix models (PWM)^40–43^. While CNNs are ideally suited to model TF binding from motif combinations and their syntax, current models have limited resolution. State-of-the-art CNN models of TF binding predict binary binding events^25–27^ or low-resolution continuous binding signal averaged across 100-200 bp windows^44^.

There is evidence that TF cooperativity can be detected in ChIP-seq experiments. For example, TFs sometimes bind indirectly to motifs of other TFs^16,20,45–47^. TF cooperativity is even more apparent when the resolution of ChIP-seq is improved by adding an exonuclease digestion step (ChIP-exo)^48^. ChIP-exo methods such as ChIP-nexus generate base-resolution footprints precisely over the motif instances bound by the TF *in vivo*^*49,50*^ and these footprints differ between directly and indirectly bound motifs^50,51^. ChIP-nexus profiles have also provided evidence that TFs may help the binding of another TF nearby^52^. Although the full extent of TF cooperativity at the level of binding is not known, these results indicate that ChIP-seq data, and especially ChIP-nexus data, are a useful readout for cis-regulatory motif syntax, if the data are modeled at sufficiently high resolution.

To discover motif syntax, we developed a novel CNN called BPNet that models the relationship between cis-regulatory sequence and TF binding profiles at base-resolution. To use a biological system with extensive prior knowledge, we studied the pluripotency TFs Oct4, Sox2, Nanog and Klf4 in mouse embryonic stem cells (ESCs) and generated ChIP-nexus data for maximal resolution. We trained base-resolution BPNet models on these ChIP-nexus profiles with high predictive performance, on par with the concordance between replicate experiments. We substantially expanded model interpretation methods to extract new motif representations that summarize their predictive influence on TF binding instead of statistically over-represented base frequencies. We then developed new methods that use the trained BPNet model as an *in silico* oracle to derive rules of TF cooperativity that depend on motif syntax.

We find that strict motif spacings in the genome are mainly due to retrotransposons, but that TF cooperativity depends on preferential soft motif syntax that is in agreement with experimentally characterized protein-protein or nucleosome interactions in ESCs. We also observed unexpected rules of TF binding cooperativity, including a broad preference for Nanog to bind DNA with helical periodicity, and validate some of these results experimentally. These results suggest that end-to-end neural network models trained on high-resolution genomics data, coupled with a dedicated suite of interpretation tools, can serve as a powerful tool for discovering the critical bases within cis-regulatory sequences, obtaining predictive motif representations and identifying the underlying motif syntax associated with TF cooperativity.

## Results

### BPNet predicts base-resolution TF binding profiles from DNA sequence

We performed ChIP-nexus experiments for Oct4, Sox2, Nanog and Klf4 in mouse ESCs and obtained genome-wide strand-specific base-resolution profiles for each TF (Figure 1A). As we have shown for previous TF ChIP-nexus data^49^, the profiles on known TF binding motifs showed consistent stereotypical footprints across various genomic regions, as illustrated by the binding of Oct4 and Sox2 to the composite *Oct4-Sox2* motif^53^ (Figure 1B). These footprints not only had higher resolution compared to ChIP-seq data, but also displayed increased motif specificity. For example, the *Sox2* motif showed a sharp ChIP-nexus footprint for Sox2 but not for Oct4, while ChIP-seq data showed binding signal for both (Figure 1C).

**Figure 1.**
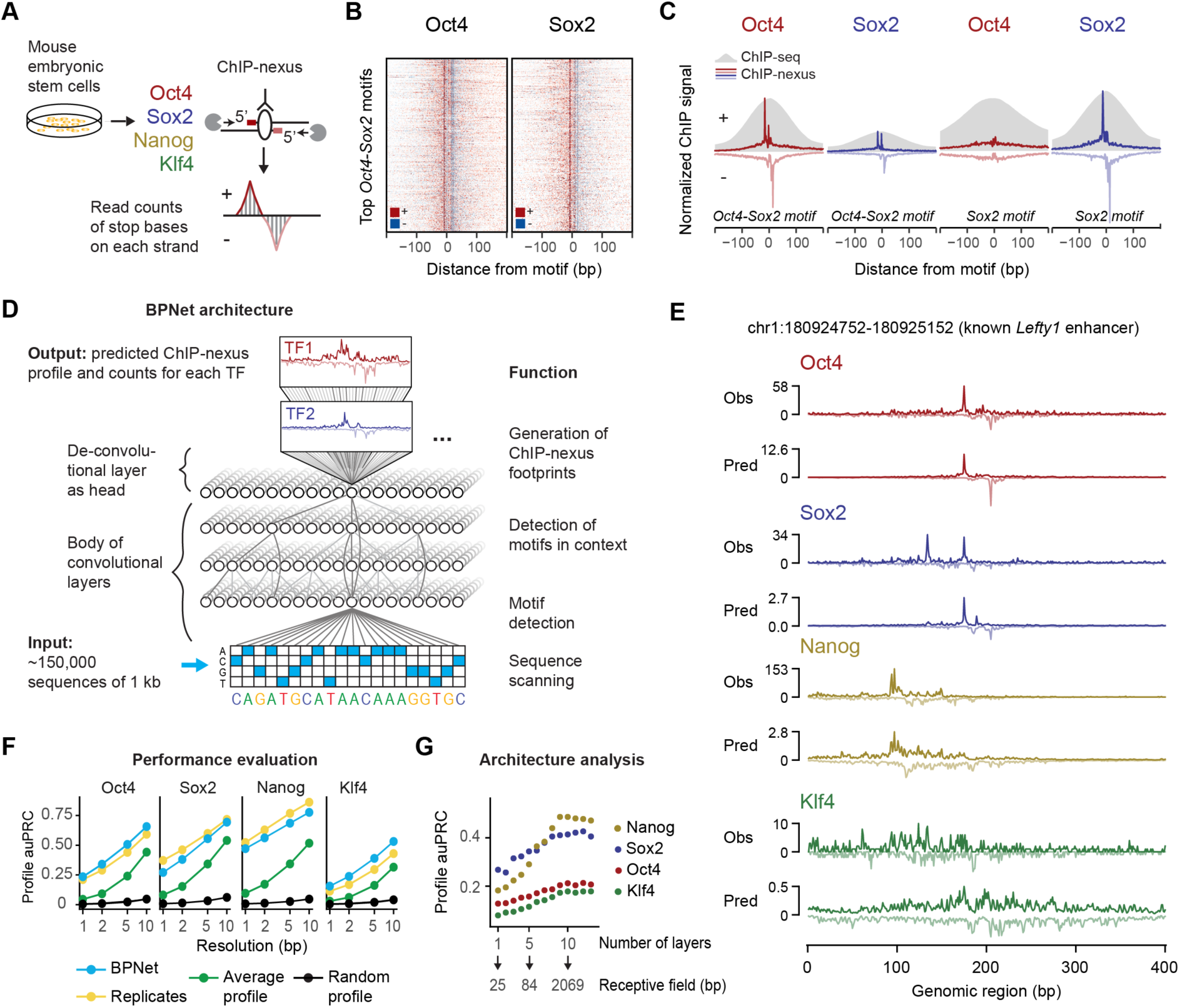
BPNet predicts ChIP-nexus signal at base-resolution. **A)** ChIP-nexus experiments were performed on Oct4, Sox2, Nanog, and Klf4 in mouse embryonic stem cells. After digestion of the 5’ DNA ends with lambda exonuclease, strand-specific stop sites were mapped to the genome at base-resolution. Bound sites exhibit a distinct footprint of aligned reads, where the positive (+) strand peak occurs many bases before the negative (-) strand peak. **B)** Profile heatmaps of Oct4 and Sox2 ChIP-nexus data (red and blue for positive and negative strand respectively, color depth represents normalized signal intensity) at the top 500 *Oct4-Sox2* motif sites. **C)** The average ChIP-nexus data (positive and negative strand at top and bottom) and ChIP-seq data (grey) at the top 500 *Oct4-Sox2* and *Sox2* motif sites for Oct4 (red) and Sox2 (blue). The ChIP-nexus data have higher resolution and show less unspecific binding of Oct4 to the *Sox2* motif. **D)** Architecture of the convolutional neural network (BPNet) that was trained to simultaneously predict the number of ChIP-nexus reads at each strand for all TFs from 1 kb DNA sequences. **E)** Observed and predicted ChIP-nexus read coverage of the positive strand (top) and the negative strand (bottom) for the *Lefty* enhancer located on the held-out test chromosome 8. **F)** BPNet predicts the positions of local high signal in the profiles at replicate-level accuracy as measured by the area under precision-recall curve (auPRC) at resolutions from 1 bp to 10 bp in held-out test chromosomes 1, 8 and 9. **G)** More convolutional layers (x-axis) increase the number of input bases considered for profile prediction at each position (receptive field) and this yields increasingly more accurate profile shape predictions on the tuning chromosomes 2-4 (measured in auPRC as above).

Since enhancers typically contain autonomous sequences of <500 bp that can function outside its genomic context^54,55^, we trained BPNet on sequences of 1 kb. We identified 147,974 genomic regions exhibiting statistically significant and reproducible enrichment of ChIP-nexus signal for Oct4, Sox2, Nanog or Klf4 and trained BPNet to predict the strand-specific ChIP-nexus profiles of all four TFs. Each profile can be characterized by two coupled properties - the total signal (read counts) and the shape of the profile (base-resolution distribution of reads). We decided to explicitly model both since the profile shape is likely primarily determined by the local sequence context, while the total signal could be influenced by factors that we do not model, including chromatin state and higher-order chromatin organization.

To accurately learn the relationship between DNA sequence and base-resolution binding profiles (Figure 1D), we designed BPNet as a CNN with the following properties. (1) It consists of multiple dilated convolutional layers^44,56^ with residual skip connections^57,58^ to learn increasingly complex predictive sequence patterns across the 1 kb sequence. (2) It uses multi-task learning to jointly train on the ChIP-nexus profiles of all four TFs. (3) It uses experimental control data (PAtCh-CAP^59^) as an auxiliary input to prevent the model from learning sequence features associated with potential experimental biases in the data (Methods). (4) It uses a multi-scale loss function to separately optimize the predictions of ChIP-nexus profile shape and total read counts. The training of the BPNet model, tuning its parameters and testing of the prediction performance were performed on different sets of genomic regions (found on distinct chromosomes). We also confirmed that mappability of regions did not bias the predictions (Figure S1).

To evaluate the predictive performance, we first inspected individual enhancers located on held-out test chromosomes such as those associated with the genes *Lefty1*^*60*^, *Zfp281*^*61*^, and *Sall1*^*62,63*^ and found that the predicted and observed ChIP-nexus profiles were noticeably similar, with highly concordant summits of footprints (Figure 1E and Figure S2A). We then systematically compared the positions of high ChIP-nexus counts between predicted versus observed profiles in all regions of the held-out test set. Strikingly, the positional concordance was on par with replicate experiments and substantially better than randomized profiles or average profiles at resolutions ranging from 1-10 bp (Figure 1F). These results show that BPNet accurately learned to predict the ChIP-nexus binding profiles of all four TFs from DNA sequence.

To identify key components for the high prediction performance, we systematically varied the network architecture (Figure S2B,C). We found that the large number of convolutional layers was critical for predicting all four ChIP-nexus data sets and was particularly important for Nanog. This indicates that the learned sequence patterns required to predict ChIP-nexus profiles span over larger sequence regions beyond individual motifs^64^, especially in the case of Nanog (Figure 1G). We also found that prioritizing profile versus total count prediction during training affected the prediction performance. While up-weighting the profile predictions improved the performance of the profile predictions, the performance for the total counts (R_s_ = 0.62) did not reach replicate-level performance (R_s_=0.94), irrespective of up-weighting (Figure S2D,E). These results are consistent with our assumption that longer sequences or other measurements such as local chromatin state may be required to better predict total TF occupancy^64^, but that local sequence context (1 kb) is well suited to accurately predict the shape of ChIP-nexus profiles.

### A suite of model interpretation tools to identify TF binding motifs and genomic motif instances

We next set out to extract the sequence features that were predictive of TF binding from the trained BPNet model. We extended our previously developed tool DeepLIFT^65^ to quantify the contribution of each base within an input sequence to the entire predicted ChIP-nexus profile of a TF on both strands (Figure 2A, Methods). For every input sequence, separate base-resolution contribution scores were inferred for each of the four TFs.

**Figure 2.**
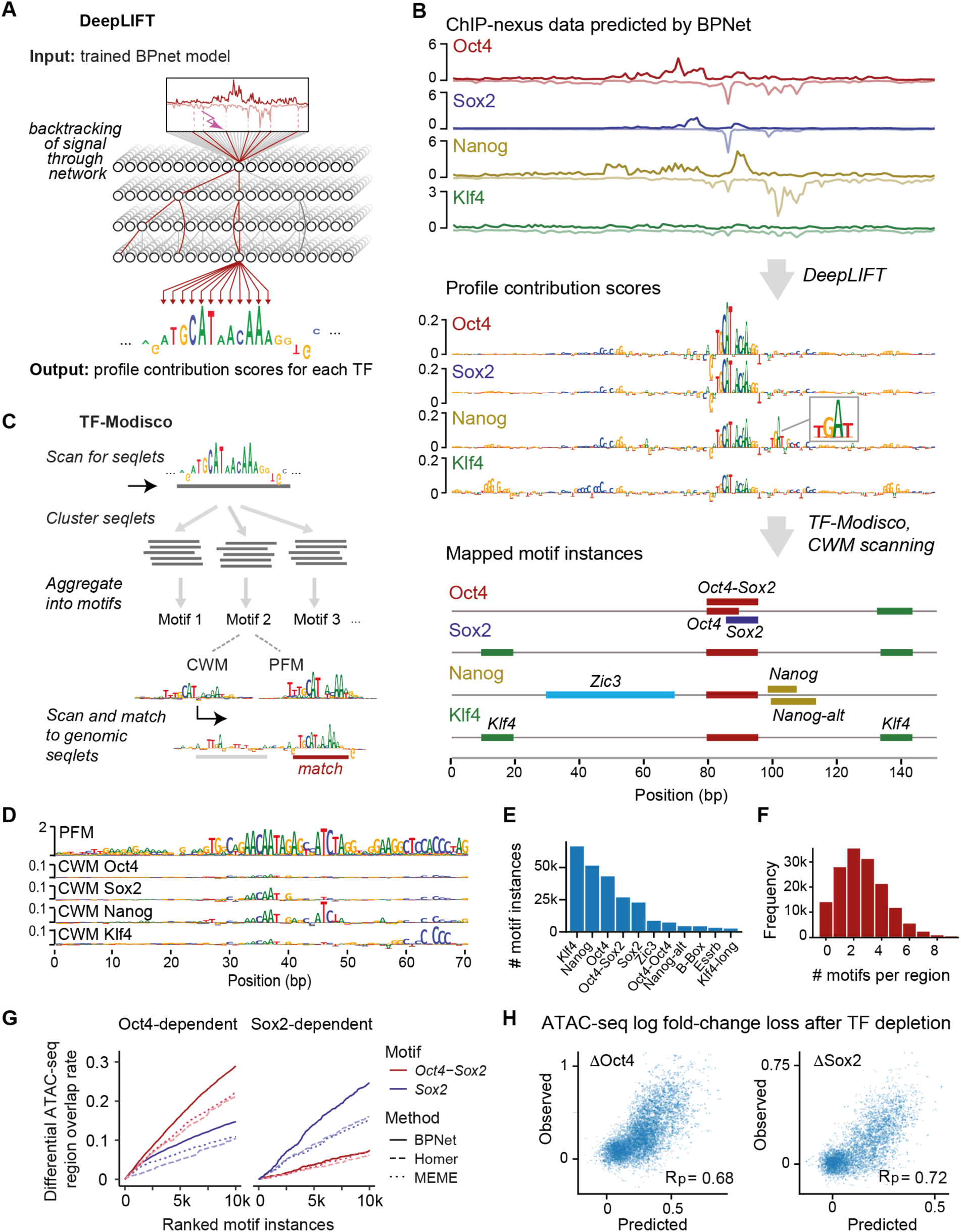
Transcription factor binding motifs can be accurately derived from BPNet and mapped to the genome using interpretation tools. **A)** DeepLIFT recursively decomposes the predicted TF-specific binding output of the model and quantifies the contribution of each base of the input DNA sequence by backtracking the prediction through the network. **B)** Procedure for inferring and mapping predictive motif instances using the known distal *Oct4* enhancer (chr17:35504453-35504603) as an example. From the predicted ChIP-nexus profile for each TF (top), DeepLIFT derives TF-specific profile contribution scores (middle). Regions with high contribution scores (called seqlets) resemble TF binding motifs. Seqlets are annotated by scanning the contribution scores with motifs discovered by TF-MoDISco (bottom). **C)** To discover motifs, TF-MoDISco scans for seqlets, extends the seqlets to 70 bp, performs pairwise alignments and clusters the seqlets. For each cluster, a motif is derived as contribution weight matrix (CWM), obtained by averaging the contribution scores of each of the 4 bases at each position across all aligned seqlets. The corresponding position frequency matrix (PFM) is the frequency of bases at each position. Motif instances are identified by scanning the CWM for each motif for high scoring matches across the profile contribution scores in the genomic regions. **D)** Example of a motif (N6) where the PFM differs from the CWMs. The PFM indicates that it is a repeat sequence (*RLTR9E*), while the CWM for each TF highlights the sequences that contribute to binding. **E)** Number of motif instances in thousands (k) found in the ∼150,000 genomic regions for the 11 representative motifs. **F)** Histogram of the number mapped motif instances in thousands (k) found per region. **G)** Evaluation of the mapped motifs using independent data: regions that lose ATAC-seq signal in response to either Oct4 or Sox2 depletion (but not both) as defined by Friman et al., 2019. BPNet motif instances of *Oct4-Sox2* and *Sox2* (ranked by contribution scores) outperformed those obtained by HOMER and MEME (ranked by PWM match scores). **H)** A linear model based on the bottleneck layer of the trained BPNet model makes accurate quantitative predictions of the log fold-change loss in ATAC-seq signal upon depletion of Oct4 (*Δ*Oct4) or Sox2 (*Δ*Sox2). Results are shown with Pearson correlation coefficient (R_p_) for the test chromosomes 1, 8, and 9 that were held out during training. See Figure S12B for a similar model based on motif instance features.

The TF-specific contribution scores are illustrated at the distal *Oct4* enhancer where all four TFs show strong predicted footprints matching the observed ChIP-nexus footprints (Figure 2B top, Figure S3A). The contribution scores are particularly high at specific subsequences, which we call seqlets and which resemble TF binding motifs (Figure 2B middle). One prominent seqlet matches the composite *Oct4-Sox2* motif, which has previously been mapped to this exact position in the *Oct4* enhancer^66^. This motif has high contribution scores for Oct4 and Sox2, which are directly bound to the motif, and for Nanog and Klf4 at slightly lower levels (Figure 2B middle). This suggests that the *Oct4-Sox2* motif is indirectly important for the binding of other TFs.

Other seqlets did not readily match known motifs. For example, we found a TGAT sequence in the middle of the Nanog footprint (highlighted in Figure 2A middle), but it was unclear whether it is a *Nanog* motif since previous reports on its consensus have been conflicting^47,67–72^. These results demonstrate the ability of contribution scores to highlight TF binding motifs, but indicate the need to identify and characterize the motifs more systematically.

To systematically discover and summarize recurring predictive sequence patterns, we used TF-MoDISco, which uses sequences and their associated base-resolution contribution scores as input^31^. For each TF, TF-MoDISCo identifies seqlets from the contribution scores in all regions and then optimally aligns and clusters them into motifs (Figure 2C). For each cluster, a novel motif representation called contribution weight matrix (CWM) is derived by averaging the contribution scores of each of the four possible bases at every position across the seqlets. A more traditional position frequency matrix (PFM) representation, which contains the normalized base frequencies instead of the average contribution scores, is also calculated (see Supplemental Q&A on CWMs and PFMs/PWMs).

TF-MoDISco discovered 51 motifs, but 18 of them had unusually long PFMs (> 40 bp) with high information content (30-100 bits) (Figure S4A, example in Figure 2D). This implies that the genomic instances of these motifs share near identical base composition across the entire length of the pattern (despite being discovered by uniquely mappable ChIP-nexus reads). Indeed, we found that the majority of them (>80%) overlapped with annotated repeat elements (Figure S4B). The most common were long-terminal repeats (LTRs) of endogenous retrotransposon viruses (ERVs), including those of the ERVK, ERVL and the ERVL-MaLR family (Figure S4C). Remarkably, the corresponding CWM representations of these long PFMs were quite different. Instead of long stretches of uniformly over-represented bases, the CWMs highlighted the shorter subsequences *predictive* of TF binding (Figure 2D, Figure S4C). This difference between CWM and PFM representations provides a means to discover and pinpoint bound motifs within retrotransposons.

The remaining 33 motifs were all interpretable TF binding motifs, but contained subsets with subtle differences, leading us to select 11 representative motifs for further analysis (Figure S5). These motifs include the well-known *Oct4-Sox2, Sox2*, and *Klf4* motifs, as well as the *Zic3* and *Esrrb* motifs, which bind pluripotency TFs that we did not profile (Figure S5 and below). All motifs were overall robustly discovered by TF-MoDISCo from five different BPNet models trained on different subsets of ChIP-nexus peak regions (Figure S6).

Using the 11 representative motifs, we then comprehensively mapped and labeled all predictive motif instances in the bound genomic regions. We scanned the base-resolution contribution scores of all regions and annotated predictive motif instances that had high contribution scores and high match scores to the CWM (Figure 2C and Figure S7). In total, we obtained 241,005 unique motif instances in the 147,974 genomic regions, with *Klf4* motifs occurring most frequently (Figure 2E). Altogether, 72,696 regions (48.1%) have at least three motif instances and 20,352 regions (13.5%) have at least 5 motif instances (Figure 2F). These genome-wide motif annotations are in agreement with motif instances supported by previous independent validation experiments^73–75^ (Figure S3B,C,D) and provide a strong foundation for analyzing genome-wide motif syntax and characterizing known functional enhancers is mouse ESCs (Figure 2B bottom, Figure S8).

The motif maps derived from BPNet outperformed those obtained by traditional approaches such as PWM scanning (Supplemental Text, Figure S9). BPNet discovered motif instances more reliably than MEME^40–43^ or HOMER^76^ and correctly identified more motif instances supported by ChIP-nexus data on held-out test chromosomes from raw sequence, especially for the short *Nanog* motif (Supplemental Text, Figure S9). Unlike PWM-based motif scanning methods that compute match scores only using raw sequence, BPNet’s CWM scanning method also incorporates the predictive contribution scores of bases, which are derived from the BPNet model accounting for the entire 1 Kb sequence context. Thus, the BPNet derived motif instances take favorable sequence contexts into account, which reduces the false discovery rate in unfavorable regions. We also found that training the model on base-resolution profiles, instead of course-resolution binary (bound vs. unbound) labels was necessary for the BPNet’s CWM scanning approach to outperform PWM scanning (Supplemental Text, Figure S10, S11).

Our method also outperformed traditional methods when using an independent, previously published ATAC-seq data set^77^ for the evaluation. Regions with differential chromatin accessibility after induced depletion of Oct4 or Sox2 in mESCs (as defined by the authors) overlapped more highly with *Oct4-Sox2* and *Sox2* motif instances ranked by BPNet contribution scores, compared to motif instances ranked by motif scores from MEME or HOMER (Figure 2G, Figure S12A). These results support the high quality and biological relevance of the BPNet mapped motif instances relative to those obtained from traditional motif discovery and scanning methods.

Finally, we found that sequence features derived from the BPNet TF binding models can also accurately predict quantitative changes in ATAC-seq signal after Oct4 and Sox2 depletion. Specifically, linear models trained using the sequence features encoded in the final convolutional layer of the BPNet model were able to accurately predict differential accessibility (Figure 2H). These models outperformed linear models trained using only the inferred motif instances (Figure S12B). These results indicate that the complete sequence representation learned by BPNet encodes predictive features beyond linear, additive effects of the motif instances. Hence, we set out to identify higher-order sequence features such as motif syntax.

### Discovery of composite motifs and indirect binding footprints

As a first step to identifying motif syntax, we inspected the motifs identified by TF-MoDISCo for composite motifs, the simplest form of motif syntax. Indeed, we not only discovered the *Sox2* motif and the monomeric *Oct4* motif^81^, but also the composite *Oct4-Oct4* motif (Figure 3A), a near-palindromic motif that resembles the MORE and PORE motifs bound by Oct4 homodimers^82,83^. This motif has not previously been shown to be bound in ESCs *in vivo*, but is known to be important during neuronal differentiation^84^. Finally, we rediscovered the *Oct4-Sox2* motif, in which the bases with high contribution scores correspond to the specific DNA contacts made by the heterodimer (based on the Oct1-Sox2 crystal structure)^53,85,86^ (Figure 3A right). Thus, we discovered composite motifs that are consistent with known structural data.

**Figure 3.**
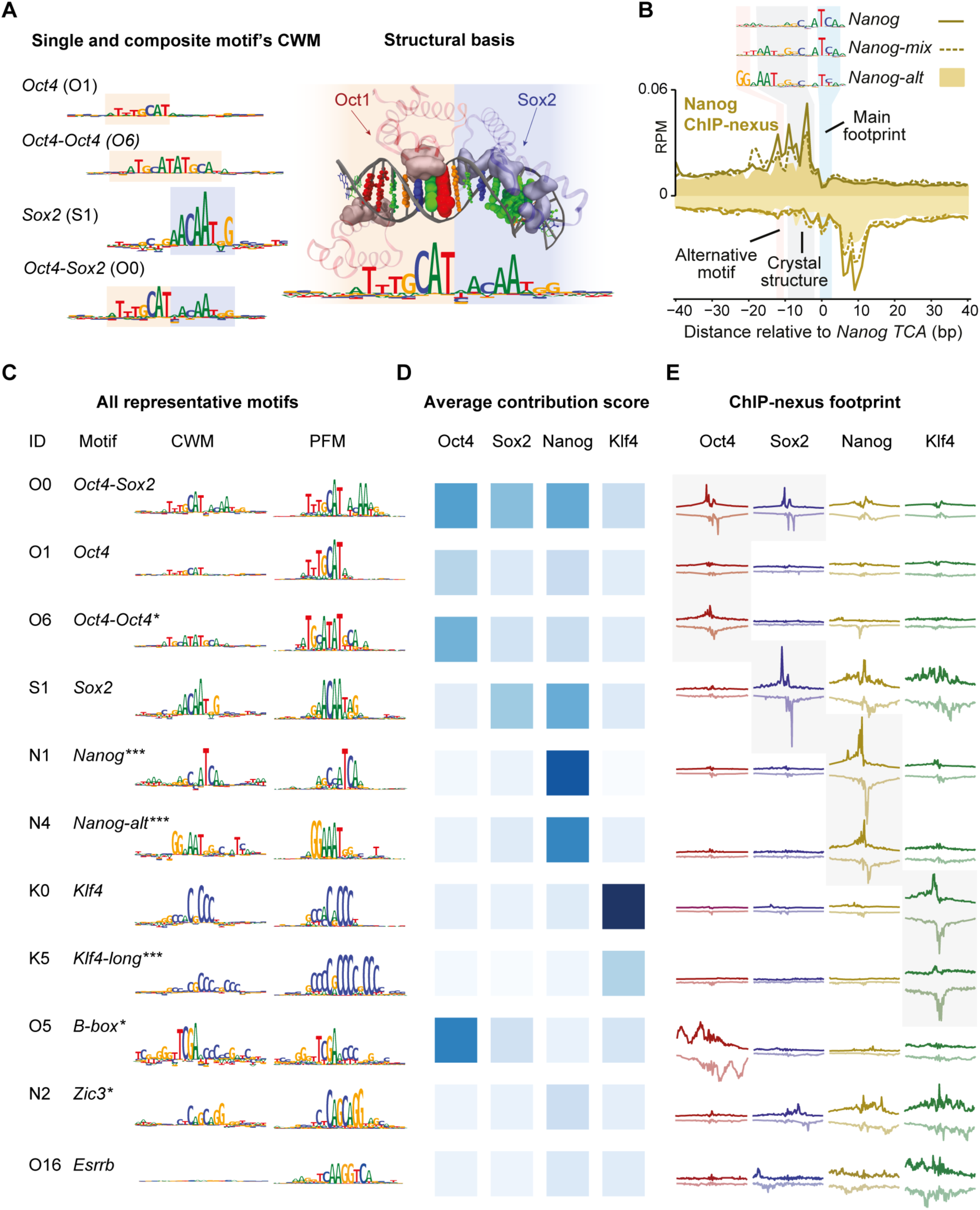
Discovery of composite motifs and indirect binding footprints. **A)** The *Oct4, Oct4-Oct4, Sox2* and *Oct4-Sox2* (left) were identified by TF-MoDISco as separate motifs, highlighting its ability to identify composite motifs. The CWM of the *Oct4-Sox2* composite motif correlates with the structure of Oct1 and Sox2 bound to the *Oct4-Sox2* motif (right). For visualization, the amino acids of Oct1 and Sox2 that contact DNA are shown as solid, and the atoms in the DNA bases, shown as colored spheres, are sized according to the contribution scores shown in the CWM below (right). **B)** Nanog ChIP-nexus binding footprints were associated with three Nanog motif variants (shown as CWM). For all motifs, the strongest footprint was found at the TCA sequence (blue). The CWM of *Nanog-mix* (N5) and *Nanog-alt* (N4) contain a sequence that matches the sequence AATGGGC bound by Nanog in a crystal structure (grey)^78^. The CWM of *Nanog-alt* contains GG (pink). **C)** The discovered representative short motifs contain known motifs, new motifs (***), and known motifs new in this context (*). From left to right: motif ID, motif name, CWM, PFM. All sequence logos share the same y-axis. The *B-box* mediates RNA polymerase III transcription^79,80^ and is associated with high levels of Oct4 binding upstream and downstream of tRNA (Figure S15C,D). **D)** The average contribution score of the motif for each TF is shown as color from white to dark blue. The highest score may indicate the TF that binds directly. **E)** The TF’s average ChIP-nexus footprint (read count distribution on the positive strand at the top and negative strand at the bottom in a 200-bp window) better indicates whether the motif is directly bound (sharp profile, marked with grey background), indirectly bound (fuzzy profile) or not bound at all. The footprints for each TF share the same y-axis.

We did not identify the less well-studied composite *Sox2-Nanog* motif^71^ and found no evidence that this motif was bound in our ChIP-nexus data (Figure S13A). Instead, we identified three *Nanog* motifs: *Nanog, Nanog-alt* and *Nanog-mix*, the latter of which is partially redundant with the first two. All have a main footprint around a TCA core sequence (Figure 3B). *Nanog* resembles the *Nanog* motif identified previously by a thermodynamic model from ChIP-seq data^72^. Consistent with direct binding, a closely matching sequence (GCCATCA) is bound by Nanog in an EMSA gel shift assay^72^. *Nanog-alt* and *Nanog-mix* contain the sequence to which monomeric Nanog is bound in a crystal structure (AATGGGC)^78^. Given these two separate direct DNA contacts, the observed Nanog binding footprint likely represents Nanog binding as a homodimer^87^. But since *Nanog-alt* contains an additional GG to the left (Figure 3B), we cannot rule out the existence of an unknown Nanog binding partner (but it is not Sox2 or Pbx, see Figure S13B,C).

The majority of composite motifs, however, came from retrotransposons. This is consistent with previous observations that retrotransposons may contain multiple ancestral TF binding sites^88–92^ (Figure S14A). Among all motif pairs, the top 1% most frequent distances mapped in 83% to ERVs and were often larger than 20 bp (Figure S14B,C), which exceeds the typical distance between motifs found in composite motifs that promote TF cooperativity^93,94^. This suggests that over-represented strict motif spacings alone are not a reliable indicator for functional motif syntax.

We next analyzed whether the 11 motifs showed evidence for mediating cooperative TF interactions in the absence of strict motif spacings (Figure 3C). By inspecting the contribution scores (Figure 3D), we found that many motifs were predicted to contribute to the binding of other TFs. Moreover, we discovered motifs of pluripotency TFs that we did not profile, including the *Zic3* and *Esrrb* motifs, which we validated with additional ChIP-nexus experiments (Figure S15). Thus, BPNet predicts that Oct4, Sox2, Nanog, and Klf4 frequently bind with the help of motifs from other TFs.

One explanation for this observation is that TFs may be indirectly recruited to motifs of other TFs^50,51^. We therefore inspected the average ChIP-nexus binding footprints of all TFs at all motifs (Figure 3E). We found that directly bound TFs have very sharp average ChIP-nexus footprints (marked in grey in Figure 3E), but that TFs also showed broader, more fuzzy footprints, which we attribute to indirect binding. The level of indirect TF occupancy correlated with the contribution score for the TF (Figure 3D,E), suggesting that the indirect footprints are predicted by BPNet.

Notably, the indirect footprints tended to occur in an asymmetric or directional manner (Figure 3D,E). For example, Nanog was bound indirectly to the *Sox2* motif, but Sox2 was not detected on the *Nanog* motif. Since Sox2 and Nanog have been shown to physically interact with each other^71,95^, this suggests that these TFs indeed cooperate in some way, but not through a composite motif. We therefore set out to analyze more systematically how motif pairs influence cooperative binding, which would represent a means to identifying functional motif syntax.

### Using BPNet as an *in silico* oracle reveals cooperative TF interactions

To systematically extract rules of TF cooperativity, we developed two approaches that interrogate the trained BPNet model *in silico* like an oracle. In both approaches, we measured how the binding of a TF to its motif is enhanced by a second motif dependent on the distance between them (Figure 4A,B, Figure S16). We focused on the motifs most strongly bound by each of the four TFs: *Oct4-Sox2* (bound by Oct4), *Sox2, Nanog*, and *Klf4*. The first approach uses synthetically designed sequences (Figure 4A), while the second uses naturally occurring non-overlapping motifs in genomic sequences with and without perturbations (Figure 4B).

**Figure 4.**
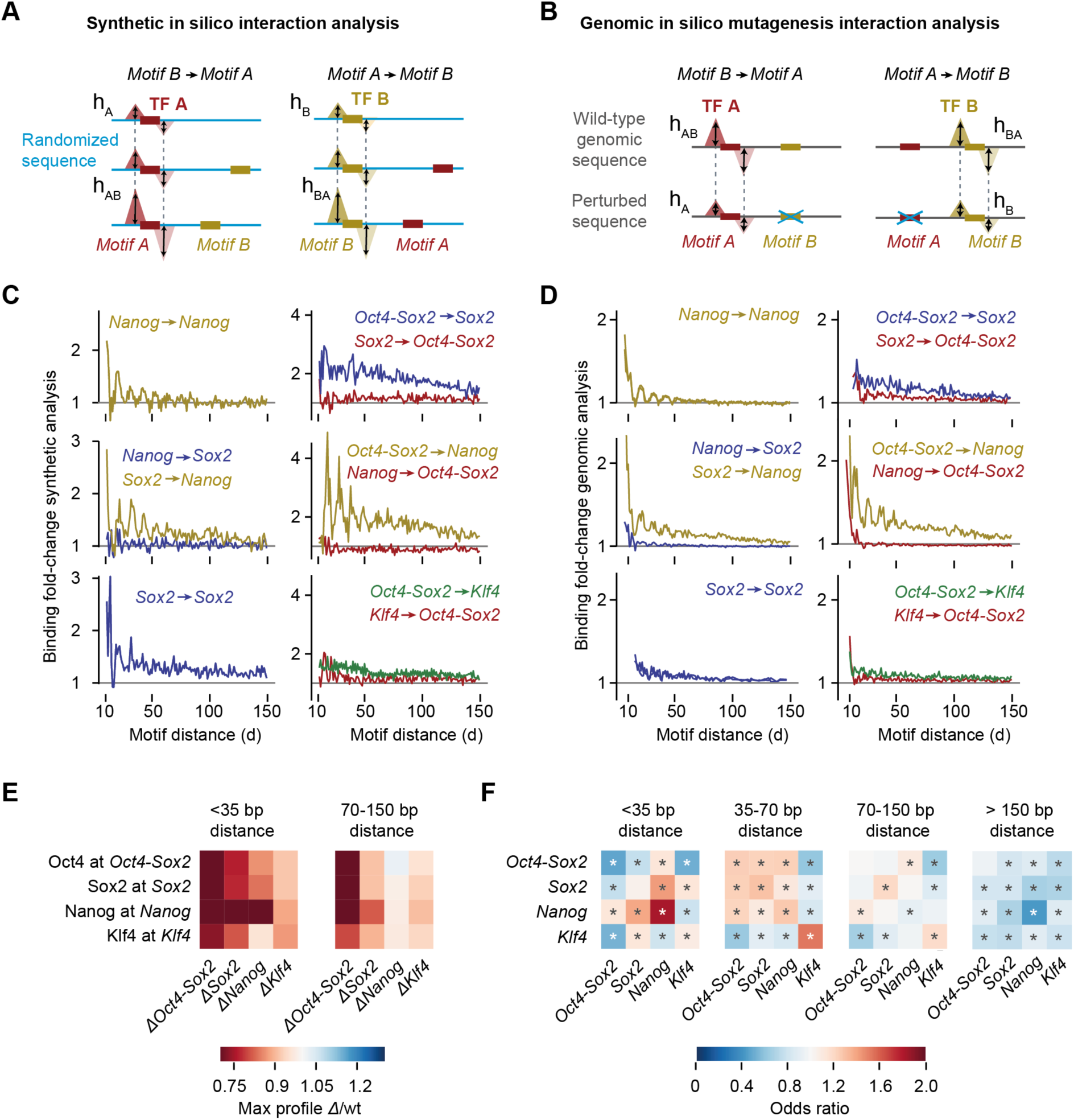
*In silico* analysis of motif interactions reveals TF cooperativity and motif syntax. **A)** *Motif A* is inserted into 128 different randomized background sequences and the average profile for TF A is predicted by BPNet. The footprint’s summits are then recorded as a reference point (dotted lines) and the height (*h*_*A*_) is measured at this position. *Motif B* is then inserted at a specific distance from *Motif A* into a new set of random sequences and the average predicted profile height at the reference summit position (dotted lines) is measured (*h*_*AB*_). To quantify the interaction between motifs as a function of distance (*d*), the fold-change of TF binding (*h*_*AB*_ /*h*_*A*_) is shown after correcting *h*_*AB*_ for shoulder effects or indirect binding footprints from the nearby motif (Figure S16A). **B)** In the genomic *in silico* motif interaction analysis, naturally occurring instances of *Motif A* and *Motif B* as determined by CWM scanning are used. The average predicted profile height and position of TF A is measured in the presence of *Motif B* (*h*_*AB*_). Then the sequence at *Motif B* is replaced with random bases, and the predicted profile height of TF A is measured at the reference point in the absence of *Motif B* (*h*_*A*_ at dotted lines). The same corrected binding fold-change *h*_*AB*_ /*h*_*A*_ as a function of *d* is used to quantify the interaction. **C)** Examples from the synthetic *in silico* analysis showing protein-range interactions involving *Nanog* and *Sox2* (left) or nucleosome-range interactions exerted by the *Oct4-Sox2* motif (bound by Oct4) on the binding of Sox2, Nanog or Klf4 on their respective motifs (right). Results are shown for the +/+ orientation of the two motifs. **D)** Similar results are obtained in the genomic *in silico* mutagenesis analysis using the average of all motif orientations. **E)** Quantification of the results shown in D as heat map. The distances < 35 bp is shown as representative for protein-range interactions, while 70-150 bp is shown as representative for nucleosome-range interactions. **F)** Odds by which two motifs are found within a specified distance from each other divided by the odds the two motifs would be found in the proximity by chance (observed by permuting the region index). * denotes p-value < 10^−5^ using Pearson’s Chi-squared test (Methods).

In the synthetic approach, *Motif A* is embedded in random DNA sequences, and the BPNet model is used to predict the fold-change in binding of TF A due to the addition of *Motif B* at a range of distances relative to the *Motif A* (*Motif B -> Motif A*, Figure 4A, Movie S1). The procedure is then repeated by anchoring *Motif B* and predicting the fold-change in binding of TF B as a function of distance to *Motif A* (*Motif A -> Motif B*, Figure 4A). The robustness of the results was confirmed by the reproducibility of the patterns across five models trained independently on different subsets of regions (Figure S17).

Using the synthetic approach on all pairwise interactions between the four motifs, we observed specific and distinct interaction patterns which were similar across all motif strand orientations (Figure S16B,C). For example, predicted Nanog binding at the *Nanog* motif was strongly enhanced when another *Nanog* motif was nearby, but interestingly, this enhancement exhibited a periodic pattern as a function of the distance between the motifs (Figure 4C). A similar periodic enhancement of Nanog binding to its motif was observed when a *Sox2* motif was nearby. The magnitude of this interaction was strongest at close distances (< 35 bp), which means it could be mediated by protein-protein interactions between Sox2 and Nanog^71,95^ or DNA-mediated allostery^4,96^. For larger distances between motifs, the increased binding of Nanog rapidly diminished, but was nevertheless still elevated further away in the presence of a *Sox2* motif (but not a *Nanog* motif). This was not true the other way around since Sox2 binding to its motif was not enhanced by a nearby *Nanog* motif (Figure 4C). Thus, BPNet predicts that Sox2 and Nanog interact and that this interaction is directional, consistent with the indirect footprints we observed.

We also observed interactions consistent with an effect associated with nucleosomes. In the presence of *Oct4-Sox2*, the predicted binding of Sox2, Nanog, and to a lesser extent Klf4, was still enhanced at nucleosome-range distances of 150 bp (Figure 4C). Interestingly, Oct4 and Sox2 have been characterized as pioneer TFs, which can bind nucleosomes and make the region more accessible for other TFs^73,97,98^. Our observed interactions suggest therefore that *Oct4-Sox2* is a strong pioneer motif. Consistent with this hypothesis, these interactions were also directional: the *Oct4-Sox2* motif greatly increased the predicted binding of other TFs, while the motifs of the other TFs did not substantially affect the predicted binding of Oct4. These differences in distance and directionality among all interactions can also be summarized as a heat map using the distance intervals of < 35 bp and 70-150 bp (Figure 4E).

In the genomic *in silico* approach, we used the original genomic sequences and predicted the binding profile of TF A to *Motif A* before and after replacing *Motif B* with a random sequence (*Motif B -> Motif A*) and vice versa (Figure 4B, example in Figure S18A). We again observed that most motif-pair interactions were directional, rather than reciprocal (Figure S18B,C,D). Overall, the interaction patterns were very similar to the synthetic approach, albeit of lower magnitude (Figure 4D, Figure S18D). The smaller effect sizes might be due to the variation in affinity of the motif instances present in the genome since the synthetic approach used the best matching sequence for each motif. It is also possible that motif perturbations can be buffered by additional motifs that are present in genomic sequences, but not in the synthetic context. In summary, both *in silico* approaches yielded similar results and pointed to protein-range and nucleosome-range cooperative interactions.

With these measurements of predicted TF cooperativity, we asked whether naturally occurring spacings between motif pairs in genomic regions might support soft motif syntax. We removed retrotransposons containing strictly spaced motifs and analyzed whether motif pairs co-occur more frequently than expected by chance at certain distances (Figure 4F, Figure S12B). The *Nanog* motifs were most strongly over-represented at short distances to *Sox2* and other *Nanog* motifs (< 35 bp), consistent with their protein-range interactions. At nucleosome-distance (70-150 bp), the *Oct4-Sox2* motif still co-occurred with *Nanog*, consistent with its pioneering role. Although BPNet is designed to capture potential motif interactions up to 1 kb apart, we did not identify significantly over-represented motif pairs beyond 150 bp (Figure 4F). Taken together, we detected genome-wide soft preferences for motif spacings that correspond to some extent with detected cooperative binding interactions and thus are likely functionally relevant motif syntax.

### Nanog binding has a strong ∼10.5-bp periodic pattern

The most remarkable soft motif syntax we observed was a ∼10.5 bp periodicity associated with Nanog. We first observed periodicity in the full-length CWM of the *Nanog* motif, which showed flanking A/T bases in a periodic pattern (Figure 5A). This pattern is not seen in the corresponding PFM representation, suggesting that the A/T bases are not statistically over-represented, but when present, contribute strongly to the Nanog binding predictions. The strong periodic pattern is confirmed in the individual contribution scores of *Nanog* motif instances, shown as heat map and average contribution scores (Figure 5B). A Fourier power spectrum analysis of the contribution scores around the *Nanog* motif revealed strong periodicity averaging around 10.5 bp (+/- 0.3 bp) (Figure 5C), which falls within the observed 10-11 bp periodicity of the DNA helix observed *in vitro* and *in vivo*^*99–102*^. This helical periodicity was also found for other motifs important for predicting Nanog binding, including *Nanog-mix, Nanog-alt, Sox2, Oct4-Sox2 and Zic3*. But the same motifs did not predict periodic binding for other TFs, suggesting that the helical periodicity is specific for Nanog binding (Figure 5D), consistent with its behavior in the *in silico* interaction analysis.

**Figure 5.**
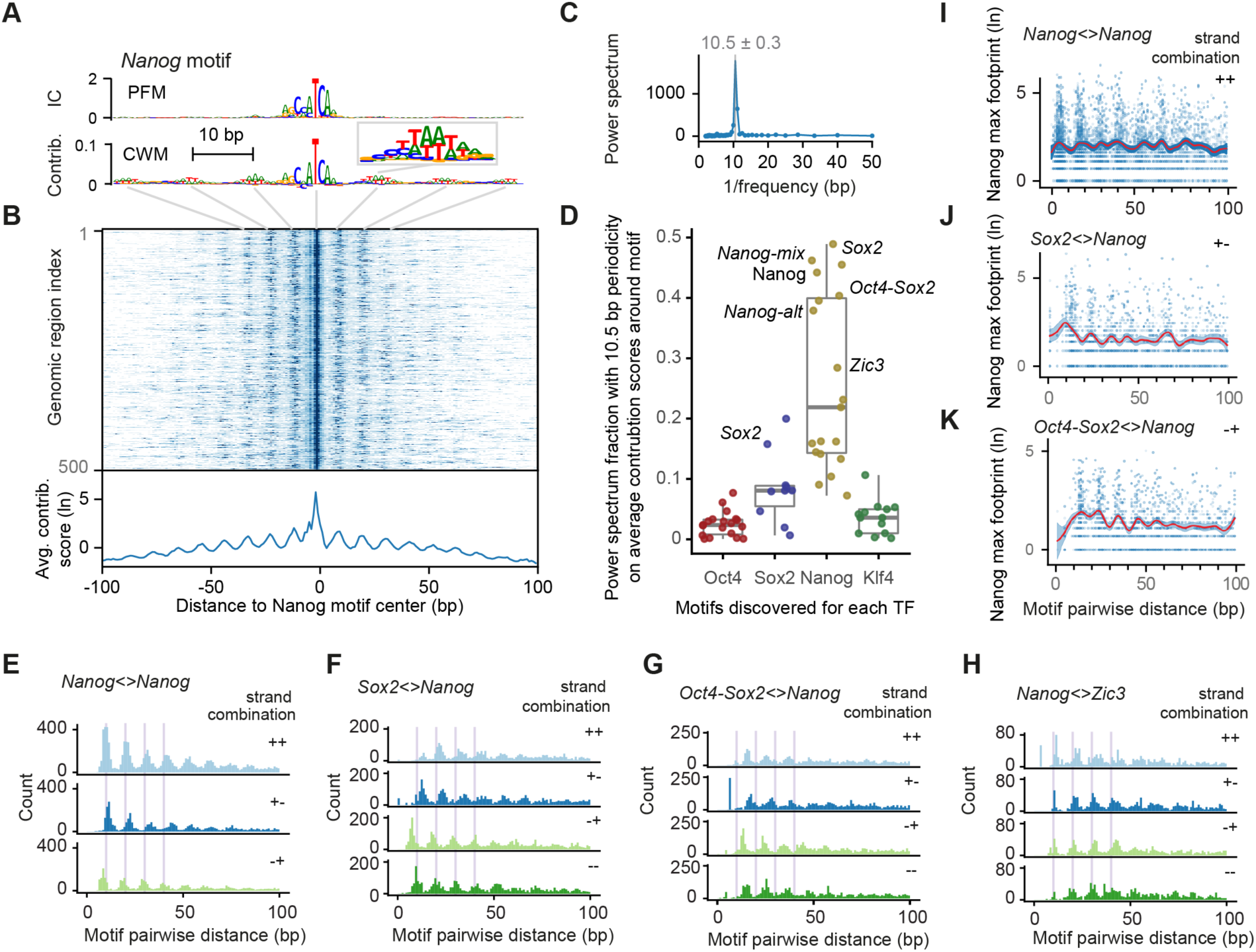
Pervasive helical periodicity between *Nanog* and partner motifs. **A)** The CWM, but not the PFM, of the main *Nanog* motif has periodically occurring contributing bases in the flanks (example in enlarged window). **B)** A heat map of the contribution scores of the individual Nanog instances also show this periodic pattern, the average of which is shown below. **C)** A Fourier power spectrum of the average contribution score around *Nanog* motif instances (after subtracting the smoothed signal) reveals an average periodicity of 10.5 +/- 0.3 bp. **D)** Fraction of the power spectrum with 10.5 bp periodicity of the average contribution scores for each TF (around each motif discovered for the TF) shows that the helical periodicity is specific for Nanog binding. Important motifs are labelled; unlabeled high-scoring motifs are from retrotransposons. **E)** The pairwise spacing of *Nanog* motif instances in all possible orientations also show a periodic pattern (++ includes the -- orientation). **F-H)** Heterologous motif combinations of *Nanog* with *Sox2, Oct4-Sox2 and Zic3* also show a preferred spacing with the same periodicity. The distance between two motifs is always kept positive by placing the second motif in the pair downstream of the first motif in the pair. All 4 motif orientations are considered: + denotes the motif lies on the forward strand and - denotes the motifs on the reverse strand. **I-K)** Nanog ChIP-nexus signal at the reference summit position for each motif instance (Figure S16A) averaged across every motif pair (blue dots), with the smooth curve fit (B-splines) depicted as a red line and the fit error bars depicted with the blue ribbon. Nanog on average binds higher when *Nanog* motifs have the preferred spacing.

To obtain further evidence of this periodicity, we tested whether Nanog’s preferred syntax was naturally found in genomic DNA sequence. Indeed, the pairwise distance between our mapped *Nanog* motif instances showed a strong helical spacing preference for a multiple of ∼10.5 bp, independent of motif orientation (Figure 5E, Figure S16B). This periodicity was reproducibly inferred from five independent models on different subsets of the binding data (Figure S19A). Despite being present in genomic DNA, the pattern had not been discovered previously^47,67–72^, presumably because traditional motif discovery and scanning methods only weakly reveal this pattern and only when specifically searching for it (Figure S9G).

The *in silico* interaction analysis also predicted enhanced periodic binding cooperativity of Nanog in the presence of other motifs. In support of this, the mapped genomic instances of *Nanog* with either *Sox2* or *Oct4-Sox2* also showed strong preferred distances of helical periodicity regardless of motif orientation (Figure 5F-G). This was also true for the distances between *Nanog* and *Zic3*, indicating that Zic3 is an additional interaction partner (Figure 5H). Furthermore, the Nanog ChIP-nexus profiles themselves also showed this periodic pattern. The average Nanog signal at *Nanog* motifs was higher when an interacting motif was present with preferred helical spacing (Figure 5I-K, 5-fold validation in Figure S19B, Figure S20). This signal in the original data likely explains how BPNet was able to learn the preferred binding pattern of Nanog during training.

The helical periodicity suggests that Nanog binding is enhanced when the relevant partner motifs are found on the same side of the DNA. Since Nanog physically interacts with Sox2^71,95^ and preferentially interacts at protein-protein distance in our *in silico* interaction analysis, it is possible that Nanog engages in cooperative protein-protein interactions similar to those observed for the lambda and lac repressors^103,104^. Alternatively, the helical periodicity could be due to preferred binding of Nanog to nucleosomal DNA from the solvent surface, which has been observed for some homeodomain TFs^105,106^.

Altogether, we identified helical periodicity as a strong cis-regulatory motif syntax for Nanog, a biophysical parameter that BPNet was not explicitly trained on. This result demonstrates the power of neural networks to discover novel patterns *de novo* without making explicit prior assumptions about the nature of the sequence features.

### CRISPR mutations validate the motif syntax between Nanog and Sox2

To experimentally validate the motif syntax identified by BPNet, we performed targeted point mutations in mapped motifs and compared the observed changes in the ChIP-nexus profiles to those predicted by BPNet (Figure 6). Since the most striking motif syntax was the helical periodicity of Nanog and the directional cooperativity with Sox2, and since the *Nanog* motif had been uncertain before^47,67–72^, we selected a genomic region that has a *Nanog* and *Sox2* motif, as well as periodic Nanog binding. Using CRISPR/Cas9 and homologous recombination, we performed two-base substitutions in either the *Sox2* motif (TTG to AGG) or the *Nanog* motif (TGA to GGC). We then performed Sox2 and Nanog ChIP-nexus experiments on wild-type and mutant ESCs, using three independently derived clones per motif mutation as biological replicates. All biological replicate experiments were highly correlated and possessed indistinguishable normalized binding across known enhancers (Figure S21).

**Figure 6.**
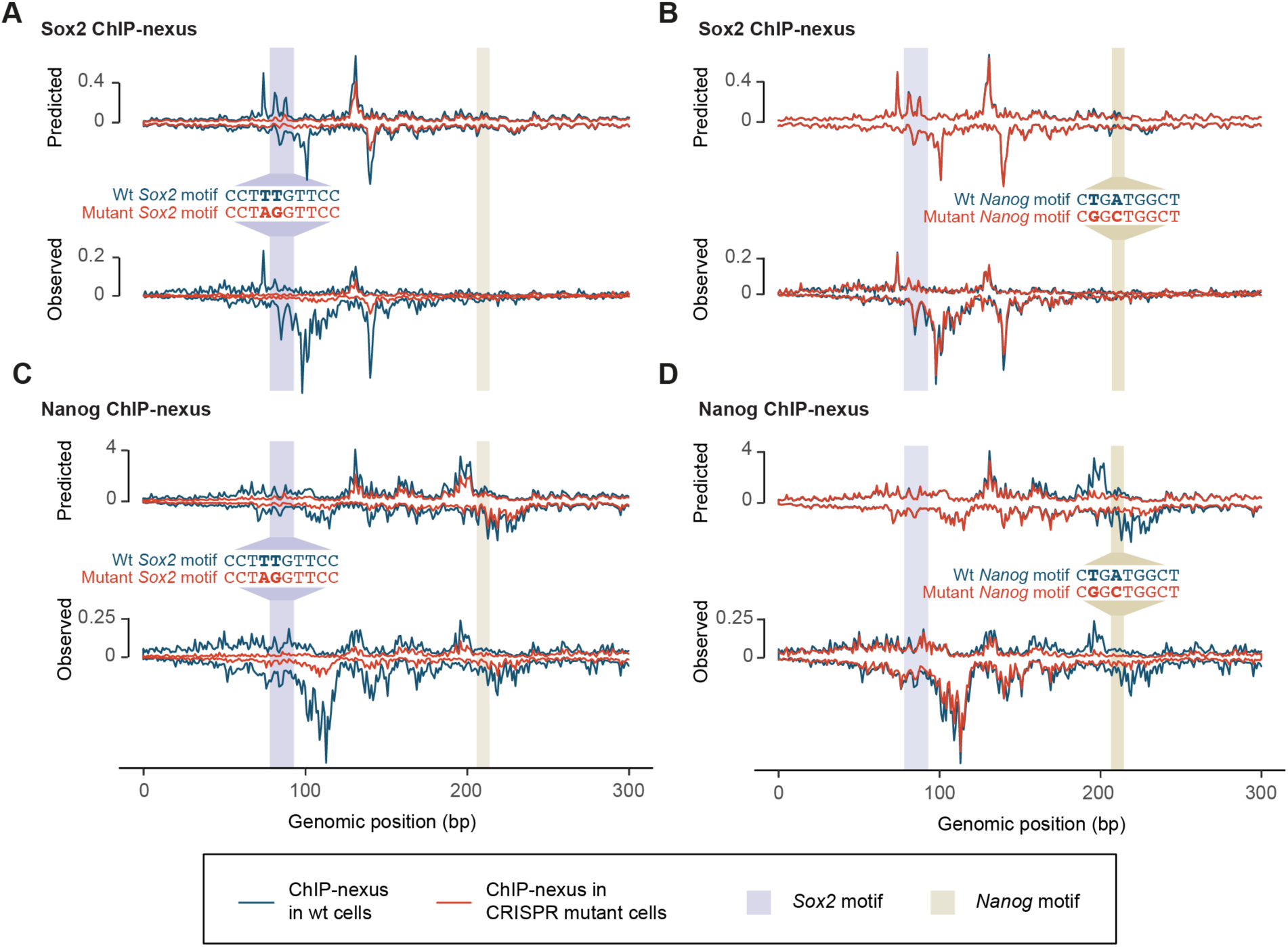
CRISPR mutations in a *Sox2* and *Nanog* motif validate BPNet’s predictions. **(A-D)** A *Sox2* motif (blue shade) and *Nanog* motif (light brown) in a selected genomic region were mutated through CRISPR/Cas9 and homologous recombination in mouse ESCs. Predicted and observed ChIP-nexus profiles (+ strand on top, - strand at bottom) in reads per million (RPM) are shown for wild-type cells (teal line) and mutant cells (scarlet line) across 300 bp (chr10:85,539,550-85,539,850). **A**) Upon mutating the *Sox2* motif, the Sox2 footprint is lost as predicted. **B)** In contrast, mutating the *Nanog* motif does not noticeably affect Sox2 binding. **C)** Consistent with directional cooperativity, the Sox2 mutation does however affect Nanog binding, which is reduced throughout the region as predicted. **D)** Similarly, mutating the *Nanog* motif not only abrogates the Nanog footprint, but also results in reduced binding nearby as predicted.

We then examined how the binding profiles were affected by the mutations. As expected, mutating the *Sox2* motif specifically abolished the corresponding Sox2 binding footprint (Figure 6A). However, mutating the *Nanog* motif did not affect Sox2 binding (Figure 6B), while mutating the *Sox2* motif strongly affected Nanog binding (see Figure S21B for a quantification of the total signal). Nanog binding was almost completely lost near the *Sox2* mutation and still reduced at the nearby *Nanog* motif (Figure 6C).

This directional cooperativity is strikingly consistent with the results from the *in silico* interaction analysis performed across all genomic sequences (Figure 4D) and with the asymmetry observed in the indirect binding footprints of Nanog and Sox2 (Figure 3C). In addition, the short-range cooperativity of Nanog was confirmed. Namely, when the *Nanog* motif was mutated, not only was the corresponding footprint of Nanog abrogated as expected, but the surrounding periodic Nanog binding was also reduced as predicted (Figure 6D).

Altogether, these results confirm that the derived syntax rules are predictive and applicable to individual examples. This demonstrate that BPNet can be used to derive novel, testable biological hypotheses on how the cis-regulatory motif syntax influences TF binding.

## Discussion

Here we introduced BPNet, a versatile and interpretable deep learning tool to learn TF motifs and the rules of syntax that best predict genomics data at base resolution. To leverage the unprecedented resolution of BPNet and showcase its ability to reveal novel biological insights, we applied it to ChIP-nexus data in mouse embryonic stem cells. The results were not only consistent with previous findings, but provided a remarkably clear picture that revealed new details and principles underlying cis-regulatory motif syntax. For example, we found that TF binding is guided by soft syntax rules, which follow clear distance or spacing preferences consistent with protein-protein interactions^16,107^, or nucleosome-mediated cooperativity^108^. These soft syntax rules have not been observed before, but represent an intermediate between the strict motif syntax associated with the original enhanceosome model^109,110^ and the very flexible syntax suggested by the billboard model^14^. The resulting TF cooperativity was often directional, extending beyond the hierarchical requirement observed for pioneer TFs ^73,108^. Finally, we observed a strong preference for Nanog to bind with ∼10.5 bp helical periodicity. Helical phasing has long been thought to be a possible element of the cis-regulatory code^21,23,103,104,109,111–114^. Our finding that the helical spacing is motif-encoded and TF-specific provides a clear guidance for this feature in future studies.

As we will outline below, BPNet represents a new paradigm for discovering relevant motifs and syntax rules underlying the cis-regulatory code. Since it is based on inferring predictive patterns using deep learning, it is a radical departure from classical methods, which are based on learning over-represented sequence patterns. In order for BPNet to outcompete classical methods, it required several important design innovations (Results, Methods and Supplemental Q&A), as well as extensive quality control and rigorous evaluations to ensure that the method works as intended (Supplemental text, Supplemental Q&A). However, the result is not just an improvement over previous deep learning methods, but represents the most powerful and general computational approach to date for deciphering the cis-regulatory code from a variety of genomics assays.

The most important innovation was the development of tools that make the trained BPNet model interpretable. For the longest time, computational models in regulatory genomics have grappled with an inherent tradeoff between prediction accuracy and interpretability, but the BPNet framework enables both. The key to its interpretability was to distill information from the entire neural network, rather than trying to directly interpret the millions of cryptic, partially redundant parameters of the trained model. This allowed us to obtain predictive motif representations and derive context-aware predictive motif instances. Importantly, by using BPNet as an *in silico* oracle, we systematically predicted the effect of mutated sequences or synthetic sequence designs, which enables us to extract the rules by which motif syntax influences TF cooperativity. Such precise oracle predictions, which are not possible with classical models, allow less scalable *in vivo* experiments such as the CRISPR editing experiments to be performed on the most interesting and promising observations.

The advantage of BPNet over classical methods is that it detects motifs and their rules of syntax in a fundamentally different way. Classical methods for motif discovery rely on motifs being over-represented over background sequences^40–43^. Similarly, existing approaches to infer syntax rules use summary statistics of over-represented co-occurrence patterns^1,19,115^. These methods have limited statistical power to test individual features present in complex cis-regulatory sequences (Supplemental Q&A). By contrast, BPNet’s vast network capacity allows it to learn complex predictive rules agnostically based on their ability to accurately predict relevant experimental profiles, without explicitly defining features *a priori*. This allows the discovery of relatively rare but nonetheless predictive motifs (e.g. *Oct4-Oct4*), as well as predictive syntax features that were not certain to be relevant such as helical periodicity or the direction of TF cooperativity.

BPNet’s approach of modeling the entire cis-regulatory sequence is also inherently better suited for deciphering the combinatorial requirements for TF binding *in vivo*. Traditionally, the gold standard for a TF binding site is that it has strong affinity for the TF in *in-vitro* experiments. Likewise, scoring motif instances using PWM models relies on statistically significant sequence matches. In both cases, a selection is typically made using arbitrary thresholds, before the role of motif combinations and syntax is considered^115,116^. However, our results suggest that *in vivo*, TF binding to a motif instance is by itself a highly cooperative process that depends on neighboring motifs and syntax. Indeed, in a given cis-regulatory context, low-affinity motif instances can be an important element of the cis-regulatory code^10,52,117^. The fact that BPNet discovered subtle predictive patterns that are not necessarily strong matches to PWM motif models (e.g. the predictive bases in the flanks of *Nanog* motifs) and outperformed classical methods for identifying motif instances that are relevant *in vivo* (Figure 2G-H, Supplemental text) suggests that modeling the predictive contribution of motif instances dependent on cis-regulatory context is a powerful new approach for discovering cis-regulatory code.

Finally, BPNet is designed to be a general and versatile end-to-end approach adaptable to a number of genomic assays. It is ideally suited to learn from high-resolution genomic data, but its base-resolution output is still beneficial for lower resolution data since it does not discard any information present in the training data profiles. For example, we successfully trained BPNet models on ChIP-seq profiles for the same TFs and obtained motifs that were highly similar to those from the ChIP-nexus models (Supplemental text: Figure S11). The number and accuracy of motif instances was lower than those from ChIP-nexus profiles models, but better than those from models trained using course-resolution binary binding labels (Supplemental text: Figure S11). Similarly, we found that BPNet can also accurately model base-resolution DNase-seq profiles^118^. These results suggest that applying BPNet to existing compendia of ChIP-seq, DNase-seq and ATAC-seq data, such as those generated by ENCODE will improve the systematic mapping of cis-regulatory motifs and their rules of syntax in a variety of cellular contexts. To foster the broad application of BPNet, we have made the entire software framework available with documentation and tutorials.

Altogether, the BPNet modeling framework promises to change future research. Learning motifs and syntax rules for a variety of genomic assays in many cell types will pave the way to a more complete understanding of the cis-regulatory code and how specific bases influence the various molecular steps associated with enhancer function. At the same time, these models provide a unique opportunity to pinpoint causal quantitative trait and disease-associated genetic variants and understand the molecular mechanisms by which they alter gene regulation. Ultimately, the ability to decipher cis-regulatory information will unlock an enormous amount of information underlying organismal development, its maintenance and opportunities for therapeutic intervention of diseases.

## Acknowledgements

We thank Mike Levine (Princeton) and Robb Krumlauf (Stowers Institute) for comments on the manuscript and Johnny Israeli for technical help at the beginning of the project. This work was funded by the Stowers Institute for Medical Research, the NIH grant 1R01HG010211 to J.Z., and the NIH grants 1DP2GM123485, 1U01HG009431 and 1R01HG009674 to A.K.. Ž.A. was supported by the German Bundesministerium für Bildung und Forschung (BMBF) through the project MechML (01IS18053F). A.S. was supported by the Stanford BioX Fellowship and the HHMI International Student Research Fellowship. Illumina sequencing was performed at the Stowers Institute for Medical Research (Anoja Perera, Michael Peterson) and the University of Kansas Medical Center Genomics Core supported by the NIH grants U54HD090216, S10OD021743 and COBRE P30GM122731-03. Generation of the CRISPR/Cas9 mouse ESC lines was performed by the following Cores at the Stowers Institute for Medical Research: Genome Engineering (Kym Delventhal, Brandon Miller, Kyle Weaver), Tissue Culture (Chongbei Zhao, Alexis Murray, Yan Wang, Olga Kenzior, Qian Jiang, Skyler Hime, Sonia Gosh) and Cytometry (Jeff Haug, Dustin DeGraffenreid).

## Authors contributions

Z.A., A.K. and J.Z. conceived the project, Z.A., A.S., A.K. and J.Z. conceived and implemented the computational methods, S.K., K.D. and R.F. performed the experiments, Z.A., M.W., A.A. and C.M. performed further computational analysis, J.Z. and A.K. supervised the project, Z.A., M.W., A.K. and J.Z. prepared the manuscript with input from all authors.

## Data and materials availability

Data used to train, evaluate, and interpret the BPNet models is available on ZENODO at https://doi.org/10.5281/zenodo.3371164. Trained BPNet models and all the model interpretation results are available on ZENODO at https://doi.org/10.5281/zenodo.3371163. BPNet model trained on ChIP-nexus data is available on the model repository Kipoi under the name “BPNet-OSKN” (http://kipoi.org/models/BPNet-OSKN/). Raw sequencing data used in this manuscript will be made available soon on the GEO archive. Code to reproduce the results of this manuscript is available at https://github.com/kundajelab/bpnet-manuscript. The ChIP-nexus data processing pipeline is available at https://github.com/kundajelab/chip-nexus-pipeline. Software to trim and de-duplicate ChIP-nexus reads is available at https://github.com/Avsecz/nimnexus/. The BPNet software package is available at https://github.com/kundajelab/bpnet/.

## Supplemental Q&A

**Q1. What are the key innovations in the design of the BPNet model compared to previous deep learning models of TF binding?**

1. BPNet is the first neural network architecture designed to model continuous base-resolution binding profiles from ChIP-nexus/exo and ChIP-seq experiments as a function of DNA sequence. This in contrast to all previous deep learning approaches that model binding data as binary binding events or continuous binding signal summarized at low-resolution (100-200 bp).
2. BPNet introduces a new multi-scale loss function to separately optimize the predictions of ChIP-nexus profile shape and total read counts.
3. BPNet introduces a new approach to automatically correct binding profiles for assay biases and prevent the model from learning spurious sequence patterns that explain assay biases. Specifically, BPNet can use a control experiment as an auxiliary input. The model is fit to the ChIP-nexus/ChIP-seq binding profiles using both the sequence and the control experiment track. The model automatically learns to regress out the signal that can be explained by the control track.

**Q2. What are the enhanced model interpretation methods introduced in the BPNet framework to infer motifs and motif syntax?**

1. Several neural network feature attribution methods (e.g. *in-silico* mutagenesis, input gated gradients, DeepLIFT, integrated gradients and GRAD-CAM) have been previously applied to deep learning models of TF binding to infer contribution scores for each base (feature) in an input sequence to a scalar binary or continuous binding prediction. Here, we introduce the first approach to infer base-resolution contribution scores for input sequences with respect to profile outputs from sequence-to-profile models. Specifically, we adapted our DeepLIFT method for profile outputs. Previously, these contribution scores have typically only been used to visualize anecdotal examples of sequence motifs in specific case study sequences or to score non-coding genetic variants. Here, we provide the first genome-wide analysis of contribution scores and show convincingly how they can accurately identify predictive motif instances genome-wide, outperforming traditional motif discovery and scanning approaches.
2. The typical approaches used to derive motifs from deep learning models of genomic sequence include visualizing the convolutional filters directly or deriving position weight matrices (PWMs) from subsequences that activate convolutional filters in the first layer. These approaches have several drawbacks since neural networks learn representations in a distributed fashion i.e. no single filter is guaranteed to capture a complete motif and there is significant redundancy in the patterns learned by different filters. Hence, interpreting filters directly often results in deriving incomplete motif patterns and many partially redundant motifs. Another drawback is that deriving PWMs that simply capture base frequencies results in a fundamental loss of information with regard to the predictive base contributions, as well as any interaction effects between bases within and across motifs. The TF-MoDISco algorithm we use here is fundamentally different in that it reconstructs less redundant and complete motif representations from the base-resolution contribution scores instead of individual filters. We show that TF-MoDISco is able to learn several novel motifs missed by other approaches for four highly studied pluripotency transcription factors.
3. We are the first to highlight the fundamental difference between contribution scores and base frequencies to derive motif representations. We introduce the contribution weight matrix (CWM) representation, which is conceptually similar to the PWM but records the average contribution rather than frequency of bases in a motif. Using transposable elements, we show very clearly the advantage of the CWM over the PWM representation. PWMs highlight the entire transposable element, whereas the CWM representation only highlights the predictive subsequences corresponding to the motifs bound by the TFs. Furthermore, the CWM motif representation for Nanog highlights the helical pattern in the flanks of the motif, whereas the equivalent PWM representation does not.
4. We develop new methods to identify predictive motif instances by scanning contribution score profiles with CWMs (derived using only sequence and the BPNet model without using the measured binding profiles). We show that this new motif scanning approach outperforms PWMs even when restricting the regions to be close to the peak summit or when augmenting the ChIP-nexus summit position for BPNet (Figure S9B). Further, the CWM scanning approach shows the most dramatic improvements for short motifs with complex periodic patterns such as the Nanog motif.
5. Finally, we combine all of these innovations in the *in-silico* oracle approach for discovering motif syntax. We develop two new approaches - one based on using simulated sequences and one based on perturbing real genomic sequences to derive robust, global rules learned by the models of how motif syntax influences transcription factor binding cooperativity. We show that these two approaches complement each other and support each other’s findings. We are the first to show that high performance neural networks of regulatory DNA sequence learn *ab-initio* subtle but critically important syntactic patterns, e.g. the pervasive 10.5 bp helical periodicity displayed by Nanog. We are also the first to show that neural networks can learn preferred soft spacing constraints between motifs that are predictive of cooperative binding. The ability of neural networks to learn these higher-order, non-linear patterns has long been known, but to our knowledge, no one has previously shown robustly that deep learning models of genomic sequences can successfully capture these higher-order patterns and that they can be extracted to reveal biologically meaningful information. Not only can we predict these patterns, but we now also validate the extracted syntax experimentally, by performing point mutations using CRISPR/Cas9 and analyzing the change in TF binding with ChIP-nexus experiments. The results clearly confirm that the periodic Nanog binding depends on the *Nanog* and the *Sox2* motifs (but that Sox2 binding does not depend on the *Nanog* motif).

**Q3. What is the conceptual difference between classical motif discovery methods like MEME/HOMER and BPNet’s motif discovery method?**

Traditional motif discovery methods (e.g. MEME, HOMER) are based on identifying statistically over-represented patterns in bound sequences relative to a background set of sequences. Our method does not rely on the frequency of sequence patterns in a foreground set of sequences relative to a background set. Instead it learns sequence patterns that are predictive of the binding profiles. Frequent patterns that have no predictive value are not learned and rare patterns that have predictive value are learned. Thus, while patterns have to be present across multiple sequences to have predictive value, our ability to discover them is not simply based on over-representation. This is exactly why we can learn several less frequent motifs predictive of ChIP-nexus footprints, while traditional methods that primarily focus on over-represented patterns miss these motifs.

The contrast between frequency of patterns and their predictive contribution to the output is the fundamental difference between the contribution weight matrix (CWM) representation that we introduce, and the classical frequency-based position frequency/weight matrix (PFM/PWM) representation (PWMs are log-odds of PFMs normalized against background frequencies). The case study of transposable elements best highlights this fundamental difference. The PFMs highlight the entire over-represented retrotransposon sequences. However, the CWMs highlight the specific predictive bases within the retrotransposons, which is a clear advantage over traditional methods.

**Q4: How is the BPNet oracle approach for syntax discovery different from classical methods?**

Previous methods derive summary statistics of over-represented or evolutionary conserved patterns of motif or TF peak co-occurrence and spacing from bound and unbound genomic sequences^1,19,115^. While these summaries are useful, they suffer from several issues. First, it is difficult to estimate the marginal effect of each property of cis-regulatory syntax (e.g. spacing between two motifs) due to systematic confounding from other syntactic properties (e.g. presence of other motifs, homotypic and heterotypic motif density) and background sequence composition of genomic sequences. Second, the number of genomic instances that sample each syntactic property may not be sufficient to obtain robust statistics. Finally, it is difficult to make conclusions about the impact of an over-represented syntactic property on cooperative TF binding without systematic perturbation experiments. These issues are typically resolved experimentally by performing *in vitro* binding experiments using libraries of carefully designed synthetic sequences that sample desired properties of interest ^6,17,112,119,120^.

Our *in silico* oracle approach that uses designed synthetic sequences mimics this experimental approach since the model is trained to predict experimental *in vivo* binding profiles. However, the model can sample substantially larger numbers of synthetic constructs by smoothly varying syntactic properties while accounting for sequence backgrounds. The *in silico* mutagenesis experiments on genomic sequences not only provide additional support for conclusions derived from the synthetic sequences, but also reduce the likelihood of making unreliable conclusions about “out-of-distribution” syntax properties that are never found in the genome.

By mimicking the experimental approach *in silico*, the oracle approach allows one to home in on precise hypotheses that can be tested using less scalable *in vivo* experiments such as the CRISPR editing experiments we present. Furthermore, the oracle approach is very general and will be useful to systematically study further syntactic properties and their joint effects in the future (e.g. trade off between affinity, motif density and spacing). These kinds of interactions would be very difficult to infer from explicit parameters in computational models.

**Q5. Can we use CWM motifs to identify motif instances in sequences that do not have experimentally measured ChIP-nexus profiles?**

Yes. The CWM scanning approach can be used to identify motif instances in any query sequence. The procedure requires two components - the input sequence and the BPNet model. These are the same two components also used for traditional PWM motif scanning (the PWM happens to the “model” in that case). The BPNet model predicts the output binding profile from the raw input sequence using a forward pass. That “predicted output” is then used to infer a contribution score profile across the input sequence using backpropagation (DeepLIFT). The CWM (also derived from the DeepLIFT contribution scores with respect to predicted profiles from all training set sequences) is then used to scan the contribution score profile of the input sequence. Nowhere in this procedure do we use the measured ChIP-nexus profiles. All components are derived just from the raw sequence using the model and the interpretation methods.

**Q6. Can BPNet be used to predict TF binding in new cell types not used in training?**

Not with the current model directly. Transcription factors typically have different genomic occupancy profiles across different cellular contexts. The direct binding motifs for most TFs are generally consistent across cell types. However, the higher-order syntactic rules and cooperative interactions with motifs of other TFs vary across cell types. A model that uses only DNA sequence as input and is trained on binding profiles of a TF in a one cell type will learn sequence features that are specific to that cell type. Because the DNA sequence of a genome is the same across different cell types, a sequence-only model of TF binding cannot predict different genome-wide TF binding landscapes in new cell types not used in training. However, the primary use-case for BPNet framework is not prediction of TF binding in new cellular contexts. Rather, we designed BPNet to enable inference of context-specific sequence determinants of TF binding. BPNet models could be extended to take as input sequence *and* cell-type specific information such as chromatin accessibility or histone mark profiles. These multi-modal models trained to account for differences in training and test cell type regulatory syntax are likely to generalize across cell types.

**Q7: What were the rigorous evaluations and independent validations that support the statistical robustness and biological validity of results?**

First, we performed extensive quality control analyses on our ChIP-nexus data, ensuring high reproducibility, sensitivity and antibody specificity (Methods). Second, we corrected for assay-specific biases by explicitly modeling control datasets (Methods) and excluded any discernible influence of mappability artifacts on BPNet’s predictions (Figure S1). Third, we showed that independently trained models using different subsets of the binding data produced highly consistent results, including the motif syntax rules (Figure S9B, S16, S19), thereby minimizing the possibility of artifacts due to memorization or over-fitting to the training data. Fourth, we found that the derived motif syntax rules were internally consistent. They were inferred from both synthetic and genomic DNA sequences (Figure 4) and the directionality was consistent with the indirect binding footprints observed in ChIP-nexus data (Figure 3). Nanog’s helical periodicity was also found in the ChIP-nexus data, the raw contribution scores, as well as the spacings between motifs (Figure 5). Fifth, the sequence representation learned by BPNet from TF ChIP-nexus data transferred seamlessly to accurately predict independent, previously published experiments, i.e. the changes in chromatin accessibility after TF depletion (Figure 2G-H). Finally, we performed CRISPR-induced point mutations in two binding motifs and showed that the changes in ChIP-nexus profiles are in remarkable agreement with BPNet’s predictions and inferred syntax (Figure 6). These careful controls and evaluations provide confidence in the ability to use BPNet as a generic toolkit for deriving biological insights about syntactic properties of regulatory DNA from ChIP-nexus and ChIP-seq experiments.

## Materials and methods

### Experiments and data processing

#### Cell culture

Mouse R1 ESCs were cultured on 0.1% gelatin-coated plates without feeder cells. Mouse ESC medium was prepared by supplementing N2B27 medium (1:1 mix of DMEM/F12 with GlutaMax supplemented with N2 and Neurobasal medium supplemented with B27, Invitrogen) with 2 mM L-Glutamine (Stemcell Technologies), 1x 2-Mercaptoethanol (Millipore), 1x NEAA (Stemcell Technologies), 3 µM CHIR99021 (Stemcell Technologies), 1 µM PD0325901 (Stemcell Technologies), 0.033% BSA solution (Invitrogen) and 10^7^ U/ml LIF (Millipore).

#### ChIP-nexus experiments

For each ChIP-nexus experiment, 10 million mouse ESCs were used. Cells were washed with PBS and cross-linked with 1% formaldehyde (Fisher Scientific) in PBS for 10 min at room temperature. The reaction was quenched with 125 mM glycine. Fixed cells were washed twice with cold PBS, resuspended in cold lysis buffer (15 mM HEPES (pH 7.5), 140 mM NaCl, 1 mM EDTA, 0.5 mM EGTA, 1% Triton X-100, 0.5% N-lauroylsarcosine, 0.1% sodium deoxycholate, 0.1% SDS), incubated for 10 min on ice and sonicated with a Bioruptor Pico for five cycles of 30 s on and 30 s off. The ChIP-nexus procedure and data processing were performed as previously described ^49^ except that the ChIP-nexus adaptor mix contained four fixed barcodes (ACTG, CTGA, GACT, TGAC) and that the PCR library amplification was performed directly after the circularization of the purified DNA fragments (without addition of the oligo and BamHI digestion). For each ChIP, 5 µg antibody was coupled to 50 µl Protein A or Protein G Dynabeads (Invitrogen). The following antibodies were used: ɑ-Oct3/4 (Santa Cruz, sc-8628), ɑ-Sox2 (Santa Cruz, sc-17320), ɑ-Sox2 (Active Motif, 39843), ɑ-Nanog (Santa Cruz, sc-30328), ɑ-Klf4 (R&D Systems, AF3158), ɑ-Klf4 (Abcam, ab106629), ɑ-Esrrb (Abcam, ab19331), ɑ-Pbx 1/2/3 (Santa Cruz, sc-888), and ɑ-Zic3 (Abcam, ab222124). At least two biological replicates were performed for each factor to obtain coverage of at least 100 million reads per TF. Single-end sequencing was performed on either an Illumina HiSeq instrument (50 cycles) or NextSeq 500 instrument (75 cycles) according to the manufacturer’s instructions.

#### PAtCh-Cap experiments

For each PAtCh-Cap experiment, 10% of sheared chromatin sample volume from 10 million mouse ESCs was used as input. Chromatin was prepared as described for ChIP-nexus. PAtCh-Cap was performed as previously described ^59^.

#### ChIP-seq experiments

ChIP-seq experiments were performed as previously described ^121^ with 10 million mouse ESCs per ChIP. For each ChIP, 5 µg of the following antibodies were used: ɑ-Oct3/4 (Santa Cruz, sc-8628), ɑ-Sox2 (Santa Cruz, sc-17320), or ɑ-Nanog (Santa Cruz, sc-30328). At least two biological replicates were performed for each factor. Single-end sequencing was performed on either an Illumina HiSeq instrument (50 cycles) or NextSeq 500 instrument (75 cycles) according to the manufacturer’s instructions.

#### Mutation of binding motifs using CRISPR/Cas9

CRISPR/Cas9 technology was used for engineering mouse R1 ES cells. The predicted *Nanog* motif on chr10: 85,539,756-85,539,765 (mm10) was mutated from C**T**G**A**TGGCT (wild-type) to C**G**G**C**TGGCT (mutant). The predicted *Sox2* motif on chr10: 85,539,634-85,539,643 (mm10) was mutated from CCTTTGTTCC (wild-type) to CCT**AG**GTTCC (mutant). GuideRNA target sites were designed using CCTop target predictor tool ^122^. The target sites were selected by evaluating the predicted on-target efficiency score and the off-target potential^123^. To generate the specific point mutations, a single stranded DNA oligonucleotide (ssODN) donor was designed containing ∼40 nucleotides of homology from the targeted cut site. A ribonucleoprotein (RNP) complex was formed by combining 90 pmol of sgRNA (ordered as Alt-R sgRNA; IDT, USA) and 10 pmol of Cas9 HiFi protein (IDT) and hybridizing for 10 min at room temperature. The RNP was combined with 100 pmol of ssODN donor and delivered to cells by Neon electroporation (1500 V, 10 ms, 3 pulses; Neon Transfection System, Model MPK5000, Life Technologies). Single cells were screened for the expected mutations through sequencing paired-end on an Illumina MiSeq instrument (250 cycles). On-target indel frequency and expected mutations were analyzed using CRIS.py ^124^. Only clones with the intentional mutation and sequence alignments above 90% were chosen for future experiments.

Per target site, three monoclonal cell lines were selected and used to perform ChIP-nexus: clones B07, B09 and F10 for the mutant *Nanog* motif, and clones B07, B11 and C10 for the mutant *Sox2* motif. In addition, at least two biological replicates of wild-type mouse R1 ES cells were prepared and used as control. ChIP-nexus was performed with 20 million ES cells and 5 µg ɑ-Nanog (Abcam, ab214549) or ɑ-Sox2 (Active Motif, 39843) per experiment as described above. Single-end sequencing was performed on an Illumina NovaSeq instrument (100 cycles) according to the manufacturer’s instructions to obtain a coverage of about 400 million reads per experiment.

#### ChIP-nexus data processing pipeline

Random barcodes and fixed barcodes were trimmed off the reads and reassigned to FASTQ labels using nimnexus (v0.1.1). The adapters were then trimmed using cutadapt (v1.8.1) ^125^. Next, the reads were aligned with Bowtie (v1.1.12) ^126,127^ using the command `bowtie -- chunkmbs 512 -k 1 -m 1 -v 2 --best --strata` to the mouse genome assembly mm10. Mapping stats were computed using SAMtools flagstat (v1.2) ^128^. Reads were filtered using SAMtools view to remove unmapped reads and mates, non-primary alignments, reads failing platform or vendor quality checks, and PCR or optical duplicates (-F 1804). Reads with poor mapping quality (MAPQ < 30) were also removed. Reads aligned to the same position with the same barcode, CIGAR string and the SAM flag were de-duplicated using nimnexus dedup (v0.1.1). The total number of final (filtered) aligned reads was 243M for Oct4, 140M for Sox2, 214M for Nanog and 176M for Klf4. The final filtered BAM file was converted to tagAlign format (BED 3+3) using bedtools ‘bamtobed’ (v2.26) ^129^. Cross-correlation scores were obtained for each file using phantompeakqualtools (v1.2) ^130^. BigWig tracks containing the strand-specific number of aligned 5’ read ends (pooled across all replicates) were generated using `bedtools genomecov -5 -bg -strand <+/->`, followed by bedGraph to BigWig conversion using UCSC bedGraphToBigWig ^131^.

Peaks were called using MACS2 (v2.1.1.20160309) by extending 5’-ends of reads on each strand using a 150 bp window (±75 bp) and then computing coverage of extended reads across both strands (shift=-75, extsize=150). For each TF, peak calling was performed on filtered, aligned reads from each replicate using a relaxed *p*-value threshold of 0.1 and retaining the top 300,000 peaks as described in ^130^. Relaxed peak calls were also similarly obtained from pseudo-replicates, which were obtained by pooling filtered, aligned reads from all replicates for a TF and randomly splitting the pooled reads into two balanced pseudo-replicates. We used the Irreproducible Discovery Rate (IDR) framework to obtain reproducible peaks across the true-replicates and pseudo-replicates ^132^. The larger of these two sets of IDR peaks (in terms of number of peaks) was defined as the “IDR optimal set” of peaks for each TF. Peaks overlaping the blacklisted regions listed in http://mitra.stanford.edu/kundaje/akundaje/release/blacklists/mm10-mouse/mm10.blacklist.bed.gz were excluded. We obtained 25,849 IDR optimal peaks for Oct4, 10,999 for Sox2, 56,459 for Nanog and 57,601 for Klf4. Regions of 1 kb in length centered at peak summits from these “IDR optimal peak sets” were used as inputs to BPNet.

We computed several quality control metrics to evaluate enrichment and reproducibility of our ChIP-nexus datasets based on the ENCODE TF ChIP-seq pipeline and quality control standards ^130^ (Supplemental table 1). We computed the fraction of reads in IDR optimal peaks (FRiP) as an estimate of enrichment. All our samples had uniformly high FRiP scores. We also computed the “rescue ratio” i.e. the ratio of the number of IDR optimal peaks from pseudo-replicates to the number of IDR optimal peaks from the true replicates, as an estimate of reproducibility. For all four TFs, ChIP-nexus samples had Rescue Ratios < 2 and had tens of thousands of reproducible peaks indicating high reproducibility of the datasets. The IDR optimal peaks from ChIP-nexus data also showed strong overlap with IDR optimal peaks from corresponding ChIP-seq data targeting the same TFs.

The nim-nexus code is available at https://github.com/Avsecz/nimnexus/. The ChIP-nexus pipeline performing the described steps (e.g. turning the raw reads in the FASTQ format to BigWig coverage tracks and the called peaks) is available at https://github.com/kundajelab/chip-nexus-pipeline. A detailed pipeline specification is available at https://docs.google.com/document/d/1h9lZ0GyVWd02RCmtaFWSaSFzrcNHoH_OgyPHMpU7b04.

#### ChIP-seq data processing pipeline

ChIP-seq datasets were processed using the ENCODE ChIP-seq pipeline https://github.com/ENCODE-DCC/chip-seq-pipeline2/releases/tag/v1.2.2. The ChIP-seq pipeline is identical to the ChIP-nexus pipeline described above except that it uses the SPP peak caller ^34^ and doesn’t use barcodes for read de-duplication.

### BPNet: base-pair resolution deep learning model

#### Architecture

BPNet is a sequence-to-profile convolutional neural network that uses one-hot-encoded DNA sequence (A=[1,0,0,0], C=[0,1,0,0], G=[0,0,1,0], T=[0,0,0,1]) as input to predict single nucleotide-resolution read count profiles. We use 1000 bp DNA sequence as inputs and 1000 bp strand-specific read count profiles for ChIP-nexus TF binding experiments as outputs. The length of the input sequence and output profiles can be easily adjusted as needed for more general use cases.

The architecture of BPNet can be compartmentalized into two parts: the body and multiple task-specific output heads. The separation of the BPNet body and head components makes the architecture more flexible, allowing the features learned in the body to be used for the prediction of multiple outputs.

The body of BPNet consists of a sequence of convolutional layers with residual skip connections and ReLU activations ^58^. The first convolutional layer uses 64 filters of width 25 bp to scan the 1 kb region for relevant sequence motifs. This layer is then followed by 9 dilated convolutional layers (64 filters of width 3 in each layer) where the dilation rate (number of skipped positions in the convolutional filter) doubles at every layer. To preserve the base-pair resolution, pooling is not used in the architecture. Thanks to a large receptive field achieved by dilated convolutions, the BPNet body is designed such that the output prediction at any position in the genome is a function of sequence patterns within +/-1034 bp around the position hence covering the whole input sequence. The model can learn a wide variety of predictive sequence patterns *de novo* including multiple sequence motifs, their positional preferences and motif combinations with different spacing and orientation constraints. The output of the final convolutional layer within the BPNet body (also referred to as the bottleneck activation map) serves as input for TF-specific output heads.

There are *2T* output heads where *T* is the number of predicted tasks (e.g. TFs). For each task, we use two output heads: i) a deconvolutional layer (filter width=25, typical ChIP-nexus footprint width) predicting the strand-specific probabilities of observing a particular read at a particular position in the input sequence and ii) a global average pooling layer followed by the fully connected layer predicting the total number of read counts aligned to the input sequence for each strand. This design allows the network to decouple learning the ‘shape’ (probability profile) of the binding profiles from the total occupancy (total read counts) over the entire input sequence. We note that for the sake of simplicity Figure 1D only shows the profile heads and not the count heads. The training occurs for all TF ChIP-nexus experiments together in a multi-task fashion. BPNet architecture (without bias correction) can be implemented in the Keras framework (v2.2.4) as follows:

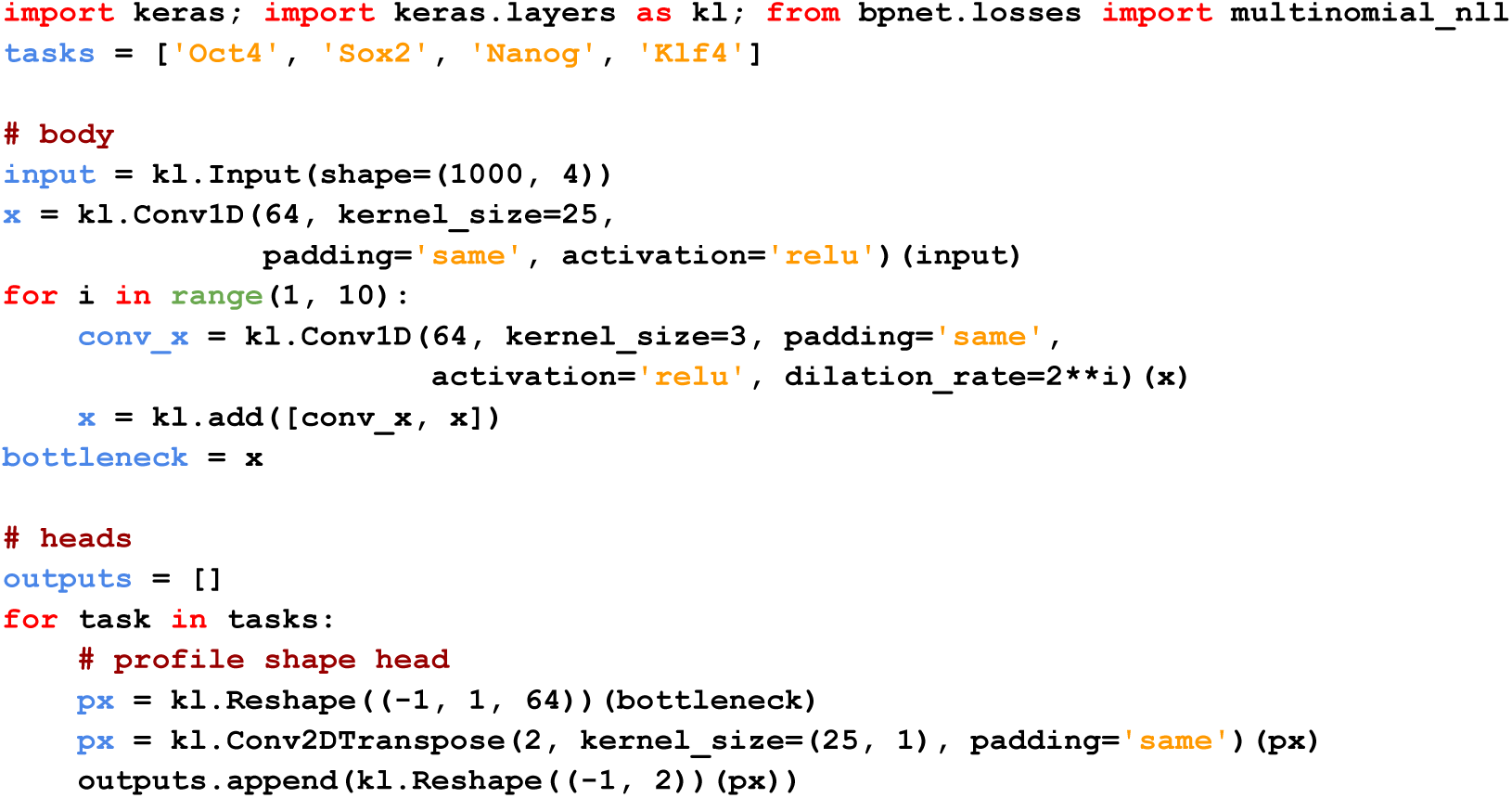

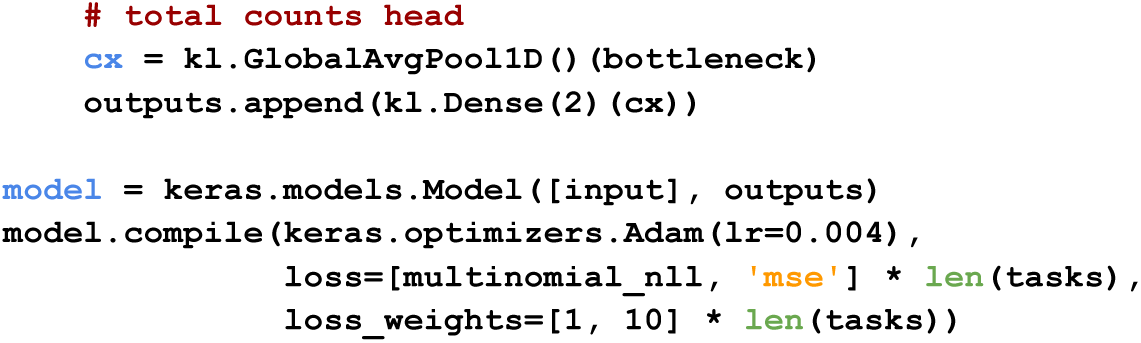

#### Loss function

Let **k**^*obs*^ be the vector of length of observed read counts for a particular strand and a particular task (i.e., transcription factor) along the sequence of length *L*. Let **p**^*pred*^ be the vector of length *L* of predicted probabilities along the sequence, such that 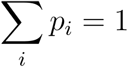 and let 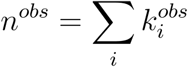 be the total number of observed counts and *n*^*pred*^ the total number of predicted counts for the sequence. BPNet is trained using the following loss function for one particular sequence, strand and task:

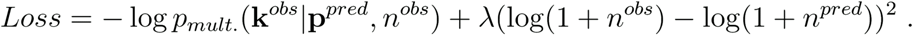

The first term evaluates the error in the shape of the predicted profile. It is the multinomial negative log-likelihood of observed base-pair read counts given the predicted probabilities and the total number of observed counts. The second term evaluates the squared error of the log total number of reads in the region. The total loss function is the sum of individual loss functions across both strands, all input sequences and all tasks (e.g. TFs).

The key question is how to choose a good value for the hyper-parameter. In supplemental text (Relationship between the Poisson log-likelihood, mean-squared error and multinomial

log likelihood), we show that if 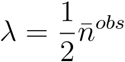, where 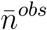 is the average number of total counts in our training set, the profile loss and the total count loss will be roughly given equal weight. As we will see later, we will use 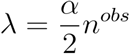 with *α* < 1 to upweight the profile predictions relative to the total count predictions.

#### Controlling for biases

Experimental assays such as ChIP-seq (and to a small extent also ChIP-nexus) have certain biases. These biases can be experimentally measured by performing control experiments such as input-DNA for ChIP-seq and PAtCh-CAP for ChIP-nexus ^59^. To prevent the sequence-to-profile model from learning these non-informative bias signals, the model tries to explain the target experimental track using both the sequence-based model predictions and the control experiment track

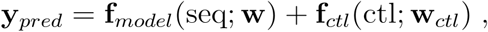

where *f*_*ctl*_(ctl: **w**_*ctl*_) is some transformation of the control track with the requirement that **f**_*ctl*_(ctl; **w**_*ctl*_) = 0 if the control track is 0 (i.e. bias not present). For the total count prediction head, *f*_*ctl*_(ctl; **w**_*ctl*_) is simply *w*_*ctl*_log(1 + *n*_*ctl*_), where *n*_*ctl*_ is the total number of reads from the control experiment in the modeled local region. For the profile prediction head, *f*_*ctl*_(ctl; **w**_*ctl*_) is a weighted sum of i) the raw counts and ii) a smoothed version of the raw counts using a sliding window sum of 50 bp. We use the sliding window to deal with typically very sparse data from the control experiment. During model training, the parameters of *f*_*ctl*_(ctl: **w**_*ctl*_) are also trained to best explain the output using the control track. We note that this framework also easily integrates multiple control tracks as well as control tracks predicted from sequence using a bias model learned on other data such as deproteinized genomic DNA for DNase-seq ^133^.

#### Training and hyper-parameter tuning

We used ChIP-nexus profiles of Oct4, Sox2, Nanog and Klf4 TFs in mouse embryonic stem cells (ESCs) to train and evaluate BPNet (≈ 100 million reads per TF, pooled from multiple replicates). The ChIP-nexus datasets exhibited high replicate concordance, signal-to-noise ratios and strong overlap of peaks with corresponding ChIP-seq experiments targeting the same TFs. PAtCh-CAP experimental data were used as the control. For each TF, the ChIP-nexus profile coverage is defined by the number of reads with the 5’ end aligned to a specific position and strand. Regions of enrichment (peaks) were identified using MACS2 ^35^ on smoothed read densities to obtain a ChIP-seq-like signal. We restrict model training and evaluation to 1 kb regions around the 147,974 peak summits of either of the four TFs in autosomes that ranked consistently across replicates genomic regions as measured by the irreproducible discovery rate (IDR) ^132^ threshold of 0.05. Regions from chromosomes 2,3,4 (20%) were used as the tuning set for hyper-parameter tuning. Hyper-parameters were manually adjusted to yield best performance on the tuning set. Regions from chromosomes 1,8,9 (20%) were used as the test set for final model evaluation. The remaining regions were used for model training.

We implemented and trained all neural network models in Keras (v2.2.4) ^134^ (TensorFlow backend v1.6) using the Adam optimizer ^135^ (learning rate = 0.004) and early stopping with patience of 5 epochs.

#### Profile evaluation metric

ChIP-nexus profiles contain TF footprints characterized by local spikes with high read counts surrounding a valley (putative TF binding site) with low read counts. Typical measures of similarity such as Pearson or Spearman correlation are not well suited to these types of profiles. To quantify the ability of the model to accurately localize footprint positions, we use a binary classification formulation to evaluate how well the model can distinguish positions with high read counts from lower read counts within each ChIP-nexus profile in the test set regions. Positions with more than 1.5% of the total number of reads in each 1kb test set region were labeled as belonging to the positive class and positions with less than 0.5% of total read counts were labeled as belonging to the negative class. These two thresholds were manually determined by visually inspecting the ChIP-nexus profiles in peak regions from the training chromosomes. The number of negative examples far outnumber the number of positive examples. Hence, we used the area under the Precision-Recall curve (auPRC) to evaluate the performance of the predicted read probability profiles relative to these binary labels. To evaluate the predictive performance at lower resolutions, we applied auPRC on binary labels and the predicted profile probabilities summarized in 2-10 bp long contiguous bins as follows: a bin was labeled as positive if there was at least one position in the bin with a positive label. If all the labels in the bin were negative, the bin was labeled as negative. Otherwise, the bin was labeled as ambiguous. For the predicted profile probabilities, the maximum value in the bin was used.

We used profiles sampled from replicate experiments to compute a corresponding upper bound for the above mentioned profile evaluation for each TF. For each TF, replicate experiments were divided into two groups with approximately equal numbers of sequencing reads. Read count profiles from one group were used as ground truth and the read counts profiles from the other group were treated as a predictor similar to BPNet. The roles of the replicate groups were then swapped and the final predictive performance was averaged across both scenarios. Random baseline was obtained by using shuffled regions for model predictions.

### Model prediction analysis in unmappable regions

Mappability of the mm10 reference genome with k-mers of length 24, 36, 50, and 100 bp as generated by ^136^ were downloaded from https://drive.google.com/drive/folders/0B1fks4X_Jjn5NDJjaE9TUmxrR28. We classified positions from the positive strand in the ChIP-nexus peaks into three groups: unmappable, mappable and ambiguous. For unmappable regions, we considered those that were not uniquely mapped by any k-mer of length up to 100 bp (value=0 in the provided uint8). For mappable positions, we considered those that were uniquely mapped with k-mers of lengths up to including 50 bp (value>0 and value<=50 in the provided uint8). The remaining positions (∼1%) were considered ambiguous and were excluded from the analysis. These positions were namely uniquely mappable by k-mers of length between 51 and 100 bp, which may or may not be longer than the used ChIP-nexus reads which were 50 bp or 75 bp long.

### DeepLIFT contribution scores for sequence-to-profile models

DeepLIFT is a feature attribution method for computing the contribution of each base (feature) in an input sequence to a specific scalar output prediction from a neural network model ^65^. DeepLIFT decomposes the difference between the output prediction based on an input sequence and the output prediction based on a neutral reference input sequence (see below for definition of reference) as an additive combination of contribution scores of all bases (*D* features) in the input sequence:

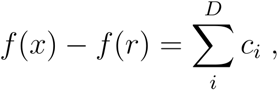

where *c*_*i*_ is the contribution of feature *i* in input *x* to the model output prediction *f*(*x*) compared to model prediction *f*(*r*) based on the reference input r. We note that *f*() is a function returning a scalar. DeepLIFT was originally developed to compute the contribution scores with respect to a single scalar output e.g. predicted output read counts at a single position on a specific strand in a profile.

For BPNet, the profile output head for a particular TF returns a 2D *L* x *S* tensor, where *L* is the sequence length and *S* is the number of output channels or strands for ChIP-nexus. Since the output of BPNet is a tensor and not a scalar, we needed to adapt DeepLIFT compute contribution scores with respect to the entire profile.

To compute base-resolution contribute scores with respect to the entire output profile, we define the profile contribution score of a base as follows:

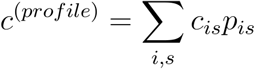

where *p*_*is*_ is the predicted probability values for position *i* and strand *s*, obtained by normalizing the profile predictions on the logit scale using the softmax function along the sequence axis: **p** = softmax(f(x)).*c* _*is*_ is the contribution score of the base with respect to the (scalar) profile prediction on the logit scale at position *i* and strand *s*. The rationale for performing a weighted sum is that positions with high predicted profile output values should be given more weight than positions with low predicted profile output values. The downside of such weighted sum formulation is that it would normally require the contribution scores to be computed *L x S* (=2,000) times for each 1 kb input sequence per TF.

To drastically speed up this computation we exploit the backpropagation algorithm used in DeepLIFT and the additive decomposition of DeepLIFT scores. We define a new TensorFlow operation as follows:

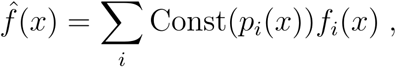

where Const denotes the `tf.stop_gradients` operation which treats the wrapped expression *p*_*i*_(*x*) as a constant. By applying DeepLIFT to 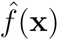 we obtain, in a single DeepLIFT backpropagation step, the desired result:

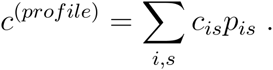

Therefore, the computational cost of computing the profile contribution scores is drastically reduced. Pseudo-code of the described operation in TensorFlow code looks as follows:

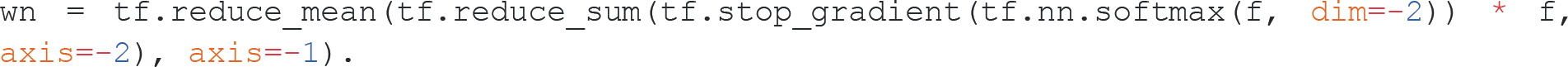

We used all zeroes for the reference input since it showed the highest correlation with in-silico mutagenesis contribution scores. The *in-silico* mutagenesis contribution scores were computed as a weighted sum of the profile prediction changes at all profile locations after introducing a mutation at a particular position. We used the DeepExplain implementation of DeepLIFT (repository fork available at https://github.com/kundajelab/DeepExplain/, commit hash: 738c7145e915a7a48f3a4248d088bcc2e1a94614) together with TensorFlow v1.6 to compute DeepLIFT contribution scores.

### Motif discovery using TF-MoDISco

We computed the DeepLIFT profile contribution scores for each TF in all 1 kb peak regions from the training, validation and test set chromosomes (i.e. peaks from all autosomes). A null distribution of contribution scores was generated by randomly selecting 4,800 peaks, extracting the sequences, shuffling them and computing the profile contribution scores for the shuffled sequences. We shuffled the sequences in such a way that dinucleotide counts are preserved. We then ran TF-MoDISco (v0.5.1.1) on each TF separately using the corresponding contribution scores of the TF in all regions where the corresponding TF was bound.

The TF-MoDISco algorithm ^31^ consists of three stages. In the first stage, the total contribution in sliding windows of length 21 (`sliding_window_size`) is computed, both for contribution scores from the real sequences and for contribution scores on the shuffled sequences. The distribution of sliding window scores on the shuffled sequences is used to define a ‘null distribution’ against which sliding windows from the real sequences that pass a FDR threshold of 0.01 (`target_seqlet_fdr`) are identified. Sliding windows are expanded on either side by 10 bp (`flank_size`) are selected in such a way that no two sliding windows overlap by more than 50%. The segments underlying these expanded sliding windows are termed ‘seqlets’, and are provided to the next stage for clustering. A total of 145,748 non-overlapping seqlets were identified. We limited the total number of seqlets to 50,000 for each run of TF-MoDISco in order to always satisfy the memory constraints (250GB).

In the second stage, seqlets are clustered into motifs. First, a similarity for each pair of seqlets is computed using the seqlet contribution scores. For a given pair of seqlets, different possible alignments of the seqlets are considered, and for every alignment, the similarity of the contribution scores is calculated using a correlation-like metric called continuous Jaccard ^31^. The best similarity across all alignments is then taken to be the similarity of the seqlet pair. The similarities of the seqlets are provided to a clustering algorithm, after transforming the similarities in a way that grants robustness to the fact that different clusters can have different densities. The clusters are found using a Louvain community detection algorithm ^137^ that automatically determines the number of clusters by optimizing graph modularity.

After the clusters have been identified, seqlets within a cluster are aligned to each other, and the coordinates of the seqlets are expanded to fill out any overhangs in the alignment. This kind of seqlet expansion makes it possible to discover motifs that are longer than the sliding window used for seqlet identification in the first stage. A Position Frequency Matrix (PFM) and a Contribution Weight Matrix (CWM) are computed from the aligned seqlets by averaging the base frequencies and the contribution scores respectively. The seqlet coordinates are then re-centered such that the region of highest contribution falls near the middle of the CWM. Because these seqlet coordinates can be slightly different from the original seqlet coordinates, the second stage is run a second time using the seqlets with the new coordinates, for added robustness.

In the third and final stage, heuristics are applied to postprocess the motifs using the default TF-MoDISco settings for version 0.5.1.1. Clusters appearing to consist of two distinct motifs are split apart, following which clusters with highly similar motifs are iteratively merged. After all merging is complete, any clusters with fewer than 60 seqlets are treated as noise and disbanded, with their seqlets reassigned to larger clusters. Finally, motifs are expanded to the length of 70 bp and then trimmed down to their final lengths by removing flanking positions with an information content (IC) of less than 8% of the information of the base with the maximal information content in the motif. Motifs supported by less than 100 seqlets or with an information content smaller than 4 bits were discarded. The PFM information content is defined as:

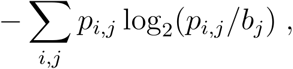

where *p*_*i,j*_ is the PFM value at position *i* and base *j* and *b*_*j*_ is the background base probability^138^. We used the following background base probabilities: A=0.27, C=0.23, G=0.23, T=0.27.

#### Identification of representative motifs

To identify and pairwise align similar motifs detected across different TFs, we performed the following motif clustering approach. First, we obtained all possible pairwise alignments of two motifs (i.e. all possible offsets and strand combinations) and identified the smallest continuous Jaccard distance metric ^31^ on the PFM information content. We then generated a pairwise distance matrix and performed hierarchical clustering in scipy (v1.2.1) using the Ward variance minimization algorithm ^139^ (`method=‘ward’`) and optimal leaf ordering ^140^. Since many of these motifs were similar or discovered multiple times by different TFs, we clustered the motifs (Figure S5B) and manually selected 11 representative TF motifs of interest.

#### CWM scanning to identify motif instances

To allow new sequences to be scored for motif instances similar to PWM scanning, we developed a method for scanning the contribution scores with the contribution weight matrix (CWM) from the TF-MoDISco motifs. We note that even though TF-MoDISco already identifies motif instances as seqlets, the detection of motif instances is not comprehensive since the number of considered seqlets (and hence the number of detected motif instances) was capped at 50,000 due to memory constraints.

There are three key differences between PWM and CWM scanning. First, a CWM instead of a PWM is used. The CWM is obtained by averaging the contribution scores of all seqlets corresponding to a specific TF-MoDISco motif. Second, in CWM scanning, the contribution scores are scanned instead of the raw sequence. Third, we use a different similarity metric between the contribution scores and the CWM. Let 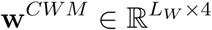 denote the CWM of length *L*_*W*_ and 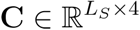 denote the contribution scores for one-hot-encoded sequence of length *L*_*S*_ ≥ *L*_*W*_. The contribution score *C*_*i,b*_ for base *b* at position *i* is 0 if base *b* was not observed in the actual sequence (i.e. if *s*_*i,b*_ = 0). We decompose the similarity metric between the CWM scanning position *i* of the contribution scores into two parts: i) the L1 norm of the contribution scores at positions between *i* and *i* + *L*_*W*_:

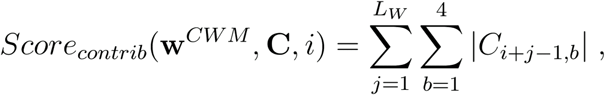

and ii) the continuous Jaccard similarity measure between the CWM and L1 normalized contribution scores:

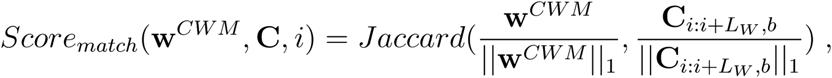

where *Jaccard* is the continuous Jaccard distance metric defined in ^31^. At each position *i*, the ‘match’ score (*Score*_*match*_) is computed for **w**^*CWM*^ and its reverse-complement version. The maximum of the two scores is used as the final ‘match’ score at each position. Note that we did not scan the ‘hypothetical contribution’ scores as performed by TF-MoDISco since we observed a higher number of false positives using that approach.

The ‘match’ score of a window in a target sequence represents its similarity to the CWM while the ‘contrib’ score determines its predictive value.

Next, we need to determine thresholds on the CWM scanning scores to call motif instances. We use the original seqlets used by TF-MoDISco to construct the CWM as a reference of high confidence motif instances to calibrate the thresholds. We compute the ‘contrib’ and ‘match’ scores for all the original CWM seqlets at their respective locations. These seqlet scores provide us with reference distributions corresponding to high confidence CWM motif instances.

We define a window in a target sequence as a motif instance if all three of the following conditions are met:

1. At least 20% of the original seqlets have a ‘match’ score lower than the ‘match’ score of the window. We use this more stringent ‘match’ score threshold (20th percentile of seqlet score distribution) in order to more effectively discriminate between instances of similar motifs. We found that lowering the threshold further often resulted in windows being spuriously assigned to several similar motifs making downstream analyses more challenging.
2. At least one of the original seqlets have a ‘contrib’ score lower than the ‘contrib’ score of the window.
3. The log odds score of the window with respect to the PWM derived from the CWM is larger than 0.

We note that CWM scanning across a target sequence does not use or need the experimentally measured TF ChIP-nexus profile associated with a target sequence. It only needs the target sequence and the trained BPNet model. The contribution scores across the target sequence used for CWM scanning are obtained from DeepLIFT by decomposing the model’s ‘predicted’ TF ChIP-nexus profile associated with the target sequence. However, the accuracy of motif instances could be further improved by using the experimentally measured and/or predicted ChIP-nexus footprints in addition to contribution scores.

We called motif instances for all sequences in the union of 1 kb wide TF peak regions (147,974) from all four TFs. Since CWMs were obtained by running TF-MoDISCo separately for each of the four TFs, we scanned CWMs derived from each TF against contribution scores of each sequence to the corresponding TF’s predicted profile. E.g. CWMs discovered by running TF-MoDISco for the Oct4 prediction task, were used to scan base-resolution profile contribution scores of all the sequences with respect to Oct4 profile prediction. We used trimmed CWMs for scanning and scoring. We removed the motif instances of short motifs which overlapped any of the motif instances matching the long motifs (PFM information content IC>30).

#### Transposable element analysis

RepeatMasker annotations for mm10 obtained from http://www.repeatmasker.org/genomes/mm10/RepeatMasker-rm405-db20140131/mm10.fa.out.gz were used to compute the overlap of seqlets with transposable elements (TEs). A seqlet was considered to overlap a TE if it was fully contained within at least one element defined in RepeatMasker annotation. Kimura 2-parameters distance ^141^ between the seqlet sequence and the consensus sequence of the motif was used to sort the seqlets in Figure S14A. This distance metric was re-implemented in Python and is equivalent to `dist.dna` function from R’s APE package with the `model=‘K80’` parameter (https://www.rdocumentation.org/packages/ape/versions/5.2/topics/dist.dna).

#### Analysis of strict spacing constraints for motif pairs

We obtained and filtered the 11 representative motif instances as described in the previous section using CWM scanning. We discarded *Sox2* sites overlapping the *Oct4-Sox2* motif and removed palindromic motif pair matches. Motif pairs were considered when spaced center-to-center between 6 bp and 100 bp. Each motif pair was checked for overlap with RepeatMasker-annotated ERVK, ERVL, ERVL-MaLR, or ERV1 genomic regions. For each motif pair, histograms were generated comparing the spacing between each motif pair instance and its ERV overlapping class. The frequency of motif spacing relative to both the motif pair and the ERV overlapping class was computed for motif pairs that occurred more than 500 times across the genome.

#### TF-MoDISco motif validation

TF-MoDISco returned three short motifs not matching the canonical Oct4, Sox2, Nanog, or Klf4 binding motifs. Two of these motifs matched TF binding motifs for *Zic3* and *Esrrb* as reported in literature. To confirm their motif identity, we performed Zic3 and Esrrb ChIP-nexus experiments and plotted their binding across the TF-MoDISco *Zic3*-like and *Esrrb*-like motifs. In both cases, this confirmed their identity. We investigated Esrrb binding across the 1,000 top-scoring genomic PWM-matches to the MA0141.1 *Esrrb* motif from JASPAR ^142^ to further verify that the PWM-matched *Esrrb* motif provided the same binding footprint as the TF-MoDISco *Esrrb* motif.

TF-MoDISco returned three *Nanog* motifs with sharp and specific Nanog binding profiles. Nanog showed differences in binding across these three *Nanog* motifs. In order to test whether a binding partner was involved, we analyzed the Sox2 and Pbx binding profiles across these three *Nanog* motifs. No binding partner was identified.

Additionally, the reported binding motif of Pbx is similar to the identified *Nanog* motifs. To ensure that the *Nanog* motif was unique to Nanog, we analyzed Sox2, Pbx, and Nanog binding across the TF-MoDISco *Nanog* and *Sox2* motif instances and the 1,000 top-scoring genomic PWM-matches to the PH0134.1 *Pbx* motif from JASPAR ^142^.

One of the three short motifs did not appear to be a known TF motif important in ESCs. We queried the TRANSFAC database ^143^ using a motif identifier tool called TOMTOM from the MEME Suite ^40^. This revealed a match with sequences associated with TFIIIC subunits. Upon further inspection, this motif was revealed to be the *TFIIIC B-box*, a binding site that contributes to the recruitment of TFIIIC binding ^144^.

#### TFIIIC B-box and tRNAs

The TF-MoDISco-returned *B-box* was the only motif identified associated with Pol III. Consistent with this motif being a Pol III motif, we found that the *TFIIIC B-box* motif frequently overlapped with tRNA genes across the mouse genome. The tRNA genes were obtained from the tRNAscan-SE predictions stored in GtRNAdb 2.0 ^145^. We then classified the *B-box* motifs based on their gene overlap and computed the copy number of the tRNAs overlapping with the mapped *B-box* motif instances based on amino acid anti-codons, separating methionine (Met) and activated methionine (iMet) as two separate amino acid classes.

### Pairwise motif interaction analysis

We studied the pairwise interaction between the following motifs discovered by TF-MoDISco:

- *Oct4-Sox2* (pattern 0 from Oct4, consensus=TTTGCATAACAA),
- *Sox2* (pattern 1 from Sox2, consensus=GAACAATGG),
- *Nanog* (pattern 1 from Nanog, consensus=AGCCATCA),
- *Klf4* (pattern 0 from Klf4, consensus=CCACGCCC).

We considered motif instance pairs (A, B) spaced at some distance *d* < 160 bp and compared BPNet ChIP-nexus profile predictions between 4 cases: where either *Motif A* or *Motif B* was replaced by a random sequence, where both were replaced by a random sequence or where both were left intact (Figure S18A). Motif instance pairs were either simulated in synthetic sequences or were detected by CWM scanning in sequences underlying ChIP-nexus peaks.

#### Synthetic sequences

For synthetic sequences, we first created 128 random background sequences of 1 kb in length by sampling the base at each position with equal probability. Next, we replaced the central bases by the consensus sequence of *Motif A* and similarly inserted *Motif B d* bases downstream of *Motif A* (*d* is the distance between motif centers). We used BPNet to predict the strand-specific ChIP-nexus profile for the primary TF of *Motif A* (e.g. Oct4 for the *Oct4-Sox2* Motif and Nanog for the *Nanog* motif). We averaged the predictions across the 128 random background sequences to obtain the profile *P*_*AB*_. We repeated the same procedure by i) inserting only the *Motif A* in the center (*P*_*A*_), ii) inserting only the *Motif B* d-bases downstream of the center, and iii) not inserting any Motif and hence only averaging the predictions across random sequences (*P*_*Ø*_). We used the predicted profile *P*_*A*_ to determine the predicted summit (maximum) location within 35 bp of the *Motif A* center for each strand. The strand-specific summit location at *Motif A* was then used to determine the profile height in all 4 scenarios averaged across the two strands. We denote the average predicted profile summit height of the 4 different predicted profiles (*P*_*A*_, *P*_*B*_, *P*_*AB*_ and *P*_*Ø*_) by *h*_*A*_, *h*_*B*_, *h*_*AB*_, and *h*_*Ø*_ correspondingly.

We define the corrected binding fold change by quantifying the influence of *Motif B* on *Motif A* as: *(h*_*AB*_ *- (h*_*B*_ *-h*_*Ø*_*)) / h*_*A*_.

A binding fold-change of 1 indicates that profile summit height of TF A is the same whether or not *Motif B* is present in the vicinity of *Motif A*. If the fold-change is higher than one, then the profile summit of TF A is higher compared to the case where *Motif B* is absent. The second term in the numerator *(h*_*B*_ *-h*_*Ø*_) corrects for predicted signal of TF A found near *Motif B* (“shoulder” effects). For homotypic motif interactions, a shoulder is present because ChIP-nexus motif footprints have a low decaying signal surrounding the summits (Figure 4E). For heterotypic motif interactions, where the TF bound to *Motif B* is different from TF A, a shoulder may nevertheless be present if TF A is predicted to show an indirect binding footprint at *Motif B* (e.g. Nanog at the *Sox2* motif, Figure 4E). By correcting for shoulder effects, we make sure that the measured interaction is not due to the indirect binding footprint at the nearby motif.

We performed the analysis for all motif pairs, strand orientations and possible pairwise distances ranging from 11 bp to 160 bp.

#### Genomic sequences

To compute the corrected binding fold-change of motif interactions in genomic sequences, we first obtained motifs instance locations in 1 kb ChIP-nexus peak regions using CWM scanning. We discarded motif instances from duplicated peak regions overlapping other peak regions by more than 200 bp as well as motif instances overlapping TEs (discovered by TF-MoDISco and mapped back to the genome using CWM scanning). Also, *Sox2* motif instances overlapping the *Oct4-Sox2* motif were discarded. For each motif pair, 4 model predictions were made:

- *P*_*AB*_: the reference sequence of the whole interval in which the motifs were present
- *P*_*A*_: motif instance B replaced by random sequence
- *P*_*B*_: motif instance A replaced by random sequence
- *P*_*Ø*_: motif instances A and B replaced with random sequence

We computed the profile heights at *motif A* profile summit locations in the same manner as for the synthetic sequences yielding 4 profile heights: *h*_*A*_, *h*_*B*_, *h*_*AB*_, and *h*_*Ø*_. We added “pseudo counts” defined as the 20th percentile of the considered quantity to the shoulder-corrected profile height of the reference sequence: *h*_*AB*_ - (*h*_*B*_ -*h*_*Ø*_) + *PC*_*AB*_ as well as the profile height of the A-only sequence: *h*_*A*_ + *PC*_*A*_. Next, we kept only the motif pairs where the shoulder-corrected profile height of the motif was in the top 20% for both motifs. This ensured that only motif pairs showing a footprint were used. Finally the corrected binding fold-change was computed for each motif instance pair as:

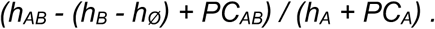

We note that there are three main differences between the synthetic and genomic sequences. First, in genomic sequences, the background sequences were not random and may contain other motifs. Second, the “perfect” consensus sequence was used for injecting motifs in synthetic sequences, whereas for genomic sequences the motif instance sequences vary and do not necessarily match the consensus. Third, the distribution of motif pairwise distances in genomic sequences is not perfectly uniform as for the synthetic case, hence some pairwise distances might be under-represented.

### Co-occurrence likelihood of motif pairs

We obtained and filtered motif instances as described in the previous section using CWM scanning. We discarded *Sox2* sites overlapping the *Oct4-Sox2* motif. To compute whether *Motif A* is located close to *Motif B* more frequently than expected by chance, we counted i) the number of times a motif instance A is close to motif instance B and ii) the number of times motif instance A is close to motif instance B if we shuffle all motif instances between peaks while maintaining the relative location within the peak. We constructed the following 2-by-2 contingency matrix *c*_*m*_:

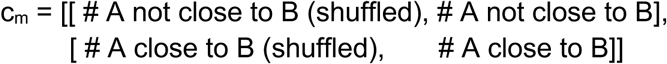

and applied the Pearson’s Chi-square test (chi2_contingency from scipy.stats) to obtain the *p*-value quantifying whether the odds-ratios (A close vs not close to B) between the observed and shuffled motif instances are significantly different. Finally, we use the odds-ratio to visualize whether A is closer to B more frequently than expected by chance:

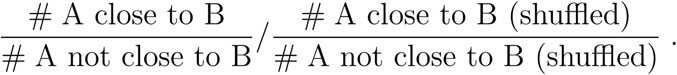

### 5-fold cross validation for analysis of robustness of BPNet models

To evaluate the reproducibility and robustness of our approach, we trained five BPNet models initialized with different random seeds on sequences from five different chromosome sets or folds. For each fold, we held-out the following validation and test set chromosomes:

|  | Validation chr | Test chr |
| --- | --- | --- |
| **Fold 1** | 1,8,9 | 2,3,4 |
| **Fold 2** | 2,3,4 | 10,11,12,13 |
| **Fold 3** | 10,11,12,13 | 14,15,16,17,18 |
| **Fold 4** | 14,15,16,17,18 | 19,5,6,7 |
| **Fold 5** | 19,5,6,7 | 1,8,9 |

Fold 1 uses the exact same validation and test chromosomes as the original BPNet. We used the exact same hyper-parameters as for the original BPNet model. TF-MoDISco and motif instance calling based on CWM scanning were run on the contribution scores for each of the five models. For each fold, the motifs most similar to the original 11 core motifs (Figure 3C) were determined by using the continuous Jaccard similarity metric for the PFM scaled by information content. Figures 5C, 6E-H and 6I-K were re-generated for all 5 BPNet models trained on different chromosome folds and their corresponding motif instances.

### Benchmarking alternative methods

#### ChExMix

ChExMix ^19,50^ is a state-of-the-art motif discovery and TF binding event calling method for ChIP-exo and ChIP-nexus data. ChExMix v0.3 with default parameters was run for each TF on the pooled BAM file containing reads of all the replicates for the corresponding TF. The same blacklisted regions (--exclude) as for peak calling in the ChIP-nexus pipeline were used. The following mm10 background file (--back) was used (http://lugh.bmb.psu.edu/software/chexmix/backgrounds/mouse.back).

#### HOMER

HOMER v2 ^**76**^ was run on the 1 kb peak regions for each of the 4 TFs profiled by ChIP-nexus with the findMotifsGenome.pl command with the following command line arguments: `-len 12 -size 200`. These specify the motif length (12) and the size of the considered regions around the peak summits (200 bp). Motif instances in the ChIP-nexus peak regions were determined using the findMotifsGenome.pl with the default arguments.

#### MEME / FIMO

MEME ^40–43^ version 5.0.2 was run on sequences extracted from the central 50 bp of the peak summits for the top 500 peaks. Command line arguments as specified by the MEMESuite webtool were used: `-dna -revcomp -mod=anr -nmotifs=3 -minw=6 -maxw=50 - objfun=classic -markov_order=0`. FIMO version 5.0.2 was used to determine the motif instances in the mm10 reference genome. We used the defaults for all parameters except for a less stringent *p*-value threshold of 0.001 (the default is 0.0001).

#### PWM-ChExMix and PWM-BPNet

Motif instances were also determined for PWMs discovered by ChExMix and BPNet. PWM score was computed using the numpy correlate function used to compute the dot-product score between the PWM and the one-hot-encoded DNA sequence. The following background probabilities were used to convert the PFM to PWM: A=0.27, C=0.23, G=0.23, T=0.27.

#### BPNet-augm

We used a sequence jittering approach to control for any potential implicit biases in the CWM scanning due to the colocalization of the summits of the ChIP-nexus profiles with the center of the input sequences used to train BPNet. For each of the original 1 kb peak regions in the test set, the predicted ChIP-nexus profiles were used to record the position of the predicted maxima (summit). Note that we never use the experimentally measured ChIP-nexus profiles. For each original 1 kb sequence, we obtained a jittered version by randomly selecting a position within +/- 200 bp of the original predicted summit and extracting the 1 kb sequence centered at this jittered center. We then used the model to infer base-resolution DeepLIFT profile contribution scores for the jittered sequence. We then scanned these contribution scores of jittered sequences with the CWMs.

#### Evaluation of validity of motif instances using ChIP-nexus profile height

We developed an approach to evaluate the validity of motif instances obtained from different motif discovery methods (TF-MoDISCo, MEME/FIMO, HOMER, ChExMix) and instance calling methods (CWM vs PWM scanning) in the absence of a ground truth set of motif instances. We expect true bound motif instances to exhibit colocalized ChIP-nexus footprints. Hence, we used the strength of the ChIP-nexus signal in the immediate vicinity of motif instances (as described below) as a surrogate measure of validity of motif instances. For all the methods, we removed all *Sox2* instances overlapping the *Oct4-Sox2* motifs. We also performed separate sets of analyses for motif instances located within +/- 100 bp and +/- 500 bp of the ChIP-nexus peak summits.

Given a set of motif instances of a motif (from each of the methods), we extracted the measured ChIP-nexus profiles centered at each motif instance. We computed an aggregate ChIP-nexus footprint of the motif by averaging the ChIP-nexus profile read counts (5’ end positions) at each position across all motif instances. We recorded the distance of the maxima of the aggregate footprint on each of the strands from the center (where the motif instances are located). We call these ‘reference summit positions’ for the motif (Figure S16A). We then compute the ‘ChIP-nexus profile height’ of each motif instance as the total number of ChIP-nexus read 5’ ends aligning to the reference footprint summit offset positions on both strands around the motif instance. It is important to note that we only compare ChIP-nexus profile height scores for motif instances from different methods within test set sequences. Using only the test set sequences ensures that BPNet derived motif instances are not implicitly using information from the measured ChIP-nexus profiles.

For each motif, a motif instance is considered to be supported by a ChIP-nexus footprint if its ‘ChIP-nexus profile height’ is greater than a predefined threshold. We selected this threshold to be the 90th percentile of the ChIP-nexus footprint height distribution over the motif instances in the test chromosome called by CWM scanning on BPNet contribution scores. This stringent threshold minimizes cross-talk from ChIP-nexus signal originating from nearby motif instances since the ChIP-nexus footprint flanks have typically 10 times lower values than the summit itself.

### Analysis of ATAC-seq data measuring differential chromatin accessibility after induced depletion of Oct4 and Sox2

#### ATAC-seq data processing

Friman *et al.* ^77^ profiled chromatin accessibility via ATAC-seq in mESCs before and after induced depletion of Oct4 and Sox2. We downloaded the corresponding paired-end ATAC-seq FASTQ files for both replicates of the Sox2 26h (ON & OFF) and the Oct4 S2iL (ON & OFF) from GSE134680 ^77^. We processed the ATAC-seq data using the ENCODE ATAC-seq pipeline (v1.5.3) at https://github.com/ENCODE-DCC/atac-seq-pipeline. We trimmed adapters from the reads in the FASTQ files using cutadapt (v1.9.1) with parameters -e 0.1 -m 5 _125_. Next, the trimmed reads were aligned to the mm10 reference genome assembly with Bowtie2 (v2.2.6) ^126,127^ using the parameters `-X2000 --mm -k 5` (report up to 5 distinct, valid alignments). Mapping stats were computed using SAMtools flagstat (v1.2) ^128^. Reads were filtered using SAMtools v1.7 view to remove unmapped reads and mates, non-primary alignments, reads failing platform or vendor quality checks, and PCR or optical duplicates (-F 1804) marked using Picard v1.126 MarkDuplicates. Reads mapping to more than 4 locations were discarded. For the remaining reads, the alignment with the best score is retained. The final filtered BAM file was converted to tagAlign format (BED 3+3) using bedtools ‘bamtobed’ (v2.26) ^129^.

Peaks were called using MACS2 (v2.1.1.20160309) by extending 5’-ends of reads on each strand using a 73 bp window (±36 bp) and then computing coverage of extended reads across both strands (`shift=-36, extsize=73`). Peak calling was performed on filtered, aligned reads from each replicate using a relaxed *p*-value threshold of 0.01 and retaining the top 500,000 peaks as described in ^130^. Relaxed peak calls were similarly performed on pseudo-replicates, which were obtained by pooling filtered, aligned reads from all replicates for each sample and randomly splitting the pooled reads into two balanced pseudo-replicates. We identified two types of reproducible peaks. First, ‘naive overlap peaks’ were defined as relaxed peaks obtained from pooled reads that overlapped relaxed peaks from both true replicates or both pooled-pseudoreplicates. Furthermore, we used the Irreproducible Discovery Rate (IDR) framework to obtain more stringent, rank consistent, reproducible peaks across the true-replicates and pseudo-replicates ^132^. The larger of these two sets of IDR peaks (in terms of number of peaks) was defined as the “IDR optimal set” of peaks. Peaks overlaping the blacklisted regions from http://mitra.stanford.edu/kundaje/akundaje/release/blacklists/mm10-mouse/mm10.blacklist.bed.gz were excluded.

#### Models for predicting differential chromatin accessibility from derived types of sequence features

The Sox2 26h (ON / OFF) ATAC-seq sample and the Oct4 S2iL (ON / OFF) samples were selected from GSE134680 ^77^ to compute the differential ATAC-seq signal (log fold-change) upon the depletion of Oct4 or Sox2 in each of the 1 kb ChIP-nexus peaks. Let represent the number of ATAC-seq 5’ read ends aligned to the *i*-th ChIP-nexus peak 1 kb region when Sox2 (TF, not motif) has been depleted and 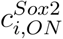 when Sox2 has not been depleted. Let 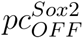 represent the 10th percentile of the 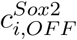 distribution across all i. We define the differential ATAC-seq log-fold change signal as follows:

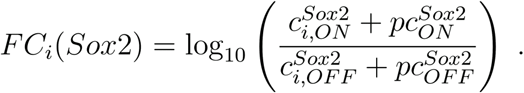

For each of the depletion experiments, we trained separate linear models (LinearRegression from scikit-learn) to predict the differential ATAC-seq log fold change signal at specific regions based on two different sets of sequence features derived from the corresponding 1 kb input DNA sequences. We restricted to regions corresponding to ChIP-nexus peaks of the four TFs that also overlapped IDR optimal ATAC-seq peaks from the wild-type (unperturbed) condition. We used the same train/test split by chromosomes as for the original BPNet training. Regions from test set chromosomes 1, 8 and 9 were used to evaluate the performance of the model. Regions from validation/tuning set chromosomes 2, 3, and 4 were not used. Regions from the remaining chromosomes were used to train the models. We evaluated the performance of the models using the Pearson and Spearman correlation metrics.

The first set of sequence features represent the complete sequence representation learned by the BPNet model trained on ChIP-nexus profiles of Oct4, Sox2, Nanog and Klf4. For each 1 kb input sequence, we computed the bottleneck activation features from the original BPNet model averaged across the spatial axis followed by a log transformation yielding 64 features.

The second set of features were derived from motif instances mapped to the 1 kb regions. For each 1 kb input sequence, we counted the number of motif instances of each of the 11 main TF-MoDISCo motifs. For each motif, we also computed the mean, maximum and sum of the motif match scores (PWM log odds) across all motif instances in the input sequence. We thus obtained a total of 44 sequence features. If no motif instances were mapped to a region, we set all the feature values for that motif to 0. We computed these features for motif instances mapped by BPNet and all other methods described in the previous section (i.e. ChExMix, MEME/FIMO, and HOMER).

#### Overlap of motif instances from different methods with differentially accessible sites due to Oct4 or Sox2 depletion

Friman *et al.* ^77^ profiled chromatin accessibility via ATAC-seq in mESCs before and after (ON and OFF, respectively) induced depletion of Oct4 and Sox2. After profiling, they classified differentially accessible states via peak calling and differential enrichment analysis between the ON and OFF ATAC-seq experiments for Oct4 and Sox2, annotating regions as Oct4-dependent (OD), Sox2-dependent (SD), or Co-dependent (CD). First, we downloaded these annotated differentially accessible regions from GSE134652. Then, we collected the *Oct4-Sox2* and *Sox2* motif instance sets mapped by BPNet, MEME/FIMO and HOMER as described above. For each motif set, we removed any *Sox2* motif instance that overlapped with an *Oct4-Sox2* motif instance. Next, we ranked each motif instance in decreasing order. BPNet motif instances were ranked by the weighted contribution scores. MEME/FIMO and HOMER motif instances were ranked by their respective motif match scores. Next, we computed the cumulative overlap fraction across each rank step for OD, SD, and CD regions using the following equation:

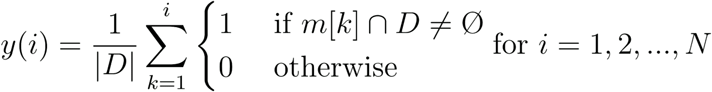

Given a ranked set of motif instances *m* of length *N* and a set of differentially accessible regions *D* (OD, SD, or CD), we counted the overlap occurrences, up to rank *i*. We then obtained the overlap rate by dividing by the differentially accessible set length *|D|*. When computing the overlap occurrences, if a region in *D* occurred more than once, only the first overlap was counted. The cumulative overlap fractions of ranked motifs were then compared between BPNet, MEME/FIMO and HOMER.

### Protein structure visualizations

The structure of Sox2 and Oct1 bound to DNA in Figure 3A was rendered in VMD ^146^ using secondary structure information from STRIDE ^147^ and surfaces from SURF ^148^, based on the NMR structure 1O4X ^149^. This Sox2-Oct1-DNA model has been used as a homology model to build the *Oct4-Sox2*-DNA complex ^149^, and is therefore representative of the structure of that complex, though coordinates for that model have not been made available.

### Periodicity analysis using Fourier transform

For each TF-MoDISCo motif of a TF, we identified the locations and sequences containing all the seqlets of the motif. We extracted base-resolution DeepLIFT profile contribution scores (w.r.t. to the TF’s profile prediction) across 200 bp sequence windows centered at each of the seqlets. We computed the average contribution score at each base across all the extracted contribution score profiles. We subtracted a smoothed version of the average contribution score (smoothing window of size 10) in order to correct for overall higher contribution scores in the center. We call this the corrected average contribution score profile of a motif.

For each motif, we computed Discrete Fourier transforms of the corrected average contribution score profile using the Numpy (v1.16.1) function `numpy.fft.rfft`. Power spectrum was obtained by taking the squared absolute value of the returned Fourier coefficients. The finite length of the corrected average contribution score profiles, results in discrete frequency values from the Fourier transform. We used half of the difference between adjacent frequency values as an estimate of the error-bars (uncertainty) around the discrete frequency values value (e.g. Figure 5C).

### Software availability

Code to reproduce the results of this manuscript is available at https://github.com/kundajelab/bpnet-manuscript. We also streamlined and generalized this code into a `bpnet` python package (https://github.com/kundajelab/bpnet/) with functionality to train and interpret base-resolution deep neural networks trained on the coverage tracks of any functional genomics assay. The ChIP-nexus data processing pipeline which includes read trimming, mapping, peak calling and generating the coverage tracks is available at https://github.com/kundajelab/chip-nexus-pipeline. The nimnexus software package for trimming and deduplicating ChIP-nexus sequencing reads is available at https://github.com/Avsecz/nimnexus/.

## Supplemental text

### Comparison of motifs and motif syntax derived from BPNet and other methods

#### PWM scanning and ChExMix

##### Comparative evaluation of motif discovery

To evaluate the extent and quality of motifs discovered by the BPNet framework in the light of previous methods, we compared our approach to ChExMix ^50^, MEME ^40–43^, and HOMER ^76^. We used each of these methods to discover motifs using ChIP-nexus peaks for each TF (Oct4, Sox2, Nanog and Klf4) (See Methods for specific parameters). For each of the methods (besides BPNet), we selected the motifs with the closest match to the 11 core TF-MoDISCo motifs discovered using BPNet (which were carefully analyzed and experimentally validated in this manuscript (Figure S9A)). We found that ChExMix, MEME, and HOMER individually discovered at most only 6 out of 11 motifs. They collectively found 9 out of 11 motifs. Only *Oct4-Sox2, Sox2*, and *Klf4* motifs were discovered by all four methods. Zic3 and Esrrb motifs were only discovered by BPNet and HOMER. The *B-box* motif was only discovered by BPNet and MEME. We speculate that these motifs were missed by ChExMix due to the associated footprints being heterogeneous and fuzzy with relatively lower read coverage. MEME/HOMER may have missed some of these since they are not as over-represented as the primary motifs. Moreover, the other methods were limited in their ability to discover long TEs motifs with the default parameters. Although changing the parameters for the other methods may allow the discovery of some TEs, the dependence on these parameters makes it difficult for the methods to discover TEs alongside short motifs in a flexible and robust manner. Moreover, these other methods cannot highlight the short constituent motifs bound by the TFs within the longer TE motifs. These results suggest that the BPNet framework provides substantial improvements in motif discovery compared to ChExMix, MEME, and HOMER.

##### Comparative evaluation of motif instances

To evaluate the quality of the called motif instances in the genome in terms of their false-positive rates, we compared the BPNet approach using CWM scanning to classical position weight matrix (PWM) scanning. To determine the false positive rate of the motif instances in the test chromosomes, we considered motif instances supported by strong ChIP-nexus footprints (‘ChIP-nexus profile height’ (See Methods) above the 90th percentile) as true binding sites (Figure S9B). Since the number of motif instances depends on the motif scoring threshold, rather than using a fixed threshold, we instead evaluated the true-positive fraction across the ranked list of motif instances. PWM scanning approaches only use the input sequences and the PWMs. CWM scanning uses TF-MoDISCo CWMs, the input sequence and DeepLIFT contribution scores to the predicted (not measured) ChIP-nexus profiles which are derived from the BPNet model. We performed motif instance evaluation only on 1 kb input sequences from the held-out (test) chromosomes, which were not used to train BPNet. Hence, both BPNet CWM scanning and PWM scanning approaches (FIMO and HOMER) do not implicitly or explicitly use the measured ChIP-nexus profiles to call or score instances. ChExMix uses the measured ChIP-nexus profiles to call motif instances. CWM based motif instances were ranked based on their ‘motif contribution scores’. PWM based motif instances were ranked based on PWM match log-odds scores. We found that across matched number of ranked instances, CWM motif instances had substantially more overlap with strong ChIP-nexus footprints (‘true positives’) and thereby exhibited a lower false positive rate compared to PWM based motif instances from FIMO and HOMER as well as ChExMix (Figure S9B, top).

Since this evaluation was performed for motif instances in 1 kb sequences centered at ChIP-nexus peak summits, most of the motif instances tend to be located near the center. To show that the improvement of BPNet is not primarily coming from the fact that it could prioritize motifs in the center higher, we computed the contribution scores with BPNet on sequences jittered +/- 200 bp around the peak summit (BPNet-augm). We found that this jittering resulted in negligible drop in performance, indicating that the superior performance of CWM scanning is not due to an implicit positional bias.

Furthermore, we repeated the evaluation by restricting motif instances to the central 200 bp around the ChIP-nexus peak summits (Figure S9B, bottom). This scenario is commonly used with PWM scanning to reduce the false discovery rate, because true motif instances are more likely to be found closer to the peak summits. Even in this scenario, BPNet’s CWM scanning outperformed all other methods for all motifs. The only exception was *Oct4-Sox2*, where several PWM methods performed as well as CWM scanning, likely due to the high information content of this particular motif. The largest improvement of BPNet CWMs over PWM methods was seen for the short Nanog motif. Even though the CWM has the same length as the PWM, the base-resolution contribution scores scanned by the CWM are dependent on the entire sequence context of the motif within the 1 kb region and thus can integrate more contextual information relevant for TF binding. E.g. a *Nanog* motif instance can get a higher contribution score if it is present in the vicinity of other ∼10.5 bp spaced *Nanog* motif instances. In contrast, the PWM scores sequence matches of each sliding window within the input sequence independently and is unable to account for the influence of surrounding nucleotides and motifs.

Finally, we note that the superior performance of CWM scanning over PWM scanning is highly reproducible when evaluated based on independent ChIP-nexus experiments using a different Nanog antibody (Figure S9B, last column). Hence, our approach of scanning the contribution scores using the CWM (instead of the raw sequence using the PWM) greatly reduces the false positive sites while still following the familiar scanning procedure as with PWMs. These results highlight the advantages of using profile contribution scores and the novel CWM motif representation to identify motif instances associated with ChIP-nexus footprints.

##### Comparative evaluation of helical motif syntax

Next, we tested whether the ∼10.5 bp helical *Nanog-Nanog* motif syntax discovered by BPNet could also be discovered using motif instances from the other methods (HOMER, MEME, ChExMix). We found that the methods showed substantially weaker signals of helical periodicity in the *Nanog-Nanog* pairwise spacing histograms across a range of distances (Figure S9C). For PWM scanning, the higher false discovery rate of motif instances likely attenuates the detection of Nanog’s helical periodicity. For ChExMix, we observed a substantial depletion of helical periodicity for spacing < 40 bp compared to BPNet CWM scanning. This depletion at close proximity could be due to two reasons. First, the optimized likelihood of ChExMix is non-convex and hence the global optimum might be difficult to find and may strongly depend on the initial conditions. Second, the key assumption of ChExMix is that the tag distribution (representing the average profile) associated with a specific motif is constant. However, this assumption is an oversimplification since ChIP-nexus profiles associated with a motif can change their form in the presence of motifs of other cooperatively bound TFs. Altogether, these results demonstrate that compared to previous approaches, the BPNet framework provides substantially improved sensitivity for detecting subtle, closely-spaced, soft motif syntax since CWM scanning yields more accurate motif instances without relying on the measured ChIP-nexus profiles.

#### BPNet’s profile regression yields more motifs and more accurate motif instances than binary peak classification

A frequently used approach for training deep learning models is to treat the TF binding prediction problem as a binary classification task ^25,26^. In this approach, the training examples are sequences extracted from contiguous bins in the genome and the sequence label is positive if a TF binding peak overlaps the bin region (and negative otherwise). The purported benefits of such a labeling approach are as follows. First, the assay-specific biases may be already accounted for in the peak-calling process. Second, the resulting machine learning task – binary classification – is well understood. Hence the standard loss function such as binary cross-entropy and the standard evaluation metrics such as the area under precision-recall curve (auPRC) can be used. However, this course-grained binary summarization of the binding profiles discards valuable fine-scale information about signal strength and shape of the profiles.

To investigate the benefit of training the BPNet model on the base-resolution ChIP-nexus profiles compared to lower resolution binary labels, we modified the BPNet architecture and replaced the output heads performing profile regression with output heads performing binary classification. The new output heads consisted of weighted global average pooling using spline transformation ^150^ and a dense layer followed by sigmoid activation. We trained the model on 1 kb input sequences sampled every 50 bp across the training chromosomes. Sequences were labeled positive if the central 200 bp of the sequence overlapped an IDR-optimal peak. The predictive performance on the held-out tuning chromosomes (2, 3 and 4) was 0.25 auPRC on average across the 4 TFs after tuning the optimal learning rate (Figure S10A). We also observed that chromosome-wide training of the binary classification models took 3 times longer (Figure S10B) than BPNet, which is trained only on ChIP-nexus profiles from 147,974 peak regions. To ensure that the dilated convolutional layers are also appropriate for binary classification, we also trained and evaluated the Basset ^29^ and factorized Basset ^151^ architectures. After tuning the dropout rate with random search, we obtained a slightly lower auPRC of 0.24 for both models, suggesting that our original architecture with dilated convolutions was also a good fit for binary classification. Next, we asked whether the predictive performance of the binary classification model could be improved by adding another output head predicting the strand-specific ChIP-nexus profile as originally done by BPNet. Indeed, the classification performance increased for all TFs yielding an average of 0.31 auPRC (Figure S10A). We conclude that the read profiles indeed provide additional information not captured by the binary labels.

We next asked whether the contribution scores of the profile regression model highlight additional motifs compared to those obtained from the binary classification model. For the binary models, we computed the DeepLIFT contribution scores for each TF task (pre-sigmoid activation) and ran TF-MoDISco with the same parameters in the same regions as previously done for BPNet. We clustered the discovered motifs based on their PFM similarity and manually assigned motif labels as done in Figure S5B. Using the contribution scores of the binary classification model, TF-MoDISco discovered 9 out of 11 main short motifs found by the profile regression model (Figure S10C, Supplemental Table 2). The 2 missed motifs, *Oct4* monomer and *B-box*, are hence not frequently used by the binary model to predict the presence or absence of the peak as they might co-occur with other more predictive motifs. Interestingly, a higher number of questionable motifs including GC sequence composition bias motifs, ambiguous motifs, and degenerate or noisy motifs were discovered from the contribution scores of the binary classification model. This suggests that the contribution scores of the binary classification model are noisier than for the profile regression model. Nevertheless, we note that the high reproducibility of the discovered motifs using two different model architectures trained on different labeling schemes for the same underlying data demonstrates the robustness of TF-MoDISco.

To compare the accuracy of motif instances for the 4 cognate motifs discovered by TF-MoDISco for both models (*Oct4-Sox2, Sox2, Nanog* and *Klf4*), we performed the instance ranking analysis on test set sequences, as for PWM scanning methods in Figure S9 considering sites with strong ChIP-nexus profile heights (footprints) as ‘true’ binding sites. The contribution scores of both models yielded a similar recall of *Oct4-Sox2* and *Sox2* motifs supported by strong ChIP-nexus footprints (Figure S10D). Strikingly, the contribution scores of motif instances from the BPNet profile model recalled a much higher fraction of *Nanog* motifs with strong footprints as compared to those derived from the binary models (Figure S10D). Since the *Nanog* motif is frequently found in complex homotypic or heterotypic syntactic arrangements with *Sox2*, the ChIP-nexus profile shape contains rich information reflecting these motif arrangements. Since BPNet was trained on ChIP-nexus profiles directly, it is able to learn these subtle patterns and encode them in the contribution scores, thereby resulting in more accurate motif instances even on unseen sequences in the test set. Additionally, CWM scanning of contribution scores from the binary classification model is comparable to PWM scanning (MEME/FIMO), suggesting that BPNet trained on ChIP-nexus profiles is the key for accurate motif maps.

Altogether, we observe that learning to predict the full ChIP-nexus profiles as done by BPNet instead of lower resolution binary classes reduces the training time by three fold, increases the number of discovered motifs with strong seqlet and biological support, reduces the number of spurious low motifs discovered and improves the accuracy of the called motif instances. Moreover, the profile predicted by BPNet assesses binding at individual motifs, which offers higher resolution to study the directionality of TF interactions mediated by motif syntax as shown in Figure 4.

#### BPNet can also be used to model and interpret transcription factor ChIP-seq profiles

The BPNet model together with the interpretation workflow using DeepLIFT and TF-MoDISco can be readily applied to any regulatory profiling experiment such as ChIP-seq, since it does not make any modeling assumptions specific to ChIP-nexus profiles. The major difference between ChIP-seq and ChIP-exo/nexus is the resolution. For ChIP-seq, the 5’ ends of the reads mapping to either strand within an enriched peak region are dispersed in a 100-200 bp window around the primary binding site (peak summit). In contrast, for ChIP-exo/nexus data, the read density is colocalized in the immediate vicinity (+/- 20 bp) of binding events.

To demonstrate that BPNet can also model ChIP-seq profiles, we performed ChIP-seq for 3 out of 4 previously studied TFs (Oct4, Sox2 and Nanog). We processed the data using the ENCODE ChIP-seq pipeline (v 1.3.6) https://github.com/ENCODE-DCC/chip-seq-pipeline2/releases/tag/v1.3.6 and generated the strand-specific 5’ read count tracks as for ChIP-nexus. We then optimized multi-task BPNet architectures to predict strand-specific ChIP-seq profiles of the three TFs from their corresponding 1 kb sequences at IDR optimal peak regions across all three factors. We used the same BPNet architecture for ChIP-seq as for ChIP-nexus and determined optimal hyper-parameters by varying each hyper-parameter individually while keeping others the same as for the ChIP-nexus model (learning rate: 0.05, 0.04, 0.02, 0.01, 0.005, 0.004, 0.002, 0.001, 0.0005; deconvolution size: 1, 10, 20, 30, 40, 50, 60, 70, 80, 100; number of layers: 1-12; profile vs total count loss weight λ: 1, 2, 5, 10, 20, 50, 100, 200, 500, 1000, 2000, 5000, 10000). We observed that the BPNet model for ChIP-seq overall required the same hyper-parameters as for ChIP-nexus. The only hyper-parameter that differed was the increased width (50) of the deconvolutional layer (compared to 25 which was optimal for ChIP-nexus). Similar to the ChIP-nexus control experiment PAtCh-Cap, we used the ChIP-seq input control experiment using a non-specific antibody to control for experimental biases (Methods). We also added data augmentation (genomic intervals jittered uniformly by [-200, 200] bp with random reverse complementation). This is more important when ChIP-seq data are trained on peaks only since the shape of the profiles will be fairly constant, hence a constant model can already fit the data well.

To gain intuition about the prediction quality of BPNet compared to replicate experiments, we investigated the known Zfp281 and Lefty1 enhancers as done before for ChIP-nexus data. Since the model evaluation was performed in peak regions, we added data augmentation (genomic intervals jittered uniformly by [-400, 400] bp with random reverse complementation) to make sure the model does not simply predict the average ChIP-seq signal centered at the peak. We observed that the predicted profile shapes significantly de-noise the base-resolution, strand-specific 5’-end coverage ChIP-seq profiles. Indeed, the predicted profiles resemble smoothed versions of the ChIP-seq 5’-end coverage profiles (averaging sliding window of 50bp, Figure S11A).

To evaluate the predictive performance of the ChIP-seq BPNet model, we performed a similar analysis as for ChIP-nexus with the difference that we assessed the quality of profile shape prediction by comparing the similarity of BPNet predictions to the ground truths smoothed ChIP-seq profiles on the test set against the similarity of ChIP-seq smoothed profiles between replicate experiments using the multinomial log-likelihood. We found that BPNet outperformed the smoothed replicate experiments in terms of profile shape prediction on almost all TFs except Nanog where both performed similarly (Figure S11B). Consistent with the ChIP-nexus models, the total count predictions from the BPNet ChIP-seq model did not surpass the concordance between replicate experiments (Figure S11C). As previously discussed, this result is expected since the total counts are likely influenced by factors besides local DNA sequence that we do not model, such as chromatin context and distal interactions with other genomic elements. Altogether, we conclude that BPNet generalizes seamlessly to learn accurate models of TF ChIP-seq profiles.

Next, we investigated the DeepLIFT profile contribution scores inferred from the ChIP-seq BPNet model for all three TFs in the well-known Oct4 enhancer. The contribution scores were computed in the exact same manner as for the ChIP-nexus model. Consistent with the inferences from the ChIP-nexus model, we found that the contribution scores also precisely highlighted the *Oct4-Sox2* motif in the center and the Nanog motif on the immediate flanks (Figure S11D). Hence, the contribution scores derived from a ChIP-seq BPNet model is able to accurately highlight the expected motifs within a well-known enhancer.

We then investigated the globally predictive motifs learned by the ChIP-seq BPNet model. We used TF-MoDISco with the same hyper-parameters as for the ChIP-nexus model. To allow for unbiased comparison of the motifs obtain from ChIP-seq and ChIP-nexus BPNet models, we retrained additional models on ChIP-nexus and ChIP-seq data using a common set of peak regions that were found by either of the two assays i.e. union of ChIP-nexus and ChIP-seq peaks, for the three TFs (Oct4, Sox2, Nanog). We further restricted model interpretation to a common set of peak regions found by both assays i.e. intersection of ChIP-nexus and ChIP-seq peaks for each TF.

Additionally, to evaluate the benefits of a profile regression model for ChIP-seq, we trained a binary classification model on ChIP-seq data in the same manner as done before for ChIP-nexus data.

We observed that TF-MoDISco applied to all the different types of ChIP-seq BPNet models discovered the majority of the expected motifs. However, the ChIP-seq models trained on binary labels found a few additional motifs that appear to be spurious and lacking clear biological significance (Figure S11E, Supplemental Table S2). These results show that ChIP-seq BPNet profile models perform comparably to ChIP-nexus BPNet profile models in terms of motif discovery with fewer spurious discovered motifs compared to models trained on binary labels.

To evaluate the quality of the CWM motif instances obtained by the four models, we use the same approach as in Figure S9B and Figure S10D in which we compare the ranked motif instances against co-localized strong ChIP-nexus footprints for each TF. We observed that ChIP-nexus BPNet profile models recalled a higher fraction of motif instances with strong ChIP-nexus footprints for the *Nanog* motif compared to ChIP-seq BPNet models (Figure S11F). Both models performed similarly well for *Oct4-Sox2* and *Sox2* motifs. Additionally, ChIP-seq BPNet profile models yielded more accurate CWM motif instances than ChIP-seq binary classification models trained on the same data as well PWM based instance calling (Figure S11G). In conclusion, with respect to accuracy of motif instances, the ChIP-nexus BPNet profile models outperform ChIP-seq BPNet profile models, which outperform ChIP-seq binary models. Hence, modeling profiles directly and improved resolution of the ChIP-nexus profiles collectively contribute to improved motif instance identification.

Altogether, these results show that the BPNet framework, which includes BPNet training, inference of DeepLIFT contribution scores, CWM motif discovery with TF-MoDISco, and motif instance identification via CWM scanning, can be readily applied to ChIP-seq data. These results were obtained with very minor hyper-parameter adjustments while explicitly controlling for assay specific biases. It should be possible to easily adapt and apply the BPNet workflow to any other regulatory profiling assays such as CUT&RUN, ATAC-seq and DNase-seq.

### Relationship between the Poisson log-likelihood, mean-squared error and multinomial log likelihood

We start by writing down the negative log-likelihood for the Multinomial distribution. Let *L* be the sequence length, *N* the total number of events (i.e. total number of read counts in the region) and *p*_*i*_ the probability of obtaining the outcome *i* (e.g. the read gets aligned to position *i*). Then, the negative log likelihood can be written as

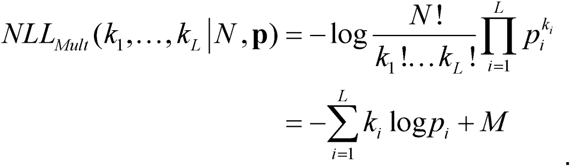

Note that we gathered all the terms independent of *p*_*i*_ ∀*i* into the constant *M*. Let’s assume the read counts at each genomic location *k*_*i*_ are distributed according to the Poisson distribution. The Poisson log likelihood for the sequence region of length *L* can be written as

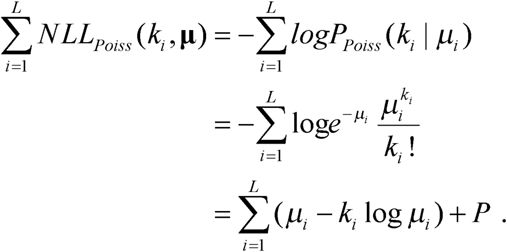

If we replace *μ*_*i*_ with *N*_*p*_*p*_*i*_, where *N*_*p*_ is the predicted number of total counts and use 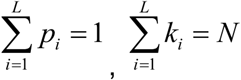, we obtain:

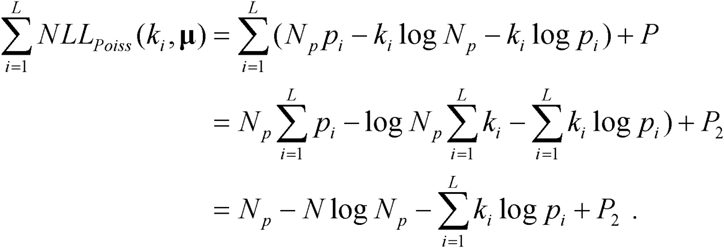

We observe that the second term equals to the multinomial negative log-likelihood. If we set 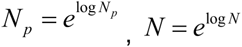, and perform a Taylor expansion

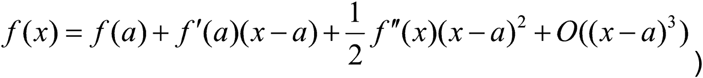

up to the squared term for variable log*N*_*p*_ around log*N* using, we obtain:

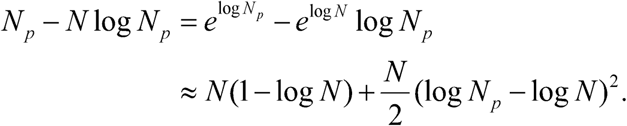

This means that we can approximate the Poisson log-likelihood by a sum of mean-squared errors and the multinomial loss function where the predicted log of total counts log*N*_*p*_ is close to the true total counts log*N*:

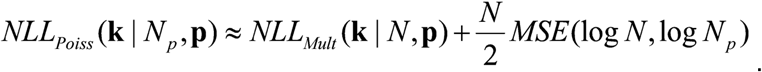

We approximate the expression further by replacing the *N* in front of MSE with 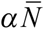, where 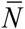 is the average (or median) value of *N* across the dataset and *α* is the tuning parameter which allows to up or down-weight the importance of total count prediction:

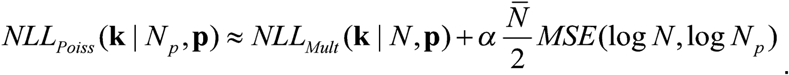

If *α* = 1, the multinomial loss and the mean squared error loss are balanced according to the Poisson log-likelihood.

## Supplemental figures

**Figure S1:**
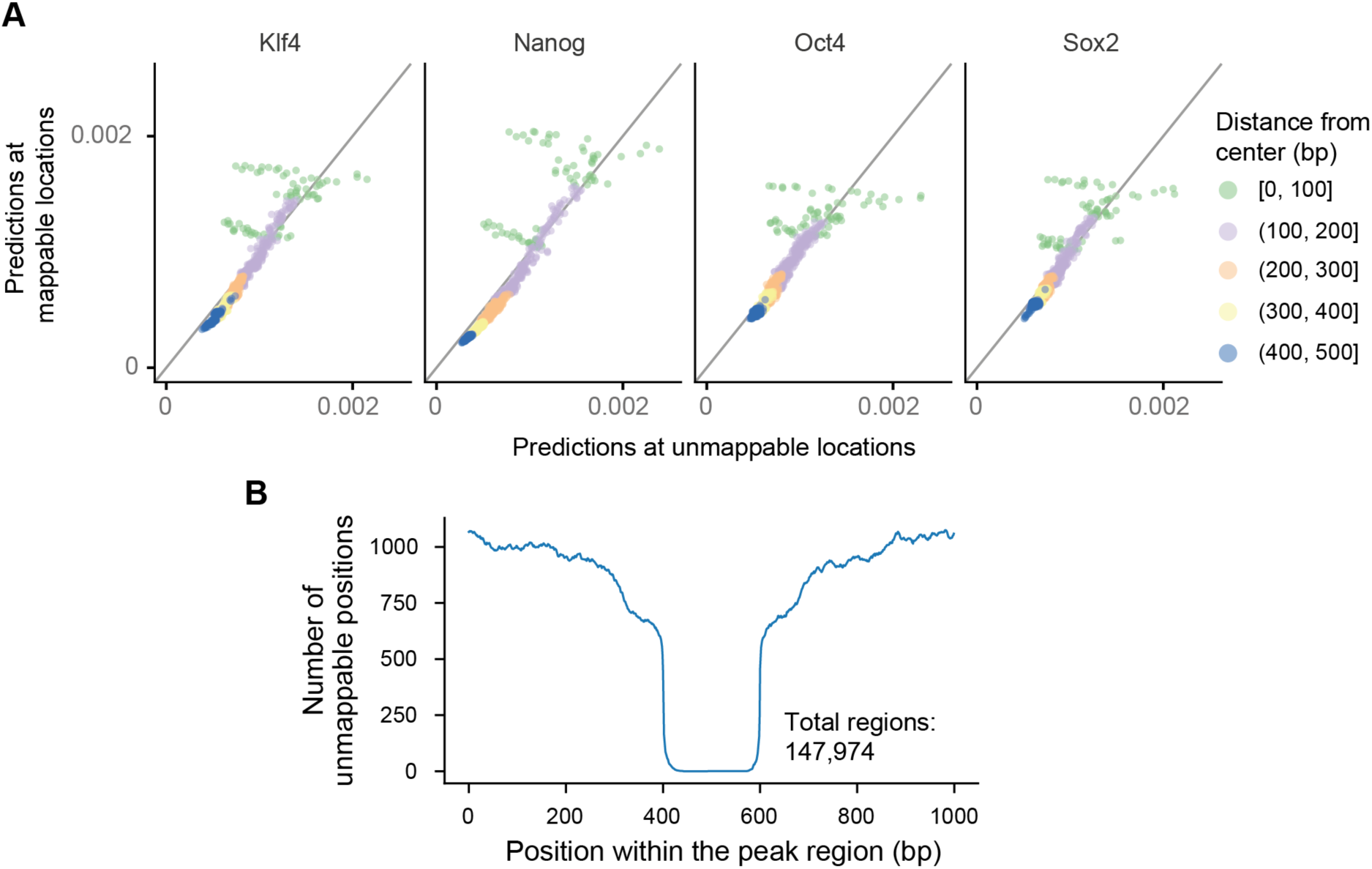
Effect of mappability on BPNet predictions. **Model predictions exhibit no systematic overfitting to unmappable positions A)** Median model predictions for the positive strand at unmappable (x-axis) and randomly chosen mappable positions (y-axis) stratified by distance from the peak summit (denoted by points and color). Each of the thousand points corresponds to a specific relative position within the 1 kb peak and the color highlights different subregions within the peak regions. Points on the diagonal mean that model prediction at unmappable positions (x-axis) is not systematically different from the mappable positions (y-axis). None of the unmappable positions are predicted to have 0 probability of reads indicating that the model is not overfitting to the false 0 counts at these positions. Points with [0, 100] distance from the center are more scattered since the median is computed over very few points for unmappable positions and because the strongest signal associated with footprints (and hence highest variance) is also observed in these regions (as shown in B). **B)** Number of unmappable positions for different positions within the 1000 bp peak region. Overall, 0.5 % of the positions were unmappable.

**Figure S2:**
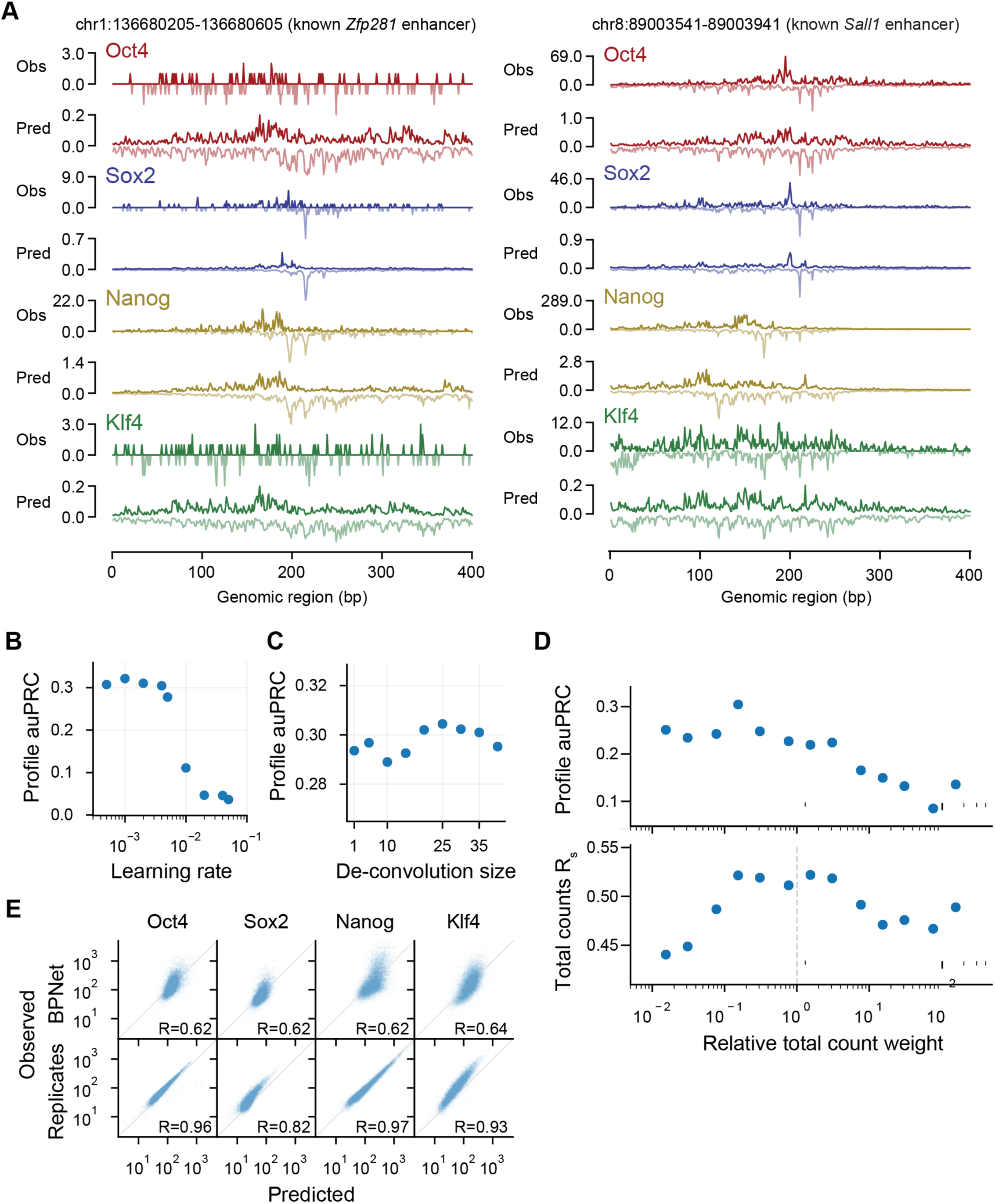
Performance evaluation of ChIP-nexus BPNet profile models. **Additional predictive performance evaluation for BPNet. A)** Observed and predicted ChIP-nexus read counts mapping to the forward strand (dark) and the reverse strand (light) for the *Zfp281* and *Sall1* enhancers located on the held-out (test) chromosome 1. **B)** auPRC of profile predictions is high across various learning rates on the tuning set chromosomes 2-4 demonstrating the robustness of the model. **C)** The deconvolutional layer slightly improves the profile predictive performance compared to a point-wise convolutional layer (deconvolution size=1). **D)** auPRC of profile predictions (top) and the Spearman correlation of total count predictions (bottom) for a range of different relative total count weight α in the BPNet loss function parameterized as λ = α/2 n_obs. Relative weight of 1 (center) denotes equal weighting of the counts and profile loss functions. The best performance is obtained for alpha < 1 showing that putting more weight to profile predictions helps for both profile and count predictions. **E)** Observed and predicted total read counts for BPNet (top) and replicate experiments (bottom) across the four studied TFs along with the Spearman correlation coefficient.

**Figure S3:**
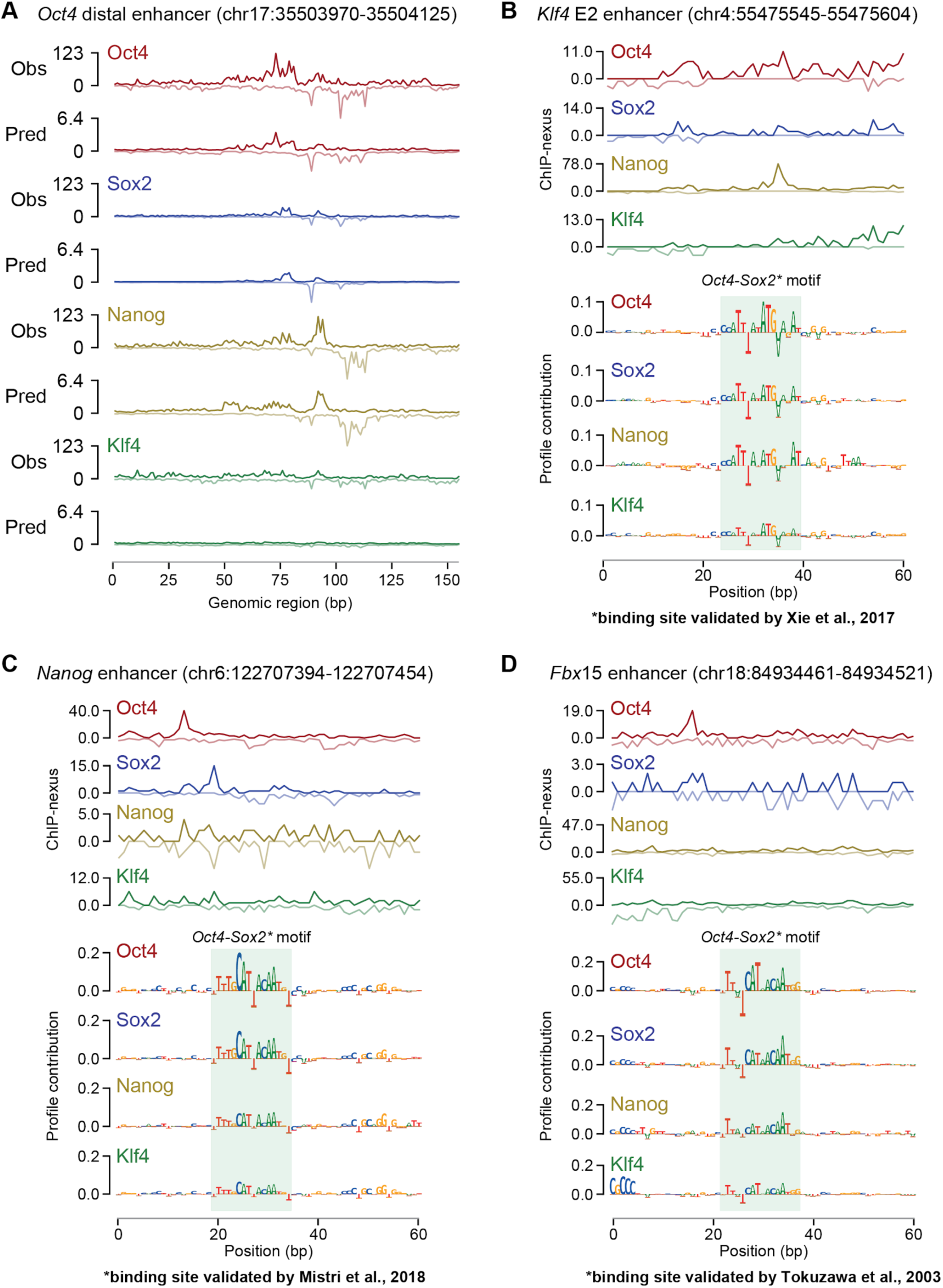
BPNet predictions and sequence contribution scores at known enhancers. **Additional BPNet predictions across known enhancer regions. A)** Observed and predicted ChIP-nexus read counts for the *Oct4* distal enhancer. **B**,**C**,**D)** Previously validated binding motifs for *Oct4-Sox2* were re-discovered by BPNet. ChIP-nexus read counts and BPNet contribution scores for three enhancers are shown. **B)** The *Oct4-Sox2* motif site in the *Klf4* E2 enhancer was validated by deleting the site using CRISPR/Cas9 ^73^. **C**,**D)** The *Oct4-Sox2* binding motifs in the *Nanog* and *Fbx15* enhancers were confirmed previously using reporter assays of constructs with various motif mutations ^74,75^.

**Figure S4:**
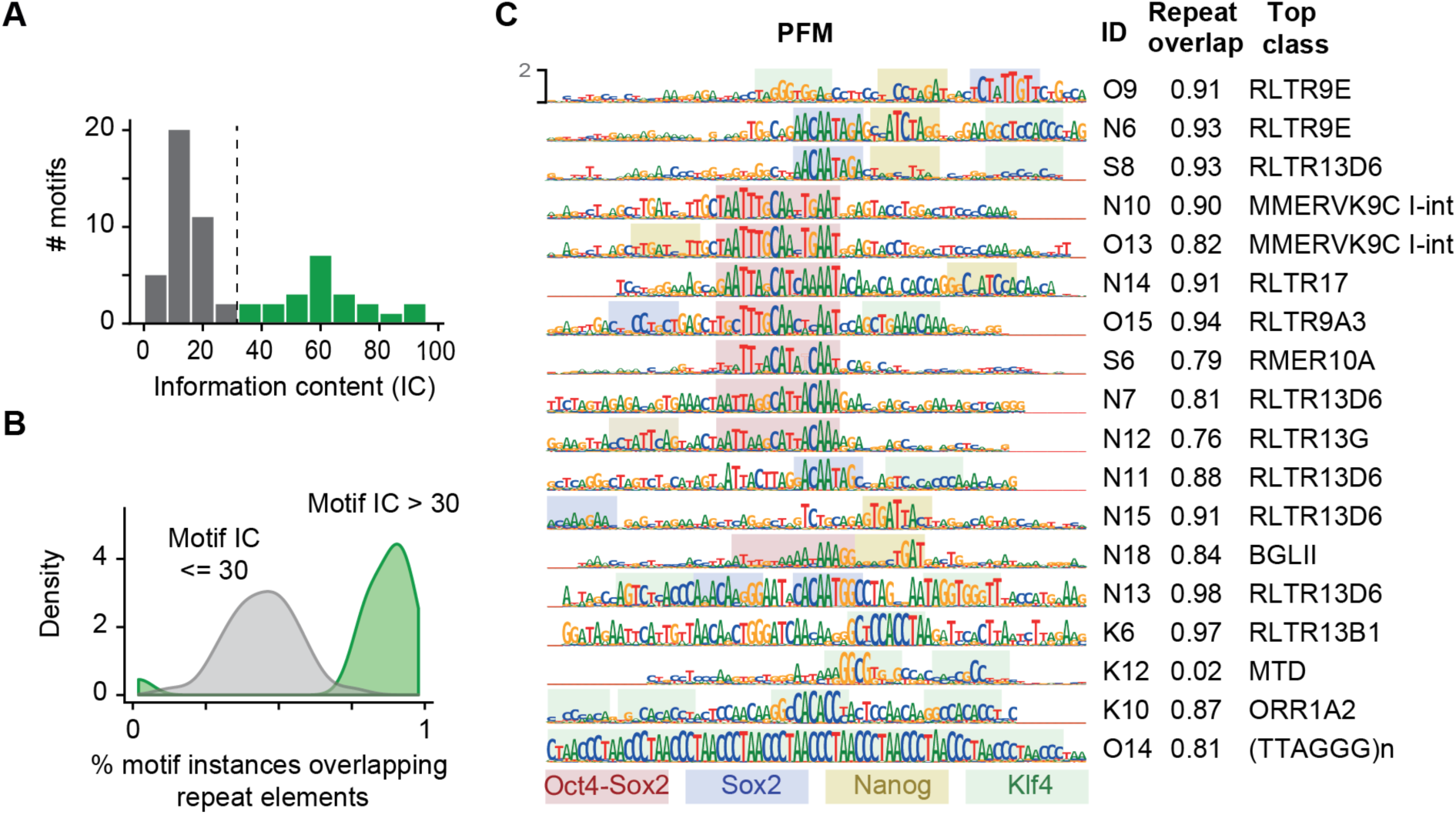
Long motifs discovered by TF-MoDISco come from retrotransposons. **TF-MoDISco discovered long motifs that originated from retrotransposons. A)** Among all motifs discovered by TF-MoDISco, 18 motifs display unusually high information content (IC) of >30 bits (green). The expected short motifs are shown in grey. **B)** Histogram of the overlap of short motifs (grey) and long motifs (green) with repeat elements shows that long motifs overlap >80% with annotated retrotransposons. **C)** Long motifs with their PFM, ID, fraction of motif instances overlapping with a repeat, and the most frequent (top class) RepeatMasker annotation. Highlighted within the repeat elements are potential motif instances of *Oct4-Sox2, Sox2, Nanog* and *Klf4* as indicated by the CWMs.

**Figure S5:**
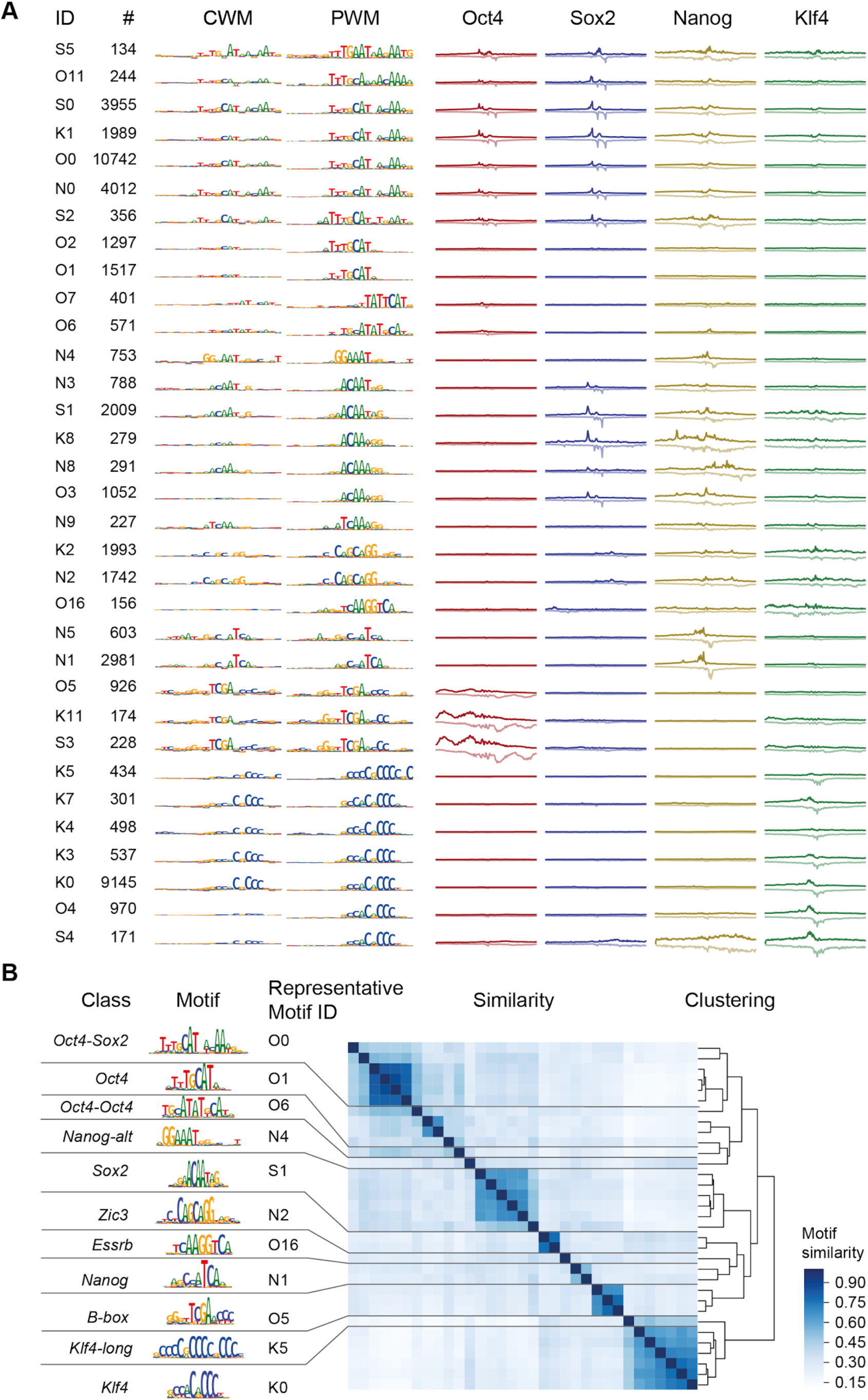
TF-MoDISco motif overview. **Overview and clustering of all short motifs discovered by TF-MoDISco. A)** All 33 discovered short motifs (information content < 30 bit) are shown with: (from left to right) motif ID, number of seqlets supporting the motif, CWM, PFM, and average ChIP-nexus read count distribution (180 bp) for each TF. All sequence logos and profile plots share the same y-axis in each column. Motif ID consists of the TF name for which the motif was discovered (O for Oct4, S for Sox2, N for Nanog, and K for Klf4) and the order (starting with 0) in which the motif was discovered by the TF-MoDISco run for the TF. **B)** To remove redundant motifs (e.g. discovered through different TFs) and identify a set of representative motifs for the downstream analysis, motifs were clustered by similarity using hierarchical clustering. The results were then manually inspected to select clusters that separate known motifs that are distinct (e.g. *Oct4-Oct4* resembles the known MORE and PORE motifs that bind Oct4 homodimers, which is different from the monomerically bound *Oct4* motif). Among very similar motifs within a cluster, we then selected the most abundant motif that was discovered for the most relevant TF (if known). The 11 representative motifs that we selected are shown on the left. Non-canonical motifs were given a name (*Nanog-alt* for Nanog alternative, *Klf4-long* for longer *Klf4*).

**Figure S6:**
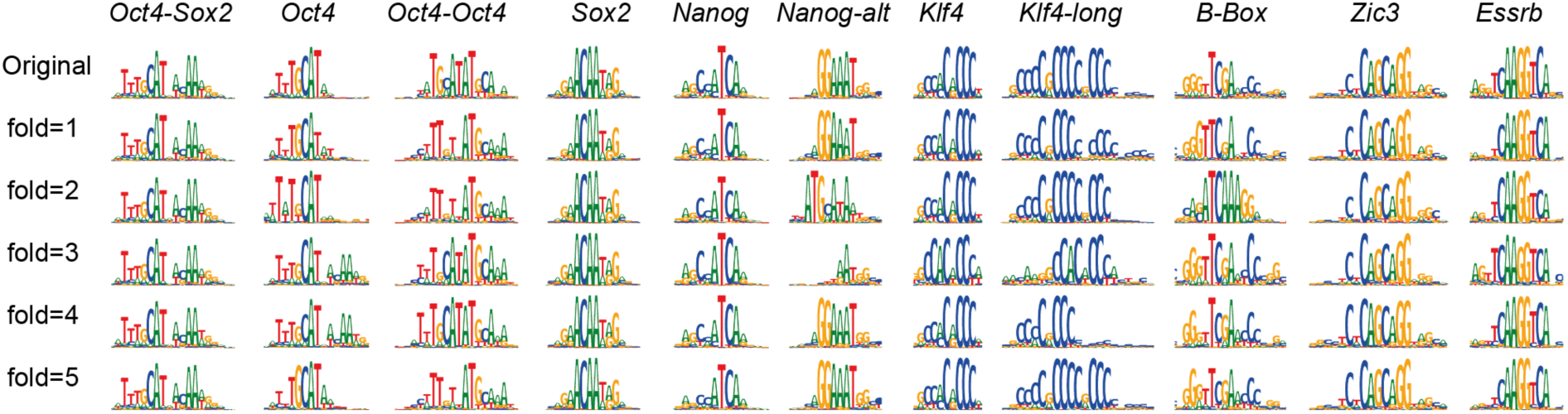
Validation of the motifs by training BPNet on five folds. **BPNet trained on different chromosome sets (folds) yields similar motifs.** The closest motif match from each BPNet/TF-MoDISCo cross-validation run (Methods) resembles the originally discovered motifs.

**Figure S7:**
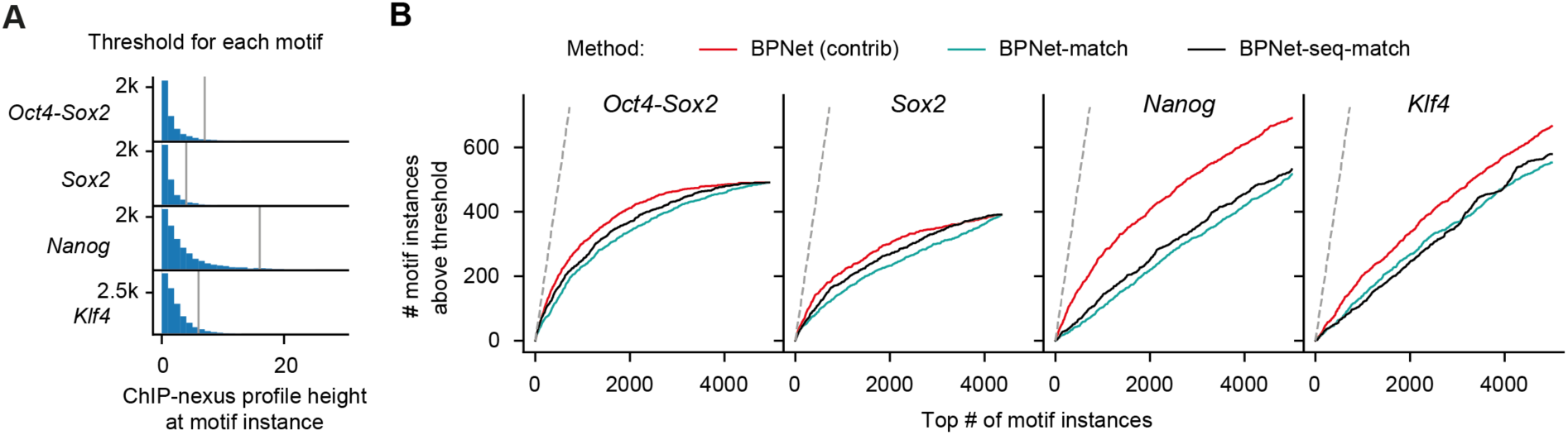
Performance of BPNet contribution scores versus match scores. **Ranking motif instances by BPNet contribution scores recovers more motifs supported by ChIP-nexus footprints than ranking by match-based scores (sequence or contribution). A)** ChIP-nexus profile height distribution at the reference summit position for BPNet motif instances of different TFs (Methods). The vertical grey lines denote the 90th percentile that is used as a stringent threshold for calling motif instances as having a ChIP-nexus footprint. **B)** Number of top-ranked N motif instances located up to 500 bp away from the ChIP-nexus peak summits showing a ChIP-nexus footprint larger than the threshold defined in Figure S7A. The top N motif instances (x-axis) were ranked either by the contribution score magnitude at the motif region (BPNet), match score between the CWM and the contribution scores (BPNet-match), or the match score between the PWM and the sequence (BPNet-seq match). Note that all motifs in this analysis were originally called by BPNet (using both the contribution score magnitude and match), hence motifs with a good sequence match and poor contribution scores (or good contribution score and poor sequence match) were already excluded during the original instance calling.

**Figure S8.**
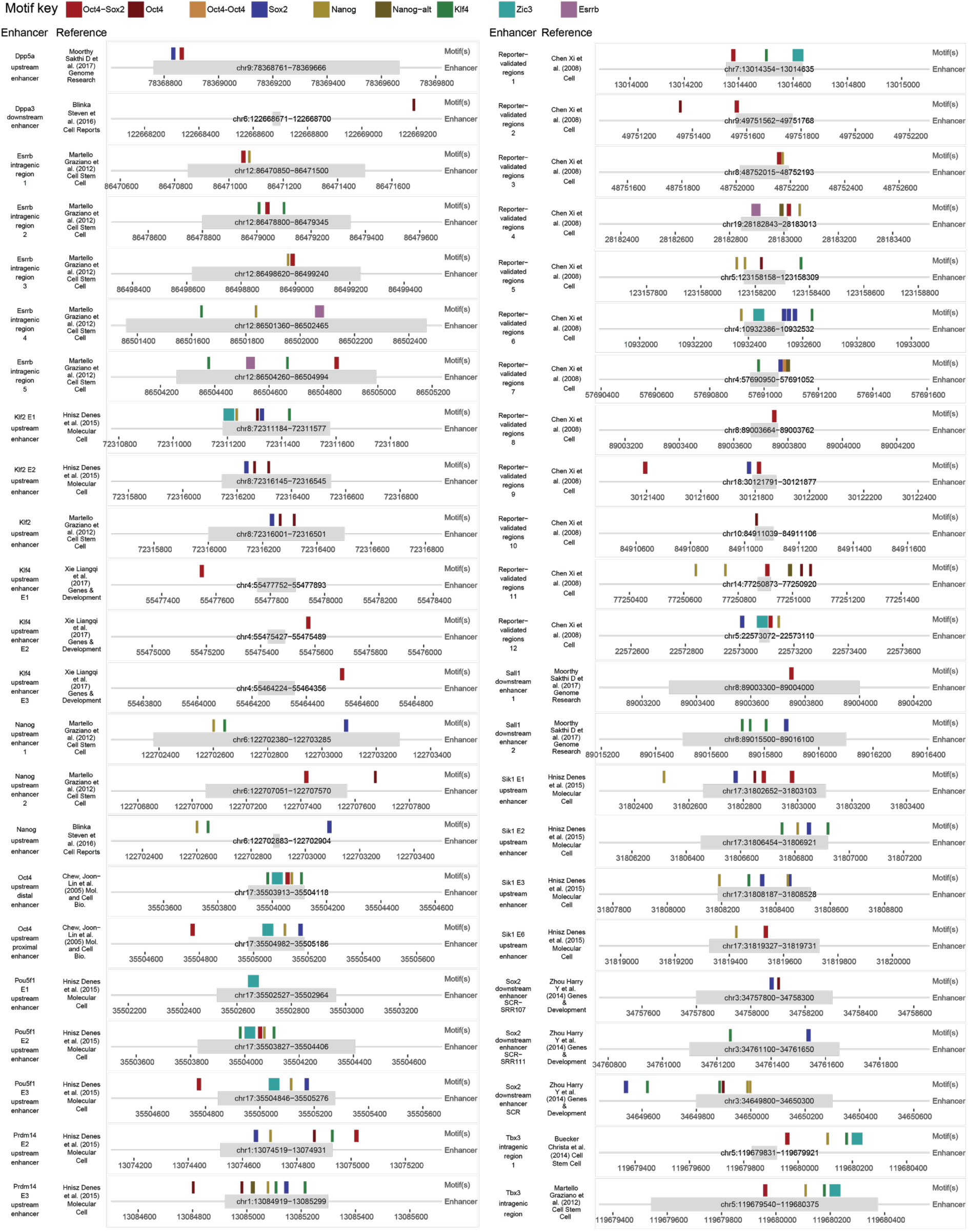
Mapped motif instances across previously identified enhancer regions. The enhancer coordinates (gray boxes) are based on previous publications ^63,67,73,152–156^ and do not necessarily imply that the region is sufficient for enhancer activity. Mapped motif instances (colored boxes) are shown in the surrounding region.

**Figure S9.**
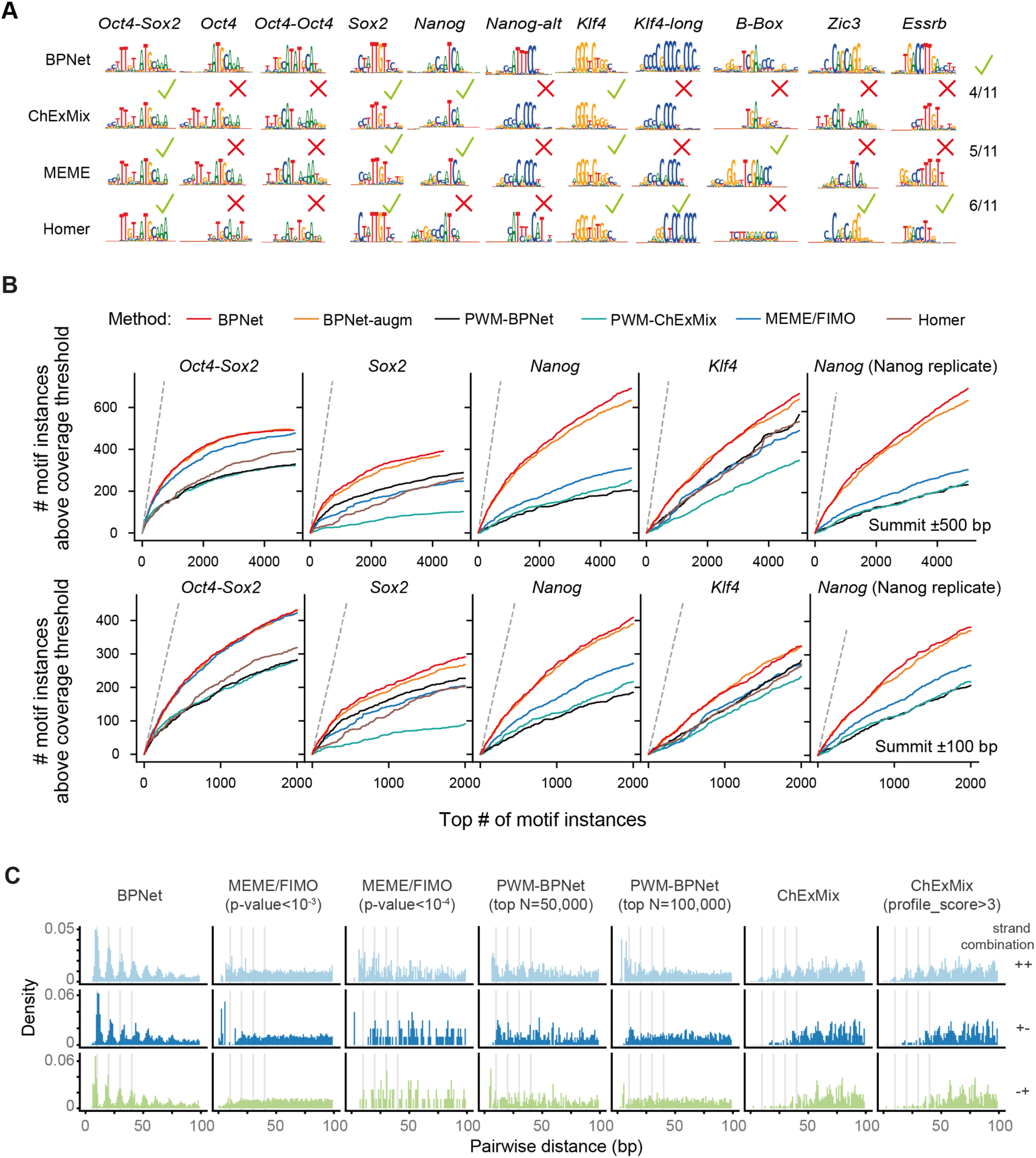
BPNet and TF-MoDISco discover more motifs than ChExMix, MEME or HOMER as well as map motif instances with greater accuracy than PWM scanning. **A)** Motifs discovered by ChExMix, HOMER, and MEME for Oct4, Sox2, Nanog and Klf4 ChIP-nexus peaks that are closest to the 11 primary representative BPNet motifs (top row). Green checkmark denotes whether the discovered motif is similar to the BPNet motif. **B)** Number of motif instances located up to 500 bp (top) or 100 bp (bottom) away from the ChIP-nexus peak summits showing a strong ChIP-nexus footprint. Only motif instances in peaks from held-out test chromosomes (1, 8 and 9) were used for the evaluation. (x-axis) top N motif instances from each of the methods were sorted in descending order of scores (PWM log odds score or CWM contrib score). For BPNet-augm, the center of the genomic region for which the contribution scores were computed was randomly jittered up to 200 bp away from the peak summit. This augmentation prevents BPNet from using the positional information of the peak summit. In the final column (Nanog replicate), the Nanog ChIP-nexus footprint was measured by a separate biological replicate using a different antibody (ɑ-Nanog from Abcam, ab214549), which was not used during training or evaluation. **C)** The pairwise spacing of Nanog motif instances located up to 100 bp away from the ChIP-nexus peak summits in all possible strand orientations (rows) for different methods and/or thresholds (columns). Results for all chromosomes are shown.

**Figure S10.**
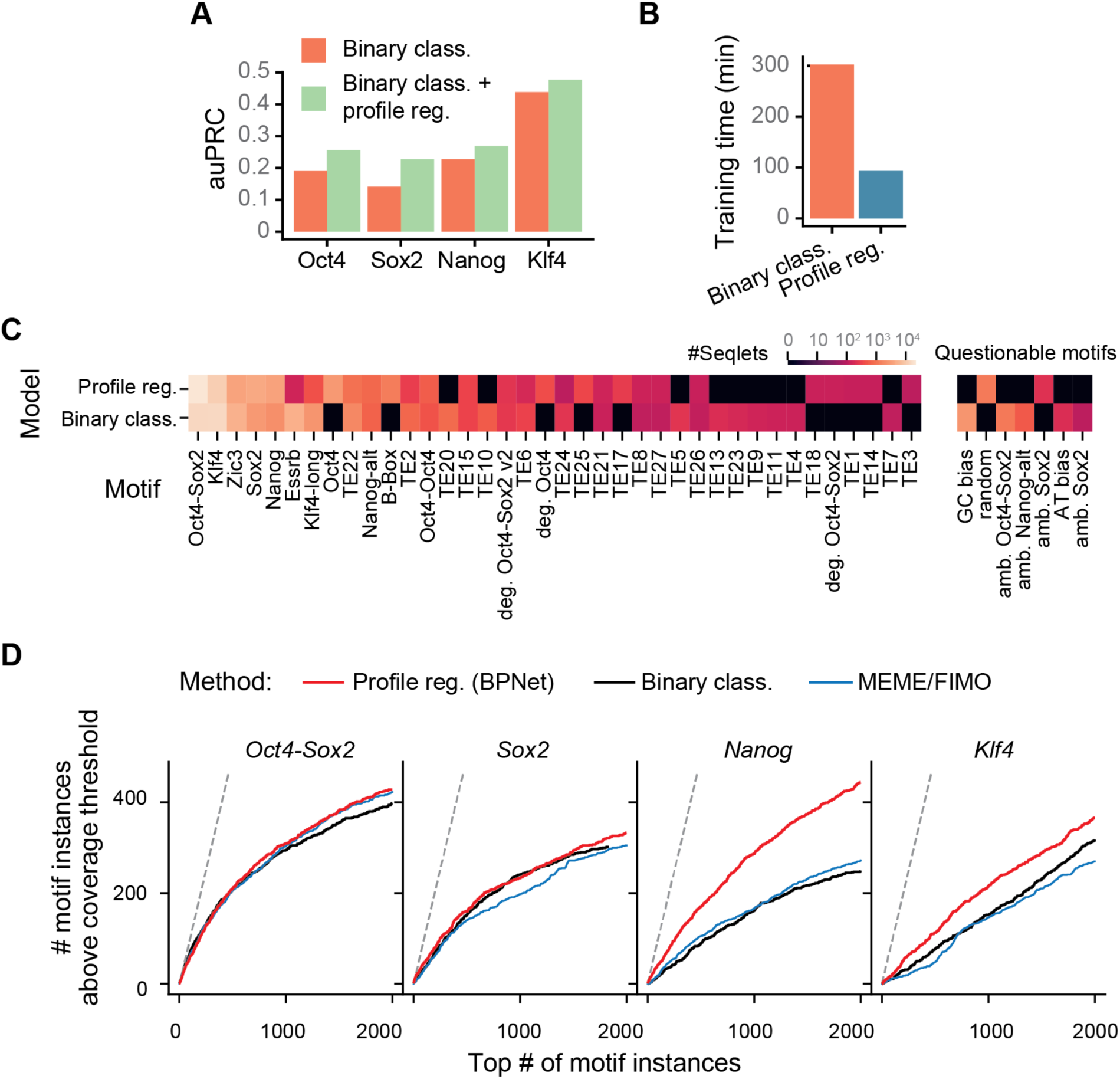
BPNet trained to predict the ChIP-nexus profile is faster and yields more accurate motif instances than a binary classification model. **A)** Predictive performance of the binary classification models predicting the presence or absence of ChIP-nexus peaks from 1 kb DNA sequences evaluated across the held-out (tuning) chromosomes 2, 3, and 4. The model trained to classify the sequences is shown in orange and the model trained to also predict the ChIP-nexus profiles from DNA sequence in addition to classifying them is shown in blue. **B)** Training time of the binary classification model trained genome-wide and the sequence-to-profile model (BPNet) trained in ChIP-nexus peaks. **C)** Detected motifs by TF-MoDISco using the contribution scores in ChIP-nexus peaks of the sequence-to-profile BPNet (profile reg.) or the binary classification model (binary class). A light color denotes a high number of seqlets for each motif. Motifs not discovered or motifs supported by less than 100 seqlets are shown in black. Questionable motifs are displayed separately on the right. **D)** The number of motif instances (500 bp within ChIP-nexus peak summit) showing a ChIP-nexus footprint (y-axis) within the top N motif instances with highest contribution scores (x-axis) from the held-out (test) chromosomes 1, 8 and 9. A site was considered to show a ChIP-nexus footprint if the number of reads at the position of the aggregate footprint summit (averaged across both strands) is higher than the 90^th^ percentile value of all motif instances detected by the profile regression model for the corresponding TF (i.e. same as in Figure S9B).

**Figure S11.**
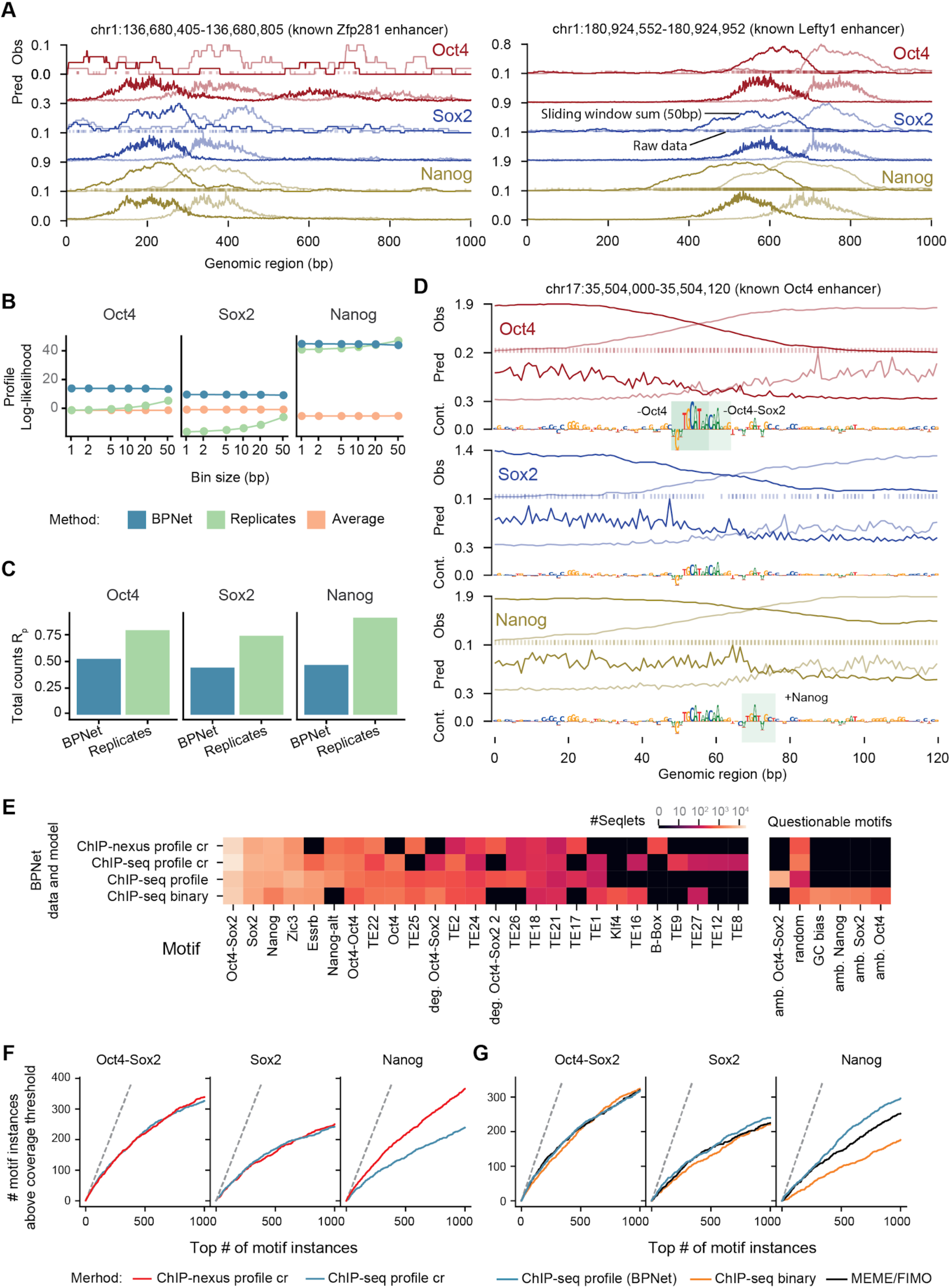
The BPNet framework can be used to model and interpret TF ChIP-seq data. **A)** Observed and predicted read counts for BPNet trained on ChIP-seq data for the *Zfp281* and *Lefty1* enhancers located on the held-out (test) chromosome 1. Reads mapping to the forward strand are displayed in dark and reads mapping to the reverse strand in light. For the observed read counts, a sliding window of 50 bp was used to smooth the raw 5’ end read counts (line). Raw counts are shown as points on the bottom at y=0. **B)** BPNet predicts the ChIP-seq profile shape better than the replicates. Multinomial-log likelihood given the observed number of total counts was used to evaluate the profile shape quality at different resolutions (from 1 bp to 10 bp windows) in held-out chromosomes 1, 8 and 9 (Methods). A log-likelihood of 0 corresponds to the constant model. **C)** Total counts in the 1kb regions centered at the peak summits in the region can be predicted (blue) at a decent accuracy level as measured by Spearman correlation but do not surpass replicate performance (green). **D)** Observed and predicted read counts as well as the contribution scores of BPNet for the known Oct4 enhancer. As for A, the observed read counts are shown both as smoothed (line) and as raw counts (points at y=0). Motif instances derived by CWM scanning are highlighted with a green box. **E)** BPNet applied to ChIP-seq discovers the majority of the motifs identified by BPNet applied to ChIP-nexus data. The models ‘ChIP-nexus profile cr’ and ‘ChIP-seq profile cr’ were trained on the union of the ChIP-nexus/seq peaks predicting Oct4, Sox2, and Nanog binding and were interpreted on the intersection of the ChIP-nexus/seq peaks. **F**) Motif instance calling with CWM scanning has higher accuracy for BPNet trained on ChIP-nexus data than for BPNet trained on ChIP-seq data (evaluated on the union of the ChIP-nexus/seq peaks, 500 bp around the peak summit using ChIP-nexus footprints as ground truth). **G)** Training a sequence-to-profile model on ChIP-seq data yields more accurate motif instances (500 bp around the ChIP-seq peak summits using ChIP-nexus footprints as ground truth) than training a binary classification model or using a PWM scanning approach using FIMO for motifs derived directly from ChIP-nexus data (Figure S9). See Figure S10D and S9B legends for a detailed description.

**Figure S12.**
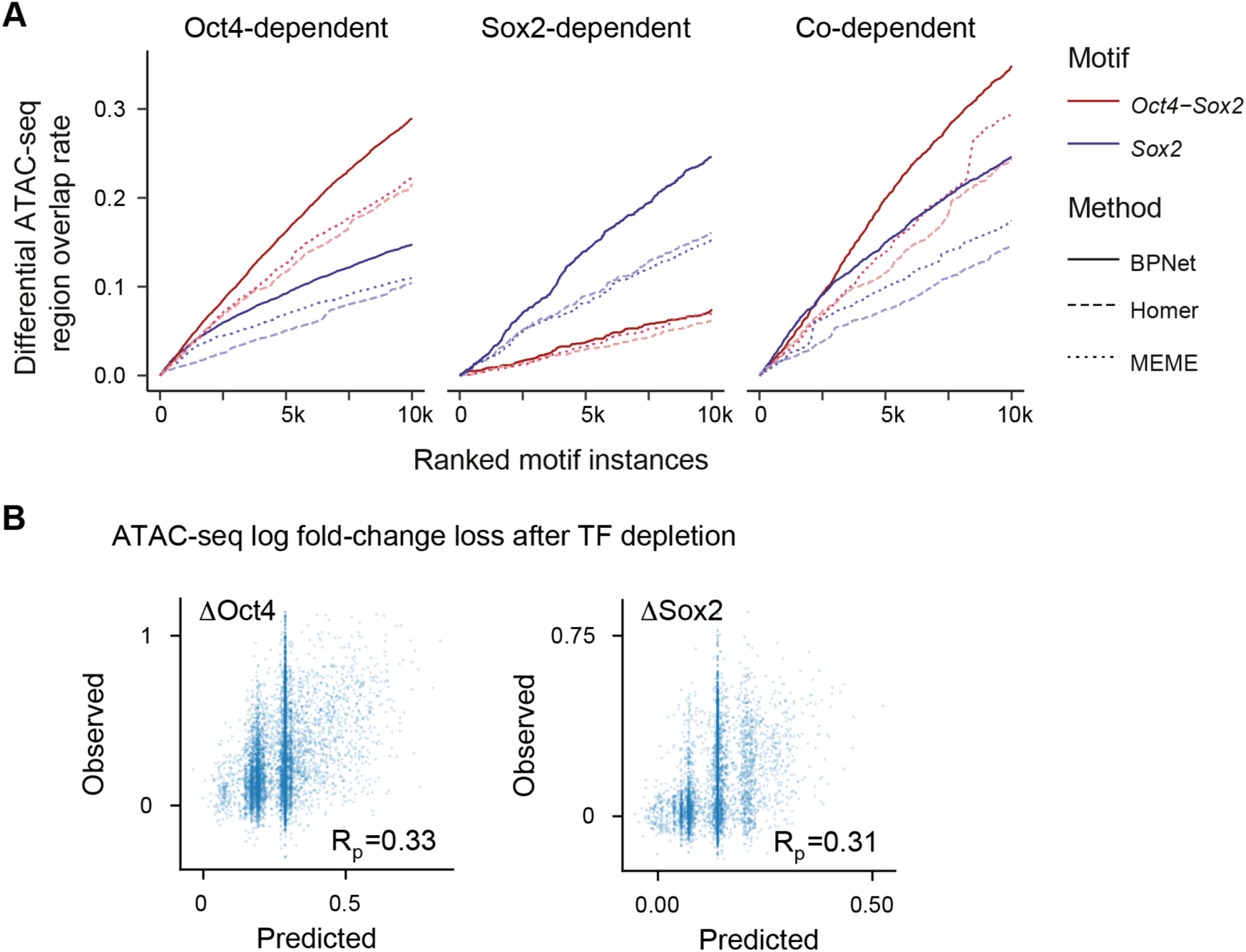
Motif validation using ATAC-seq data. **A)** Overlap of ranked *Oct4-Sox2* and *Sox2* motif instances (in thousands *k*) with regions that lose ATAC-seq signal in response to either Oct4 or Sox2 depletion as defined by ^77^. In addition to the data shown in Figure 2G, the results for co-dependent regions are shown, in which ATAC-seq signal is lost in response to either Oct4 or Sox2 depletion. Both *Oct4-Sox2* and *Sox2* motifs are present. Motif instances ranked by BPNet contribution scores also outperformed those obtained by HOMER and MEME (ranked by PWM match scores). **B)** Linear regression model based on motif instance features, rather than the BPNet bottleneck layer (Figure 2H), of the ATAC-seq log fold-change between Oct4 ON and OFF (*Δ*Oct4, left) or Sox2 ON and OFF (*Δ*Sox2, right). Note that the Pearson correlation (*RP*) coefficient for both motif instance based models is only around half of that of the models based on the bottleneck layer. The motif instance features were the number of BPNet motif instances located in the 1 kb ChIP-nexus peaks and their average sequence match scores (Methods). Similar results were obtained for motif instances derived by MEME/FIMO, ChExMix, and HOMER. This result indicates that the mapped motif instances are strongly enriched in differentially accessible sites after TF depletion and contribute to the prediction of differential ATAC-seq signal. However, the sequence representation (bottleneck layer) learned by the BPNet model encodes additional information (such as motif syntax) beyond linear, additive effects of motif instances, thereby significantly improving prediction of differential accessibility.

**Figure S13:**
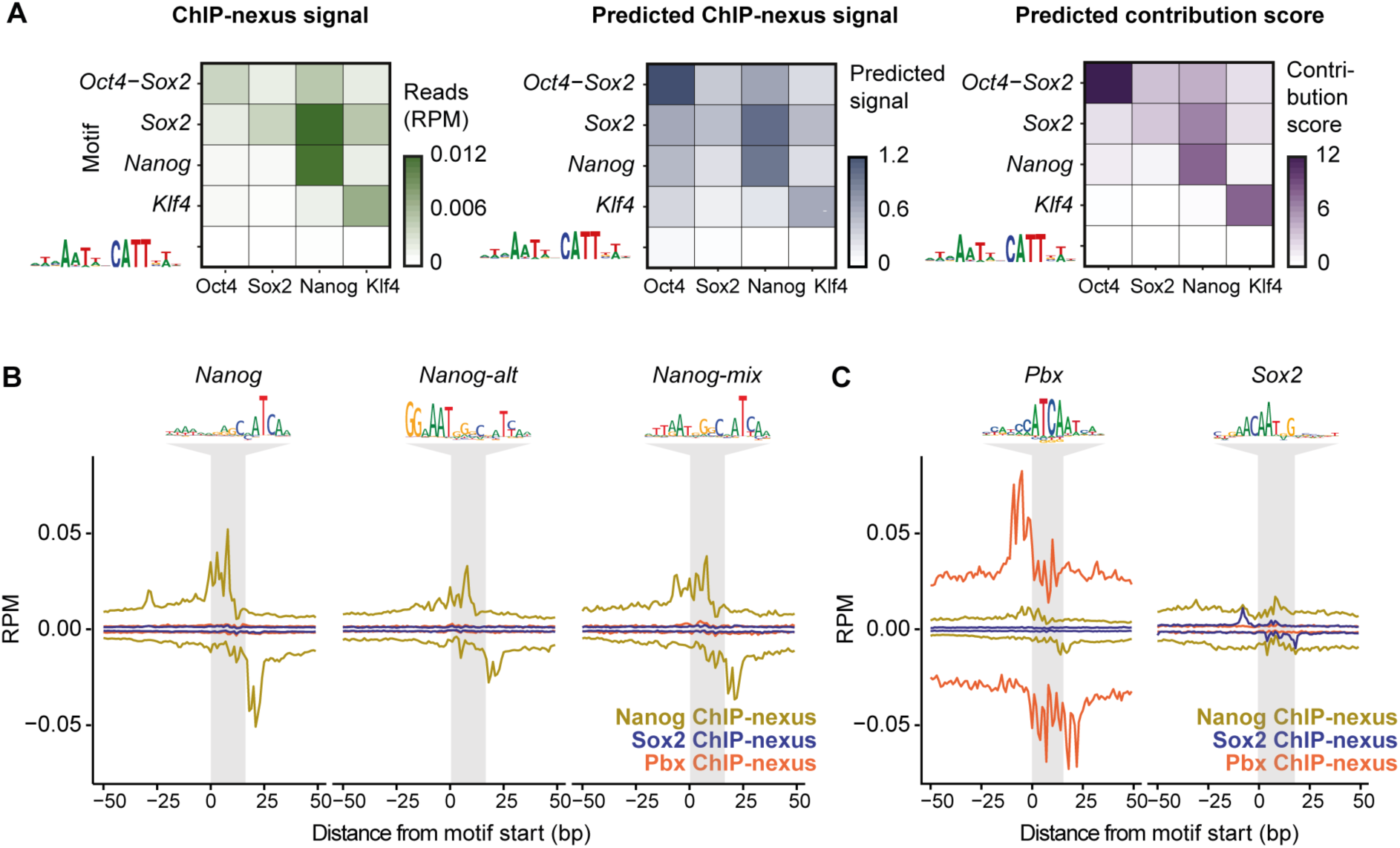
No evidence that Nanog binds with a partner. **Lack of evidence for a Nanog binding partner. A)** Median ChIP-nexus signal, predicted BPNet signal, and DeepLIFT contribution of Oct4, Sox2, Nanog, and Klf4 show no signal across the genomic instances matching the putative *Nanog-Sox* heterodimer motif (RMWMAATWNCATTSW) ^71^. The signal for *Oct4-Sox2, Sox2, Nanog*, and *Klf4* motif instances are shown as control. **B)** Since the *Nanog* motif resembles the known Pbx binding motif, we performed Pbx ChIP-nexus experiments to test whether Pbx might be a binding partner for Nanog. However, the average Nanog, Pbx and Sox2 ChIP-nexus binding profiles show no detectable footprints for Pbx or Sox2 on the three *Nanog* motifs, arguing against Pbx or Sox2 being stable interaction partners. However, an unknown interaction partner cannot be ruled out. **C)** The average Pbx ChIP-nexus footprint on the known *Pbx* motif from JASPAR ^142^ (top 1000 based on PWM-match score) confirms that the Pbx ChIP-nexus experiment worked (left). Likewise, Sox2 shows specific binding to its identified Sox2 motif (right). Note that the y-axis is set to the same RPM scale in A and B to allow comparisons of signal strength.

**Figure S14:**
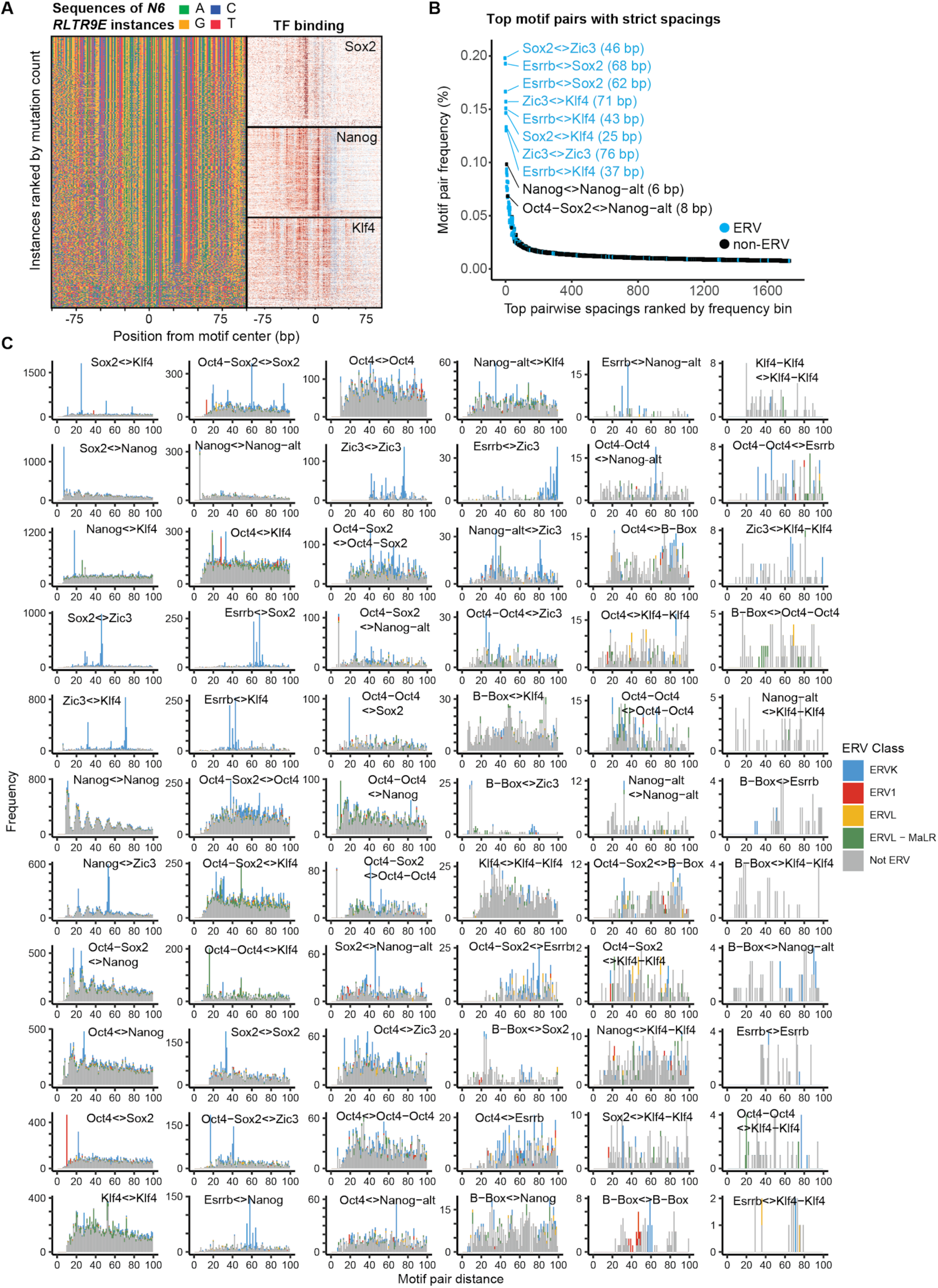
Strict spacings between motifs are likely due to retrotransposons. **Most over-represented instances of strict spacings between motifs are due to ERVs. A)** To show that TF binding occurs with strict spacings in retrotransposons and that this is likely ancestral, the *RLTR9E N6* motif is shown as an example. Sequences of the individual instances in the genome were sorted by the Kimura distance from the consensus motif, with the most similar sequences on top (which are likely more ancestral). Nanog, Sox2 and Klf4 ChIP-nexus binding footprints are shown in the same order on the right (+ strand reads in red, - strand reads in blue), revealing that the binding site spacing is largely constant across all sequences. **B)** Analysis of the most frequent distances between motif pairs (with >500 co-occurrences, distance measured at the trimmed motifs’ centers). The top 1% most frequent distances mapped in 83% to ERVs and were often larger than 20 bp. **C)** Histograms depicting the frequency of center-to-center motif pair spacings across the 11 representative motifs. Colors represent ERV classes which overlap with the corresponding motif pairs.

**Figure S15.**
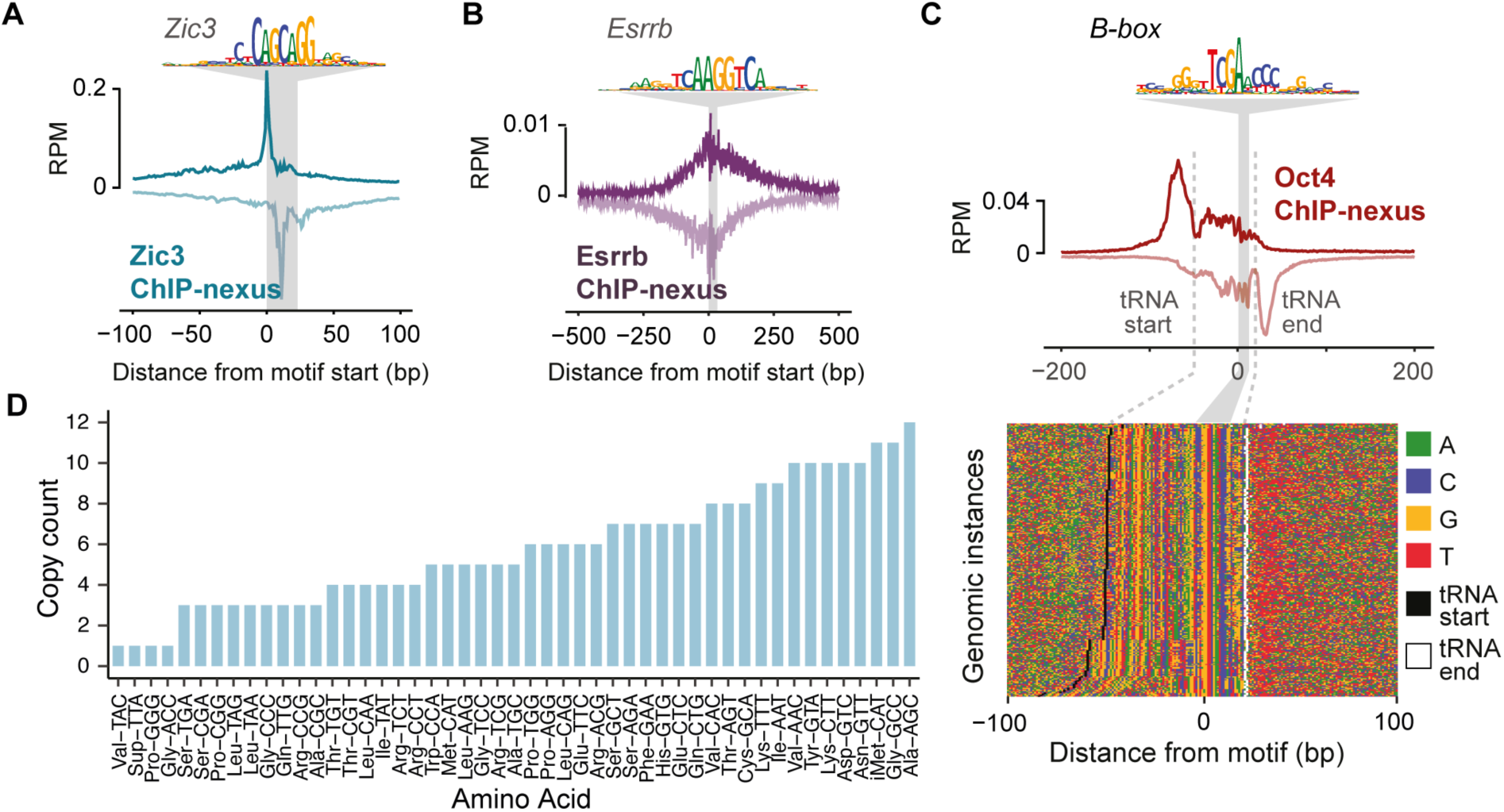
Validation of discovered motifs. **A)** To validate the identified *Zic3* motif instances, Zic3 ChIP-nexus experiments were performed. The average signal across the *Zic3* instances reveals a strong Zic3 binding footprint. **B)** A similar validation was performed for the *Esrrb* motif instances, revealing that the Esrrb ChIP-nexus signal is present but more diffuse at the discovered *Esrrb* motif instances. **C)** To better understand the binding of Oct4 to the *B-box*, which is frequently found in tRNA, tRNA-overlapping *B-box* motif instances were reoriented to match the transcriptional direction and sorted by tRNA gene start proximity. This reveals Oct4 binding at tRNA gene start/stop sites. **D)** Amino acid anti-codons and their copy count of the tRNAs that overlapped with the *B-box* motif instances.

**Figure S16:**
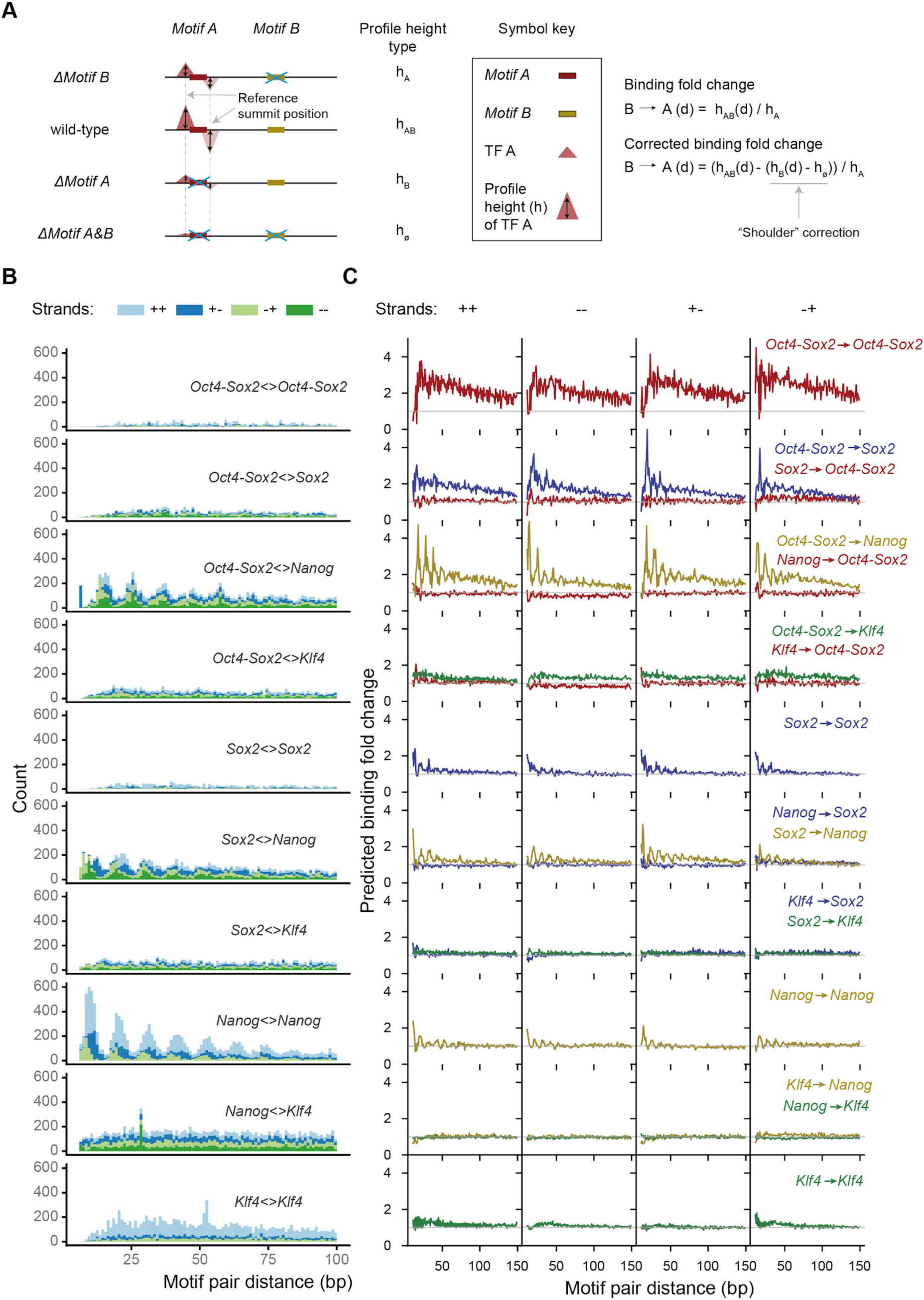
Pairwise in silico motif interactions for all strand orientations. **Analysis of all pairwise interactions between the four main motifs. A)** The influence of *Motif B* on the binding of TF A at *Motif A* is quantified by the fold change of predicted profile height at the reference summit position when *Motif B* is present or absent nearby (*h*_*AB*_ vs *h*_*A*_). The binding fold-change is corrected for the “shoulder” effect of *Motif B* by subtracting the predicted profile height when only *Motif B* is present in the sequence. **B)** Spacing distribution of all CWM-derived motif instance pairs in the genome stratified by motif identity and strand orientation. Note that for homotypic interactions, ++ and -- are the same and are shown as ++. **C)** *In silico* analysis of motif interactions on synthetic sequences measuring the predicted binding fold-change for all motif pairs across all strand orientations. Note that no clear differences between the possible strand orientations were detected.

**Figure S17:**
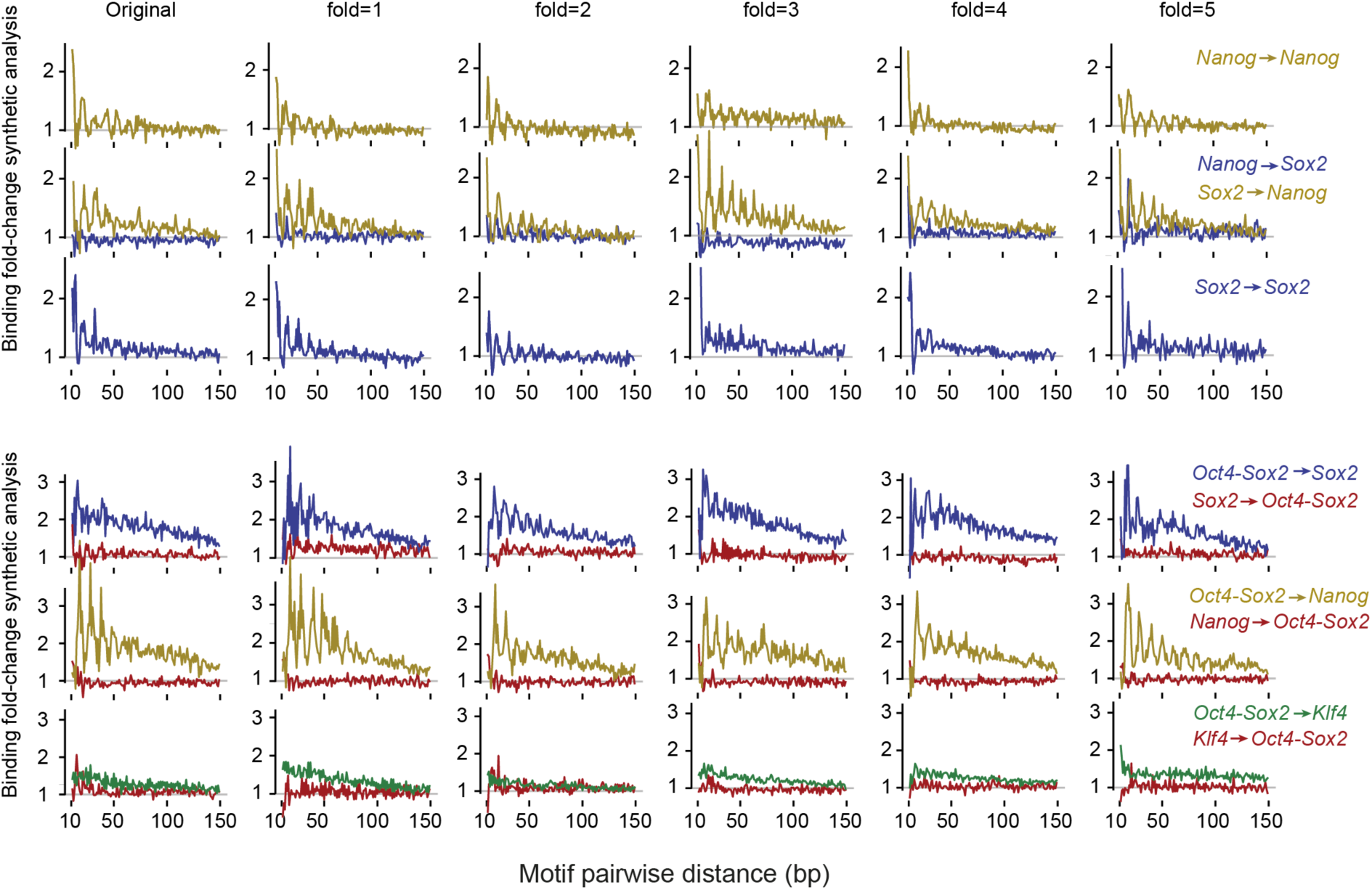
Validation of in silico motif interactions from BPNet model trained on five folds. **BPNet trained on different chromosome sets (folds) yields similar *in silico* interactions results.** *In silico* interaction analysis as shown in Figure 4C using BPNet trained on five different chromosome folds and using the motifs discovered by TF-MoDISCo for each fold (Methods). The observed interactions are highly reproducible across all folds.

**Figure S18.**
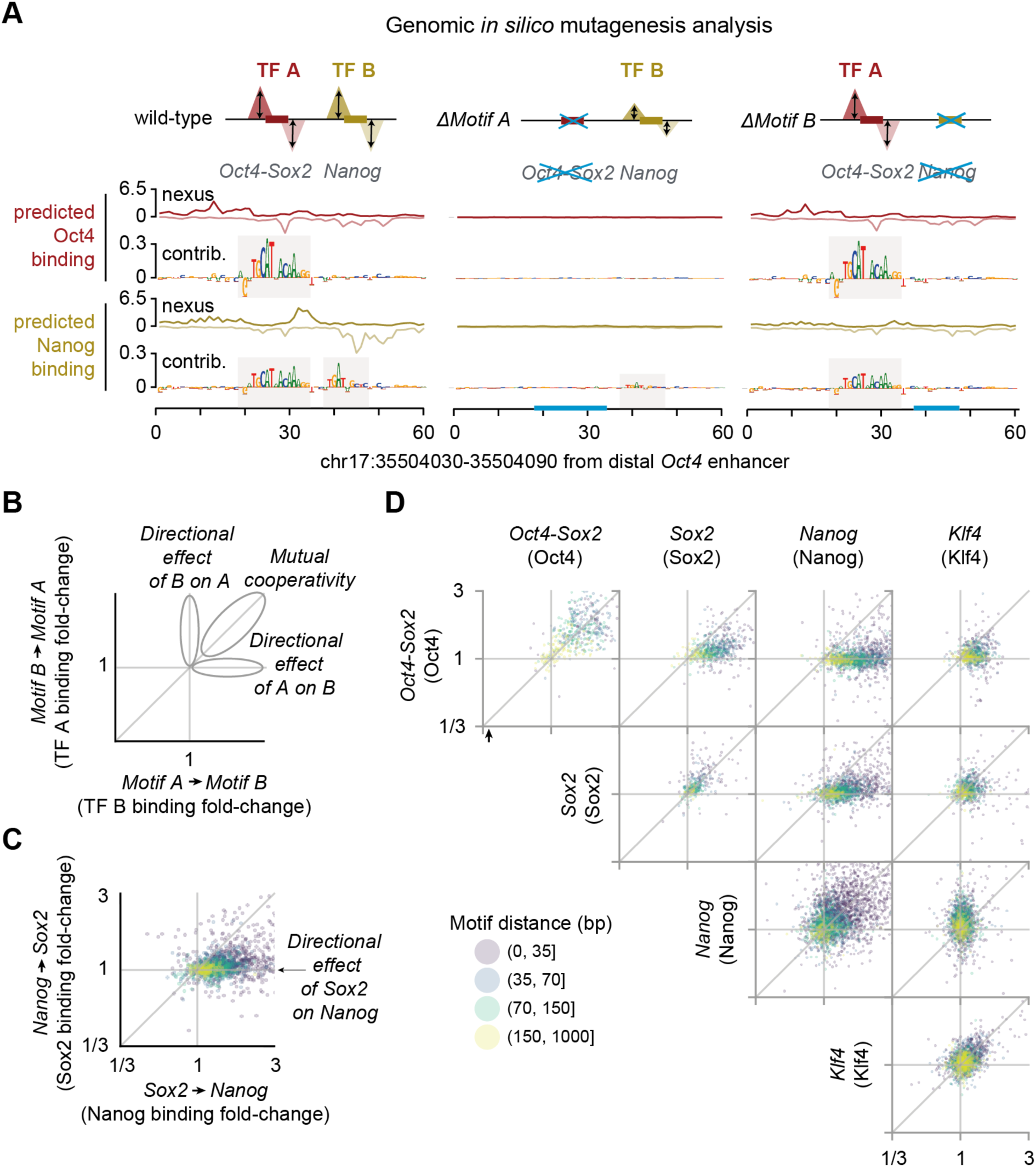
Additional information on the genomic *in silico* interaction analysis. **A)** Example genomic *in silico* mutagenesis analysis at the distal Oct4 enhancer. Predicted ChIP-nexus profiles and the contribution scores greatly decrease at both motifs (*Oct4-Sox2* and *Nanog*) when erasing the *Oct4-Sox2* motif (through random sequence insertion). By contrast, when the Nanog motif is erased (right), the predicted profile and the contribution scores of *Oct4-Sox2* motif remain intact. **B)** Such directional effect of motifs can be quantified by the corrected binding fold change (Figure S16A) for all motif pairs in the genome and visualized as a scatterplot. **C)** Example scatterplot for the interaction between Sox2 and Nanog. Sox2 shows a positive directional effect on Nanog most profound for short motif distances (<35 bp). **D)** Predicted binding fold changes for all motif pairs in genomic sequences.

**Figure S19.**
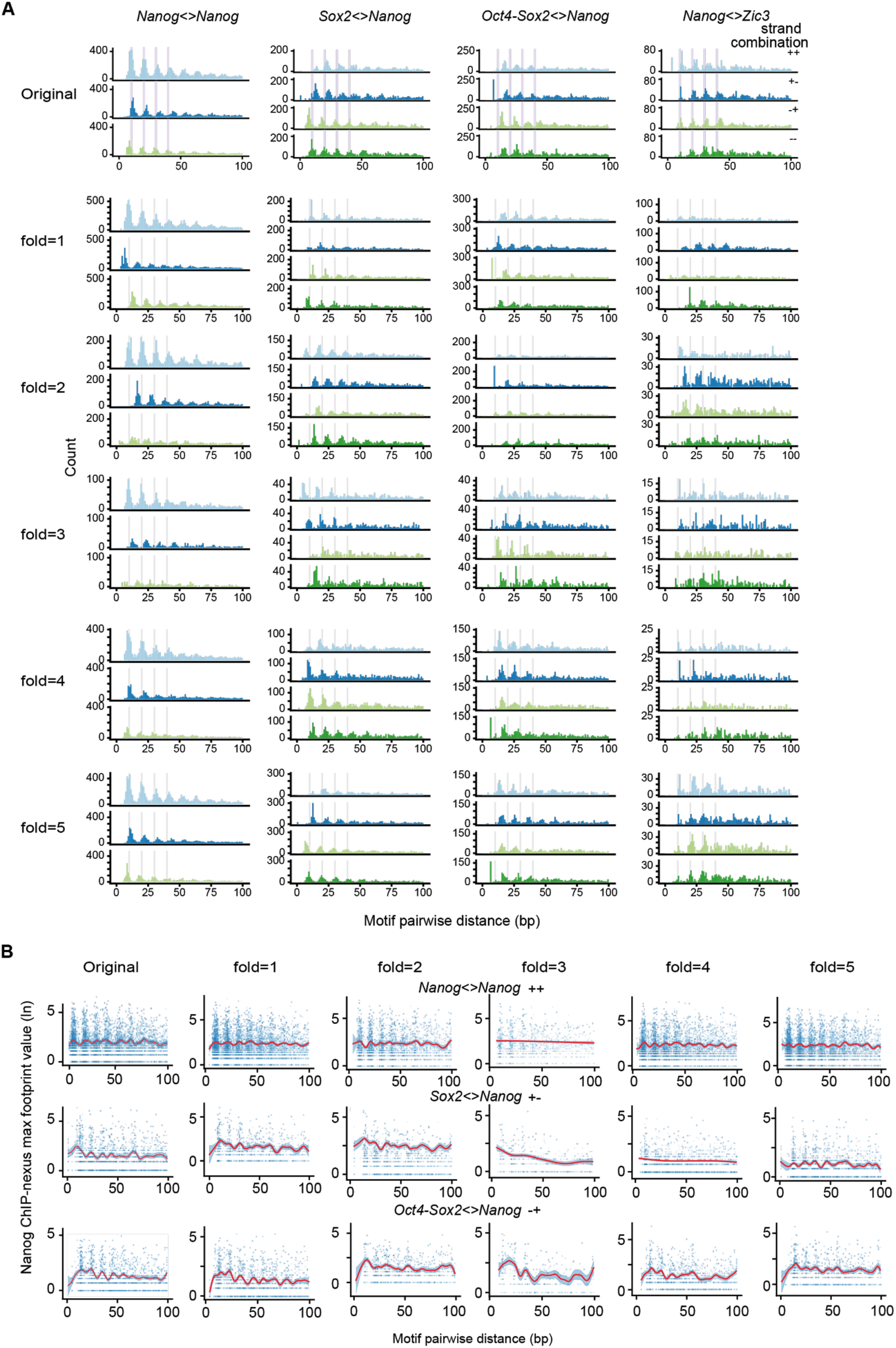
Motif syntax analysis for BPNet trained on different chromosome sets (folds). **A)** Distance distribution of motif pairs (as shown in Figures 5E-H) for motif instances obtained from BPNet models trained on each of the five different chromosome folds. The motif instances were obtained by scanning the contribution scores with the CWMs obtained from the different chromosome folds as shown in Figure S6. **B)** Nanog ChIP-nexus signal at the reference summit position for motif pairs as shown in Figure 5I-K for different BPNet models trained on different chromosome folds and the corresponding motifs.

**Figure S20:**
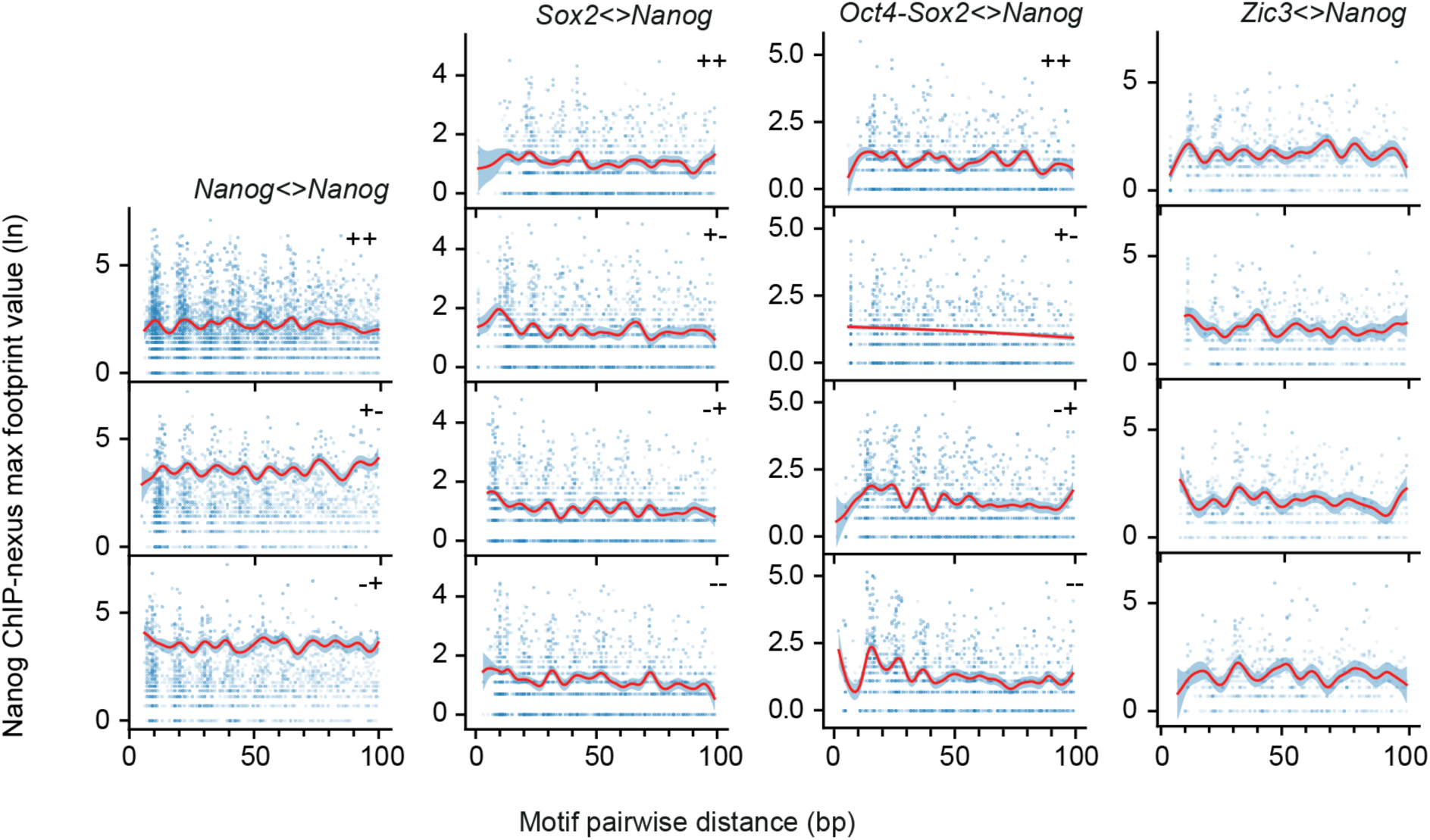
Periodicity of Nanog binding for different motif pairs. **Nanog exhibits helical periodicity independent of motif orientation.** Nanog ChIP-nexus binding maximum signal across *Nanog* motif pairs (blue dots), where the median of each motif pair distance is depicted as a red line. Nanog on average binds higher when the partner motifs (*Oct4-Sox2, Sox2, Zic3*) are within the preferred spacing to *Nanog.* This trend is observed for all motif orientations (unless there are not enough data points as observed for one of the *Oct4-Sox2<>Nanog* strand orientation).

**Figure S21.**
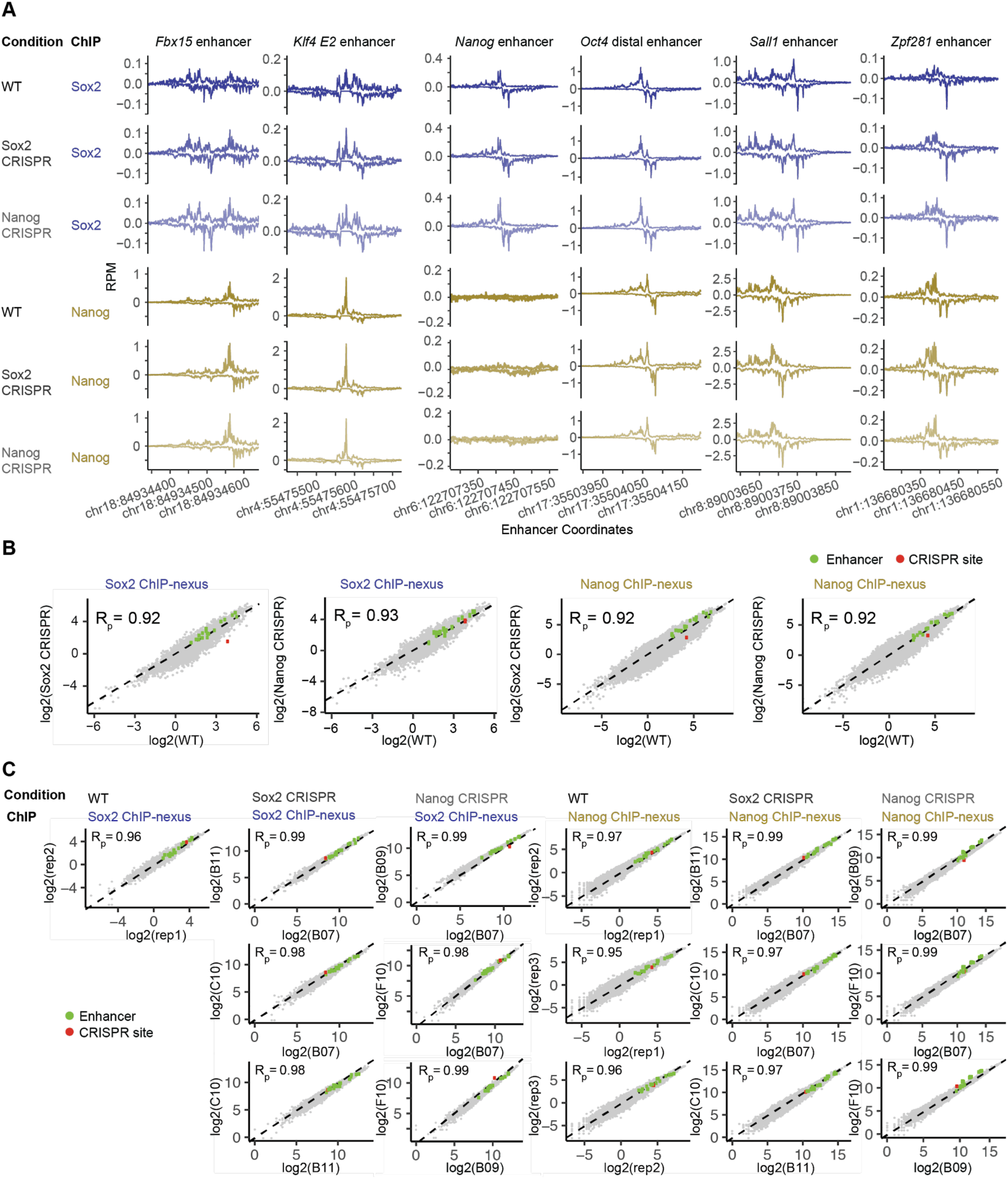
CRISPR validation experiments. **A)** Nanog and Sox2 ChIP-nexus profiles normalized to reads per million (RPM) show highly similar profiles and read counts across known enhancer regions for wild-type (WT) and CRISPR ESCs with either a mutated *Sox2* motif (Sox2 CRISPR) or mutated *Nanog* motif (Nanog CRISPR) at a selected genomic region (chr10: 85,539,626-85,539,777). **B)** Pairwise comparisons of ChIP-nexus RPM counts between WT and CRISPR ESCs at bound genomic regions (151 bp centered on the respective motif): Sox2 ChIP-nexus counts on *Sox2* motifs and Nanog ChIP-nexus counts on *Nanog* motifs (motifs based on the original model). The bulk data (gray) are highly correlated and known enhancer regions as shown in Figure S8 (green) are highly reproducible between ESC lines. Note the specific loss of counts in the selected mutated genomic region (red) over wild-type. Pearson correlations (R_p_) between groups are shown in the top left of each scatter plot. **C)** Pairwise comparisons of biological replicate experiments across WT and CRISPR ESCs show even higher reproducibility. Each point represents ChIP-nexus RPM counts across 151 bp genomic windows, centered on the respective motif. Known enhancer regions from Figure S8 (green) and the selected mutated genomic region (red) remain consistent between replicate sets. Pearson correlations (R_p_) between replicate pairs are shown in the top left of each scatter plot.

## Supplemental tables

**Table S1.** List of all ChIP-nexus and ChIP-seq replicate experiments and the associated quality-control metrics including the number of unique de-duplicated reads, highest number of IDR peaks between replicate pairs, number of “optimal IDR peaks”, the IDR rescue ratio, and fraction of reads in IDR optimal peaks (FRiP).

**Table S2.** Clustered motifs and their labels. Motifs were obtained by TF-MoDISco ran on BPNet models trained on 6 differnet datasets: i) seq/profile.peaks-union (ChIP-seq profile model trained on a union of ChIP-nexus/ChIP-seq peaks), ii) seq/binary (binary classification model trained on genome-wide ChIP-seq peaks), iii) seq/profile (ChIP-seq profile model trained in ChIP-nexus peaks), iv) nexus/profile.peaks-union (ChIP-nexus profile model trained on a union of ChIP-nexus/ChIP-seq peaks), v) nexus/binary (binary classification model trained on genome-wide ChIP-nexus peaks), vi) nexus/profile (ChIP-nexus profile model trained in ChIP-nexus peaks). Each motif logo shows the sequence information content of a PFM. The logo title consists of the manually assigned motif label (e.g. TE1, Oct4-Sox2) and the motif ID composed from the model name, the task name and TF-MoDISco motif ID (e.g. seq/profile/Nanog/m0_p13).

## Supplemental movies

**Movies S1-6.** BPNet profile predictions averaged across 128 random sequences with two motifs inserted at different positions. Centers of the motifs are marked by the vertical gray line. Motif distance is shown on the right. For each motif, the predicted profile of the corresponding TF is shown on the y-axis.

